# Mitochondrial Genome-Based Phylogeny of Turbellarians and Evidence for Accelerated Mitochondrial Evolution in Symbiotic Species

**DOI:** 10.1101/2025.07.01.662465

**Authors:** Jia-Qi Wang, Rui Song, Ming-Dian Liu, Tong Ye, Hong Zou, Gui-Tang Wang, Wen-Xiang Li, Ivan Jakovlić, Dong Zhang

## Abstract

**Background:** Flatworms are a highly diverse phylum with over 26,500 predominantly parasitic species. A minor portion of this diversity comprise predominantly free-living “turbellarians” Phylogenetic relationships within turbellarian orders remain debated, with recent mitochondrial genome studies also questioning the monophyly of the “Neoophora clade”. Some unique mitochondrial gene features have also been observed in this group. Within Turbellaria, the order Rhabdocoela includes significant symbiotic lineages, such as the endosymbiotic Umagillidae, Pterastericolidae, and Graffillidae, and the ectosymbiotic Temnocephalidae, which notably exhibits characteristics akin to a parasitic lifestyle. Given evidence linking parasitic lifestyles to accelerated mitogenomic evolution, we hypothesize that similar patterns: symbiotic turbellarians have mitochondrial genomes that accelerate evolution compared to free-living turbellarians. This study presents the first complete mitochondrial genome of the ectosymbiont *Craspedella pedum*, provides mitogenomic insights into turbellarian phylogeny, and addressing our hypothesis.

**Results:** The mitochondrial genome of *Craspedella pedum* is a circular DNA molecule of 18,456 base pairs, containing the standard 36 flatworm mitochondrial genes, a duplicated *trnT*, two *cox1* pseudogene fragments, a putative *atp8* gene, and several distinctive NCRs. Phylogenetic analyses based on 47 mitochondrial genomes, including the newly sequenced *C. pedum* and two assembled species from SRA database, using CAT-GTR model and BI and ML algorithms further confirmed that the paraphyly of the Neoophora clade and the basal position of Catenulida and Macrostomida. Different from previous study, Rhabdocoela forms a distinct clade within the turbellarians diverged immediately after Macrostomida before Polycladida and Tricladida. Within Rhabdocoela, *C. pedum* formed a direct sister clade with Typhloplanidae, suggesting a close phylogenetic relationship between the two. We also identified the paraphyletic of the Planoceridae of Polycladida and the unstable position of Planariidae within Tricladidain. Furthermore, we found a trend that the *nad4L-nad4* gene box in turbellarians may have evolved from an overlapping state, to the insertion of a non-coding region (NCR), and subsequently to a separated configuration, correlating with species divergence. The selection pressure analysis showcased selective relaxation from free-living species of Rhabdocoela to symbiotic species of Rhabdocoela. Furthermore, we also detected a relaxation from certain tubellarian lineages to Rhabdocoela. Along with higher GORR and longer Brl, Rhabdocoela and its symbiotic group possesses a more rapidly evolving mitochondrial genome.

**Conclusions:** The mitochondrial genome of *Craspedella pedum* displays uncommon characteristics, combined extended branch lengths and elevated GORR, suggesting rapid evolution and extensive rearrangements. Our phylogenetic analysis, integrating additional mitogenome data, further corroborates the paraphyly of the Neoophora clade and offers novel insights into turbellarian phylogeny. For the first time, we confirmed the accelerated evolution of mitochondrial genomes in symbiotic tubellarians compared to free-living ones, as evidenced by a longer average Brl, a higher average GORR, and relaxed selection pressure. Additionally, with the same evidences, we found that Rhabdocoela also exhibited accelerated mitochondrial genome evolution in planarians.

## Background

Platyhelminthes (flatworms) is a highly diverse phylum with more than 26,500 described species [1]. Its monophyly is strongly supported, and it is divided into two subphyla: Catenulida and Rhabditophora [2, 3]. The latter clade comprises a vast majority of flatworm species, including the strictly parasitic superclass Neodermata (the largest monophyletic radiation of parasitic animals); the remaining paraphyletic group of lineages is commonly referred to as “turbellaria” (it included Catenulida in some older classifications) [4]. While most of them are free-living, turbellarians also comprise multiple lineages with symbiotic lifestyles (both endosymbiotic and ectosymbiotic) [5]. They account for approximately one-quarter of described species in the phylum, and they are classified into 10 recognized orders: Macrostomida, Polycladida, Rhabdocoela, Tricladida, Prorhynchida (also called Lecithoepithelidata), Gnosonesmida, Proseriata, Fecampiida, Prolecithophora, and Bothrioplanida [6]. Most of these have ciliated epidermis [7] and have successfully radiated into almost all marine and continental aquatic habitats, as well as many humid terrestrial environments [8].

Molecular phylogenetic studies have relatively consistently confirmed the monophyly of these turbellarian orders [9-11], but the relationships among them remain debated [2, 3, 11-13]. A key feature in the reproductive biology of most Platyhelminthes is ectolecithality, a form of oogenesis where nearly yolkless oocytes and tightly bound yolk cells are deposited together in egg capsules [14]. However, three taxa within the phylum, Catenulida (class), Macrostomorpha, and Polycladida, maintain an endolecithal condition, where the yolk is internal [15]. This distinction led to the establishment of the “Neoophora clade”, comprising all ectolecithal lineages [11, 15]. Monophyly of the Neoophora clade first received support from analyses based on 18S and 28S rRNA and partial mitochondrial DNA genomes [11], but subsequent phylogenetic analyses based on transcriptomic data and nuclear protein-coding sequences indicated that Neoophora is a paraphyletic clade.

The mitochondrial genome has proven to be an useful tool for phylogenetic reconstruction [16] and has been widely applied in phylogenetic analyses of parasitic groups [12, 13, 17, 18]. The monophyly of the Neoophora clade also does not appear to be supported by recent complete mitochondrial genome-based phylogenetic analyses. One study produced a topology in which two ectolecithal groups Tricladida and Rhabdocoela, were sister groups, forming a clade that is closely related to the endolecithal group Polycladida, with the position of the endolecithal group Macrostomorpha varying across different topologies [12]. A subsequent study, which incorporated more mitochondrial genome data from Rhabdocoela, reconstructed Macrostomorpha as the “basal” turbellarian radiation, and Rhabdocoela, Tricladida, and Polycladida formed a polytomous clade [13]. Both studies also showed the relationships of some families within these orders.

As regards their mitochondrial genomes, turbellarians exhibit several unique architectural features, comprising an extra-long *cox2* gene in some species of Tricladida [19-21], and the existence of a highly divergent *atp8* gene, initially thought to be lost in the entire phylum Platyhelminthes, but later found in many turbellaria, albeit not in Neodermata [13, 22-25]. The overlapping element of *nad4L*-*nad4* gene box was also found exist in some turbellarian species [13]. Further, in contrast to the highly conserved architecture (i.e. gene order) across a vast majority of parasitic Neodermata, turbellarians exhibit rapidly-evolving gene orders [13, 24]. In an apparent contrast to this, parasitic Neodermata exhibit significantly faster sequence evolution rates than turbellarians [24].

Rhabdocoela is an order of turbellarias wherein some lineages constitute the major symbiotic groups within tubellarian, comprising the endosymbiotic families Umagillidae, Pterastericolidae, Graffillidae, and the ectosymbiotic family Temnocephalidae along with several other free-living families [26]. Temnocephalida, associated with economically important freshwater crustacean species, includes species that exhibit some characteristics of a parasitic lifestyle, including significant morphological changes such as the presence of suckers and tentacles [27-29]. Additionally, they feature a posterior adhesive disk and the absence of locomotory cilia, resembling characteristics found in certain neodermata parasite species. Furthermore, their epidermis, which contains multiple syncytial plates, is considered synapomorphic for the group [30]. The anterior tentacles and the singular posterior attachment organ cooperate to perform essential functions in both locomotion and attachment. Temnocephalida species employs a low, looping motion, alternately attaching the ventro-distal regions of the central three tentacles and the ventral surface of the posterior attachment organ [31]. In the light of recent evidence that the adoption of parasitic lifestyle (an in particular endoparasitism) is associated with highly elevated mitogenomic evolutionary rates [24], along with evolutionary relaxation of *atp8* and increased genome plasticity in endosymbiotic rhabdocoels [13]. We hypothesize that similar patterns: symbiotic turbellarians have mitochondrial genomes that accelerate evolution compared to free-living turbellarians. As there are currently (May, 2025) no sequenced mitogenomes of ectosymbiotic turbellarians, this hypothesis remains untested.

*Craspedella pedum* (Cannon and Sewell, 1995) is a temnocephalid ectosymbiont found in the branchial chamber of Australian crayfish *Cherax quadricarinatus* (von Martens, 1868) (Decapoda: Parastacidae) [32-34]. Preliminary studies have been conducted on the embryonic development [35] and sensory cells in the suckers [36] of this species. Furthermore, its symbiosis has been shown to cause white spot syndrome in its host [37]. Similarly, a recent study using a multi-omics approach reveals that *Temnocephala digitata* (Monticelli, 1902), a species within the same family, disrupts the nutrient metabolism and immune homeostasis of *Macrobrachium rosenbergii* (<u>Decapoda</u>: <u>Palaemonidae</u>) [28]. In order to test the above hypothesis, herein, we sequenced its mitochondrial genome (mitogenome), and based on the constructed phylogenetic tree, we compared three pieces of evidence: branch length (Brl), GORR (gene order rearrangement), and selection pressure. Additionally, with an expanded mitochondrial genome dataset, we aim to elucidate the phylogenetic relationships among turbellarian orders from the mitochondrial genomic perspective.

## Methods

### DNA amplification and sequencing

Partial sequences of NADH dehydrogenase subunit 4 (*nad4*), 12S ribosomal RNA (*12S*), and cytochrome c oxidase subunit 2 (*cox2*) were initially amplified by PCR using degenerate primer pairs. Based on the sequences of these fragments, specific primers were designed for subsequent PCR amplification. The PCR reaction was performed in a 20 μl mixture containing 7.4 μl of double-distilled water, 10 μl of 2× PCR buffer (including Mg2+ and dNTPs; Takara, Dalian, China), 0.6 μl of each primer, 0.4 μl of rTaq polymerase (250 U, Takara), and 1 μl of DNA template. Amplification was carried out under the following conditions: pre-denaturation at 98°C for 2 minutes, followed by 40 cycles of 98°C for 10 seconds, 48–60°C for 15 seconds, and 68°C for 1 minute per kilobase. The final extension was performed at 68°C for 10 minutes. The PCR products were then sequenced bi-directionally at Sangon Company (Shanghai, China) using the primer-walking strategy, as previously outlined. [38].

### Mitogenomic annotation and analyses

Global annotation of the mitochondrial genome was performed using the MITOS Web Server [39] with the RefSeq89 reference database [40] using genetic code 9 (echinoderm and flatworm mitochondrial). Transfer RNA genes (tRNAs) were annotated using both MITOS and ARWEN [41]. For tRNAs with uncertain automatic annotation, MAFFT version 7.149 [42] was used to align the anticodon regions with those of closely related species for confirmation. For the two ribosomal RNA genes (rRNAs), boundaries were determined by alignment with related species after MITOS annotation. Protein-coding genes (PCGs) were verified by comparing MITOS predictions with the Open Reading Frame Finder results [43] and closely related orthologues to identify start and stop codons. The sequence alignment of five NCRs was performed using MAFFT [42], integrated into Phylosuite [44, 45].

The *atp8* gene was not recognized by the automated annotation tools. Manual annotation was carried out by adapting the procedures described in [13, 22, 23], beginning with the translation of non-annotated ORFs into amino acid sequences using the Translate tool at ExPASy [46] and genetic code 9. The sequences were then analyzed using the ProtScale tool on the same website to generate hydropathy plots, and the SMART tool [47] was used to predict signal peptides and transmembrane regions.

The *cox1* pseudogene fragments (two sequences similar to the standard *cox1* gene) were aligned with the complete annotated *cox1* gene using MAFFT to determine the reliability of identification and location. Additionally, the sequences were subjected to a BLASTx search against a protein database, while also searching for conserved domains. PhyloSuite was used to generate the GenBank submission file.

### Comparative mitogenomic and phylogenetic analyses

Gene order and branch length of each species were also extracted using Phylosutie and Gene order visualized using iTOL. The gene order distance matrix of PCGs was calculated using the Common Interval measure in CREX [48], with the putative flatworm ground pattern of PCGs [22] used as the reference. Based on the gene order distance matrix and the tree-building results from the second round of the CAT-GTR model, we calculated the average values and standard deviations of gene order distances and branch lengths for free-living group (3 free-living species of Rhabdocoela) and symbiotic group (4 symbiotic species of Rhabdocoela). We also calculated for Rhabdocoela, Polycladida and Tricladida. To further verified whether there are significant differences of branch length and gene order rearrangement rate between symbiotic group and free-living group, we employed Permutation Test [49] in R. For comparisons between the orders, we performed the Kruskal-Wallis Test [50], which is more suitable for comparing multiple sample groups.

To investigate and compare the overlap region of *nad4L-nad4* box sequences within and between orders, as well as to trace the evolutionary trajectory of the *nad4L-nad4* box breakage in certain species, the *nad4L* and *nad4* gene sequences extracted by PhyloSuite were imported into MAFFT for alignment.

All genes (excluding *atp8*) from *C. pedum* and its sister species in the inferred topology - *Bothromesostoma personatum* (Schmidt, 1848) were extracted, aligned, and concatenated in Phylosuite. The resulting sequence data were then imported into DnaSP v5 [51] to calculate Nucleotide diversity (π) with a sliding window of 200 bp and a step size of 20 bp. Using counterpart FASTA format sequence of *C. pedum* as import, PhyloSuite was used to calculate relative synonymous codon usage (RSCU) for PCGs of *C. pedum*. Tandem Repeats Finder was used to identify repetitive sequences in non-coding regions [52]. MFOLD web server [53] was used to predict secondary structures of RNAs in non-coding regions.

45 available turbellarian mitogenomes were retrieved from the GenBank. A few species contained annotations labeled as “misc_feature” or had blank annotations, with sequence IDs starting with “MF” (see Table S1). We followed the annotation steps described above to reannotated these sequences. PhyloSuite was used to extract mitogenomic data, and conduct phylogenetic analyses using a range of plug-in programs. Following Monnens et al, two species of Gnathostomulida, *Gnathostomula armata* (Riedl, 1971) and *Gnathostomula paradoxa* (Ax, 1956), were chosen as outgroups [13]. Two datasets comprising aligned, trimmed, and concatenated sequences of mitochondrial genes were used for phylogenetic reconstruction: (1) nucleotide sequences of 36 mitochondrial genes (PCGsRNA dataset), and (2) the amino acid sequences of 12 PCGs (AA dataset). Both datasets excluded *atp8*. The extracted genes were imported into MAFFT [42], integrated into Phylosuite, for multiple sequence alignment. The alignment of protein-coding genes (PCGs) was optimized using MACSE [54], and Gblocks [55] was employed to trim the aligned sequences. For amino acid (AA) and RNA sequences, trimAl [56] was applied for alignment trimming. We used TreeSuite a plugin of PhyloSuite to calculate the substitution saturation (see Fig. S1-2) and signal-to-noise ratio for two datasets (PCGsRNA and AA) (see Table S2) and assess the suitability of the mitochondrial genome data for phylogenetic analysis. The trimmed alignments were then concatenated using PhyloSuite, and imported into ModelFinder [57] and PartitionFinder2 [58] to determine the optimal evolutionary models and partitioning strategy (see Table S3) for Maximum Likelihood (ML) and Bayesian inference (BI) phylogenetic analyses, carried out using IQ-TREE version 2.2.0 [59] and MrBayes 3.2.7 [60] respectively. We tested the performance of the CAT-GTR model in PhyloBayes [61] using the PCGsRNA dataset. Furthermore, in order to obtain the most stable phylogenetic tree and enhance the reliability of mitochondrial genome evolution rate analysis (as mentioned below), we assembled two mitogenomes (Accession Number: ERX6791043 and ERX6233253) of Rhabdocoela from all available sequencing data in the SRA database of NCBI using Mitoz 3.6 [62]. They were then combined with data from 47 (including 2 outgroups) previously obtained species for a subsequent round of CAT-GTR analysis. Phylogenetic trees and gene orders were visualized in iTOL [63] using PhyloSuite-generated annotation files.

### Selection pressure analyses

First we used KaKs_Calculator [64] to infer dN/dS values (dN/dS; non-synonymous/synonymous mutations respectively) between *C. pedum* and its sister species – *B. personatum*. Following this, we analyzed both trimmed and untrimmed PCG sequences, concatenating them for selection pressure analyses using RELAX [65]. The tree structure generated using the CAT-GTR model (incorporating two newly assembled mitogenomes) was used as the input tree. We conducted the following three rounds of analysis: 1) Marking the entire Rhabdocoela clade as foreground, with the remaining species as background; 2) Separately marking the free-living and symbiotic rhabdocoels as foreground, with the remaining species as background; 3) Marking the symbiotic rhabdocoels as foreground, with the free-living rhabdocoels as background. Since RELAX does not support manually marking background branches, in step three), we preserved the evolutionary relationships and branch lengths, and pruned the tree using MEGA [66] to retain only Rhabdocoela species. Additionally, we performed a reverse validation of step three by marking free-living rhabdocoels as the foreground and symbiotic rhabdocoels as the background.

## Results

### Mitochondrial genome characterization

The sequenced mitochondrial genome of *C. pedum* was a circular DNA molecule with a length of 18,456 bp. It contained the standard 36 mitochondrial flatworm genes, including 12 protein-coding genes (PCGs), 22 tRNAs, and two rRNAs. In addition, we identified a putative *atp8* gene, one duplicated *trnT*, and two *cox1* pseudogene fragments. The mitochondrial genome also features five major non-coding regions (NCRs 1-5). All genes are transcribed from the same strand, and all 12 PCGs used the conventional ATG or GTG start codon, and TAG or TAA stop codon. Exceptions were *atp6* and *cox3*, where we identified the abbreviated T--stop codon. The genome contained six gene overlaps, with overlap sizes ranging from 1 to 16 bp, and the largest overlap between *nad4L* and *nad4* genes (Fig. 1 and Table S4).

**Fig. 1.**
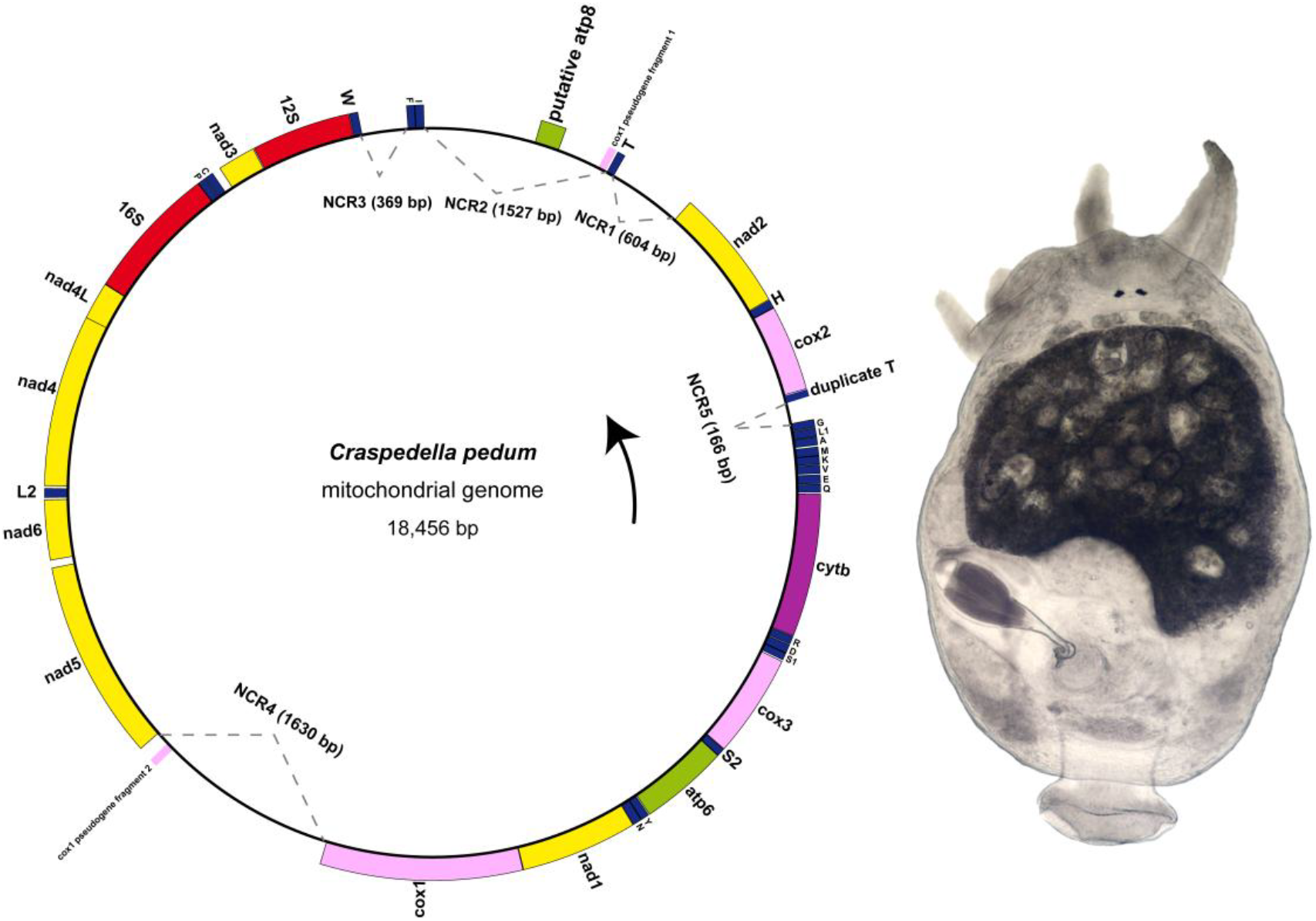
The map of the circular mitochondrial genome of *Craspedella pedum*, and its microscopic observation image. Arrows indicate the direction of transcription.

The overall A+T content of the genome was 71.2%, with AT-skew and GC-skew values of -0.228 and 0.283, respectively. The gene with the highest A+T content was *nad6* at 76.5%, while the lowest was *cox1* at 66%. The gene with the highest GC skew was *nad4L* (0.477), while the lowest was *atp6* (0.194) (Table S5). The most frequently used (474 times) codon was UUU (Phe), and the least used codon (0 times) was CGC (Arg) (Table S6).

*Craspedella pedum* had 23 tRNA genes, including one duplicated *trnT* (Fig. 1, Table S4 and Fig. S3). Most of the tRNAs could be folded into the conventional cloverleaf structure, except for *trnS1* and *trnS2*, which lacked the D loop (Fig. S3). *trnS1* uses the TCT anticodon, *trnS2* uses TGA, *trnL1* uses TAG, and *trnL2* uses TAA. The first position of the anticodon sequence for all tRNA genes is either C or U (Fig. S3 Table S4).

### Non-coding regions, *cox1* pseudogene fragments, putative *atp8*

Non-coding region sizes ranged from 1 bp to 1630 bp. Among these, there are five “major” (>100 bp) non-coding regions: NCR1 (604 bp), NCR2 (1527 bp), NCR3 (369 bp), NCR4 (1630 bp), NCR5 (166 bp) (Fig. 1). Some features that may be helpful for a better understanding of mitognoemic evolution and rearrangement mechanism in this lineage were identified in these NCRs: a direct repeat of 80 bp in NCR3 (Fig. S4); a putative *atp8* gene sequence in NCR2 (Fig. 1); NCR4 (261 bp downstream of the head) shares a similarity with NCR2 (61% similarity), and a cox1 pseudogene fragment was identified within the similar region. The tail of NCR1 (approximately 166 bp) shows high similarity (90% similarity) to NCR5. NCR3 and NCR4 did not align with any other NCRs (Fig. S5-7).

As regards the putative *atp8*, it comprised 202 bp, and a transmembrane region was identified between amino acid positions 10 and 32 in the putative protein (Table 1). The hydrophobicity patterns show some similarity to putative (or accurate annotation) *atp8* in some other turbellarians (containing two outgroups) (Fig. S8). Additionally, a signal peptide was predicted (40% likelihood) between amino acid positions 1 and 28 (Fig. S9), but the amino acid sequence does not start with “MNLC” (Table 1).

**Table 1.** The basic information of published (or putative) *atp8* in each species.

| Higher taxon | Species | Number of amino acids | First four amino acids | Transmembrane region |
| --- | --- | --- | --- | --- |
| Rhabdocoela | *Craspedella pedum* | 66 | MNLC | 10–32 |
| Macrostomophora | *Macrostomum lignano* | 55 | MPQL | 10–32 |
| Catenulida | *Stenostomum sthenum* | 43 | MYQM | 7–29 |
| Gnathostomulida | *Gnathostomula armata* | 52 | MKLP | 10–29 |
| Rhabdocoela | *Syndesmis echinorum* | 54 | MPQL | 7–29 |
| Rhabdocoela | *Syndesmis kurakaikina* | 47 | MPHF | 7–29 |
| Proseriata | *Nematoplana sp.* | 52 | MPHV | 7–29 |
| Gnathostomulida | *Gnathostomula paradoxa* | 54 | MPHM | 7–24 |
| Polycladida | *Prosthiostomum siphunculus* | 53 | MPQM | 7–29 |

### Gene order, branch length, and *nad4L-nad4* box

Rhabdocoela had a higher gene order rearrangement rate (GORR) and longer branch length (Brl) compared to other turbellarians. Additionally, within Rhabdocoela, the symbiotic species exhibited a higher GORR than the free-living species (Table 2). Our test results indicate that, with the exception of the GORR comparison between symbiotic and free-living species where the difference could not be definitively determined as non-random (p-value = 0.054), all other comparisons between data groups showed significant differences (p-value < 0.05) (Table S7).

**Table 2.** The average values and standard deviations of GORR and Brl for each group (with the number of samples for each group in parentheses). Table footnote. N.A., not applicable for groups with small sample sizes.

| Group | Brl | Standard deviation | GORR | Standard deviation |
| --- | --- | --- | --- | --- |
| Symbiotic rhabdocoels (4) | 8.808 | N.A. | 147.500 | N.A. |
| Free-living rhabdocoels (3) | 7.429 | N.A. | 134.667 | N.A. |
| Rhabdocoela (7) | 8.271 | 1.213 | 142.000 | 8.485 |
| Polycladida (13) | 6.076 | 0.088 | 141.467 | 6.830 |
| Tricladida (23) | 8.064 | 0.757 | 131.810 | 11.022 |

The *nad6-nad5* and *nad4L-nad4* gene boxes are conserved in *C. pedum, B. personatum*, and *Graffilla buccinicola* (Jameson, 1897), except *G. buccinicola* lacked the *nad4L-nad4* box. These gene boxes were also conserved in most other turbellarians (Fig. 2). In some species of the orders Rhabdocoela, Tricladida, and Polycladida, we observed that, in addition to at least one gene intervening between the *nad4L* and *nad4* genes, there is also the presence of a *nad4L-nad4* box where no genes are inserted between *nad4L* and *nad4*. This second phenomenon occurs in two patterns: (1) overlap between *nad4L* and *nad4* (16bp-50bp), and (2) the insertion of a non-coding region (NCR) of variable length (17bp-220bp) between *nad4L* and *nad4*. For species with overlapping sequences between *nad4L* and *nad4*, the overlap is 16bp in Rhabdocoela, 50bp in *Schmidtea mediterranea* of Tricladida, and 32bp in all other species of Tricladida. In Prolecithophora, the overlap is 32bp, 25bp in Proseriata, and 16bp in one species of Polycladida (Fig. 2). The overlapping sequences were highly conserved within each order and exhibit considerable similarity between the orders. The overlapping sequences of Polycladida align well with those of Rhabdocoela, and together, they match the first 16bp of the 32bp overlap in Tricladida. In each order, the overlapping sequences cannot align with the *nad4L* tails or *nad4* heads of non-overlapping species (Fig. S10).

**Fig. 2.**
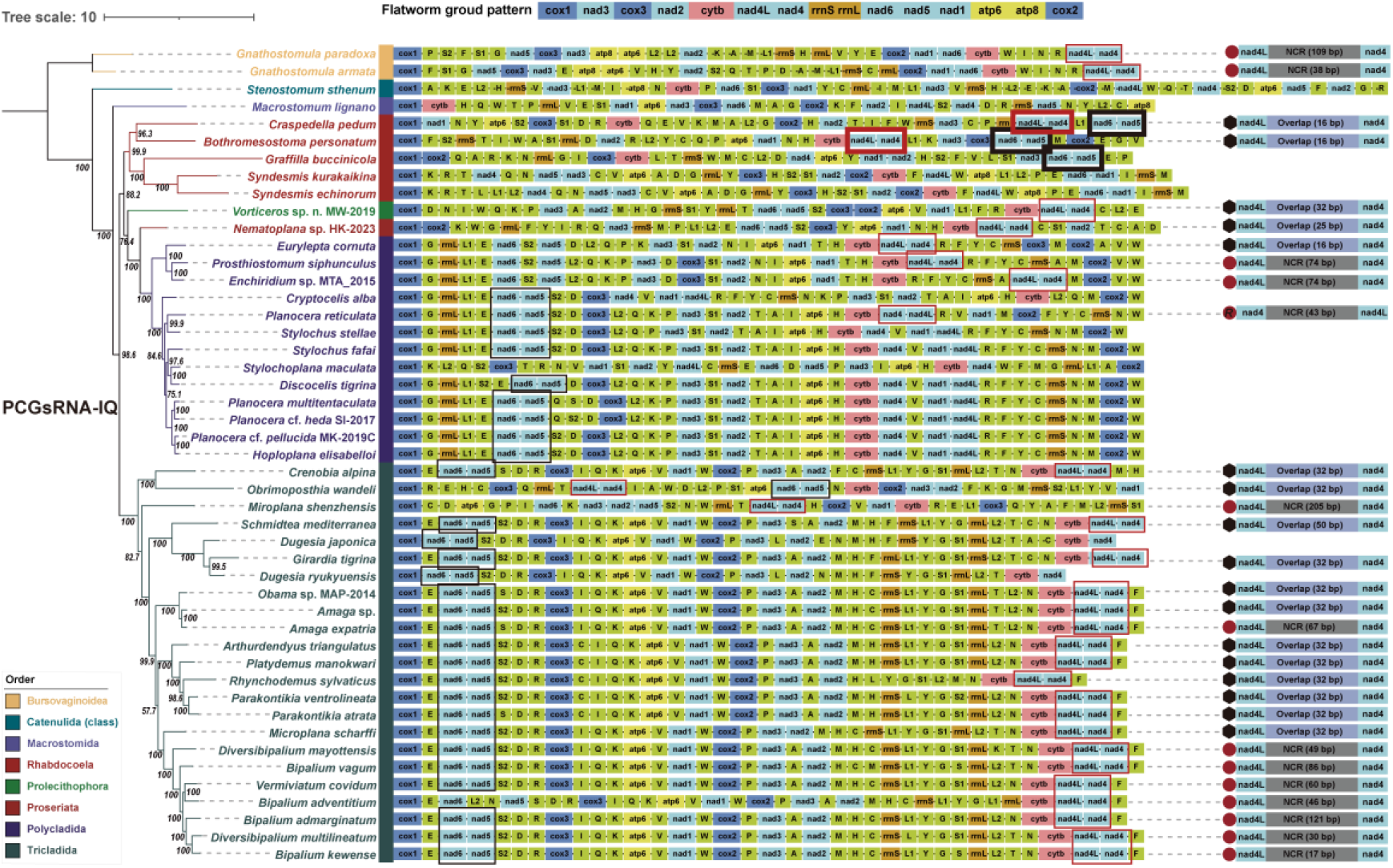
Phylogeny reconstructed using the PCGsRNA dataset and ML algorithms (PCGsRNA-IQ tree) with its gene order and the *nad4L-nad4* gene box shown on the right. The top represents the flatworm ground pattern. Red and black boxes in the figure highlight conserved gene boxes within turbellarians.

### Nucleotide diversity and dN/dS analysis

Genes exhibiting high nucleotide diversity included *nad4L* (0.547), *nad2* (0.467), *nad5* (0.435), *nad6* (0.427), *atp6* (0.419), and *nad4* (0.404), while *cox1* (0.328) and *cytb* (0.367) showed relatively low diversity (Fig. 3A) between *C. pedum* and *B. personatum*. The dN/dS ratio analysis against *B. personatum* indicated that *cox1* (0.056), *cytb* (0.085), and *nad1* (0.090) are evolving under strong purifying selection, whereas *nad4L* (0.287) is evolving under less stringent mutational constraints. Furthermore, all genes undergo purifying selection (Fig. 3B).

**Fig. 3.**
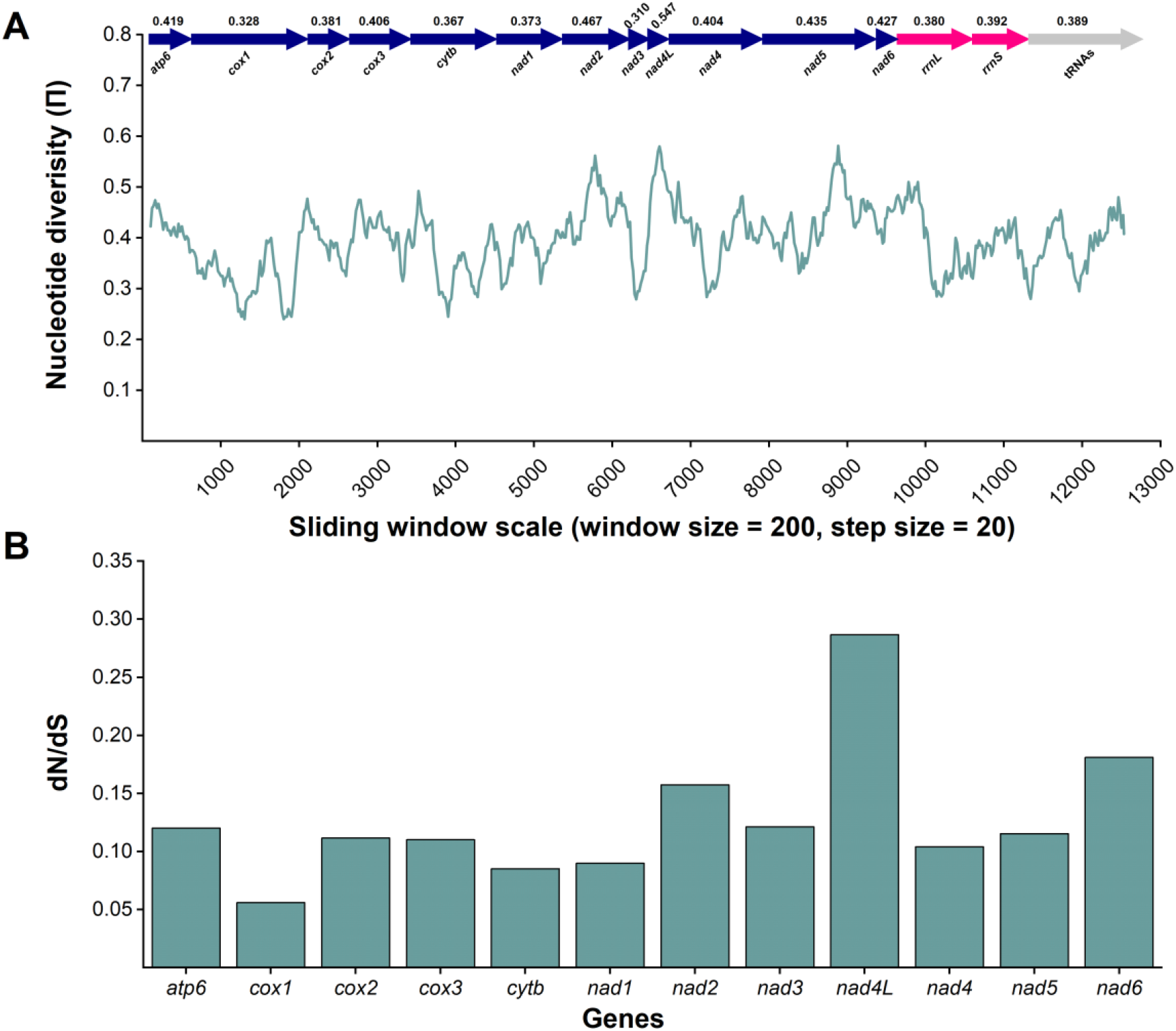
Sliding window and evolutionary rate analyses of the *Craspedella pedum* mitogenome. (A) Sliding window analysis of concatenated alignments of 12 PCGs, 2 rRNAs, and 22 tRNAs. The black line represents nucleotide diversity (window size = 200bp, step size = 20bp). Gene names, boundaries, and average nucleotide diversity are shown above. (B) Ratios of non-synonymous (dN) to synonymous (dS) substitution rates for protein-coding genes.

### Phylogenetic analyses

All orders were monophyletic in our analyses, and the inclusion of the Non-Neoophora clade Polycladida resulted in the paraphyly of the Neoophora clade (Fig. 2, 4 and Fig. S11-14).

**Fig. 4.**
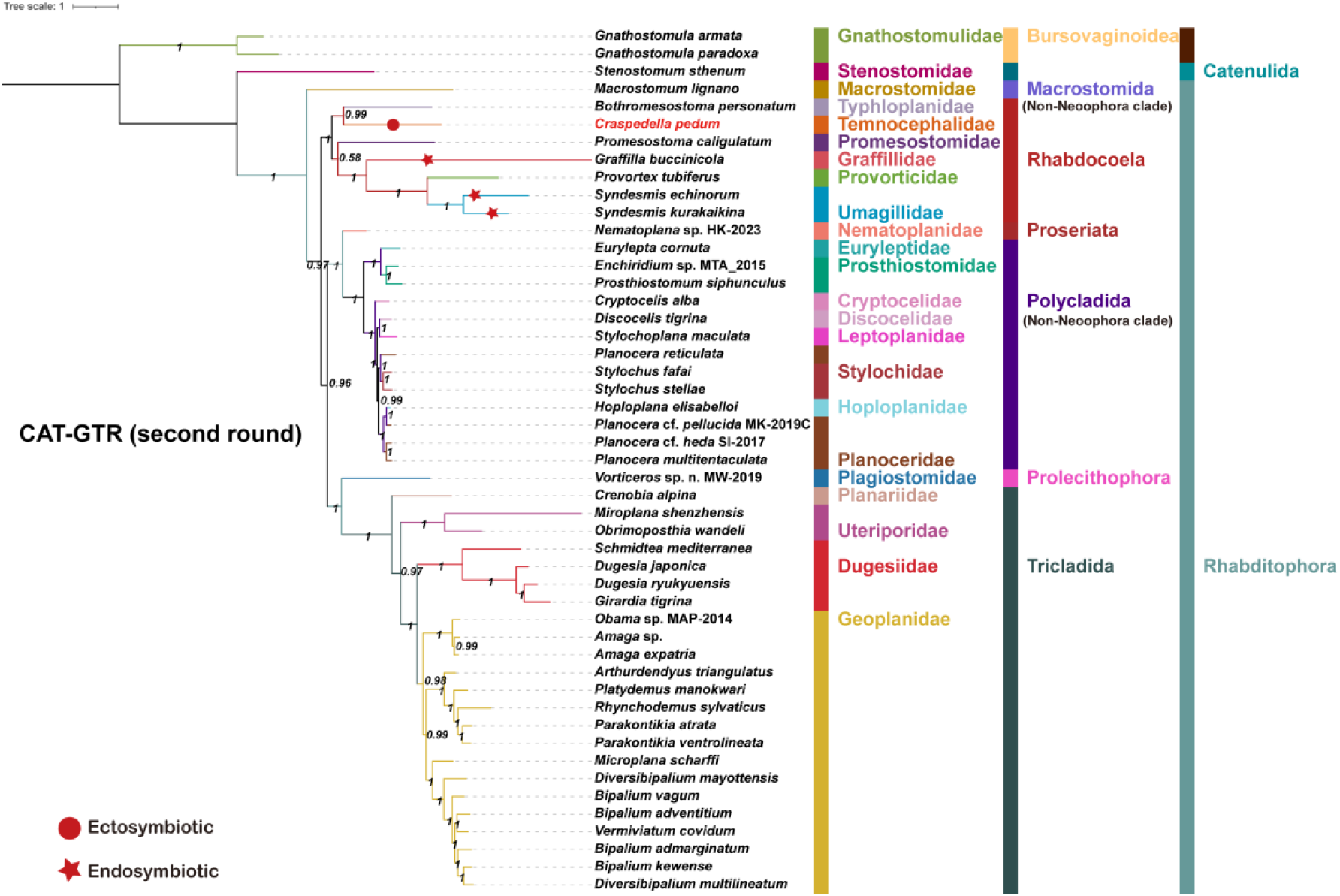
Second round phylogeny reconstructed using the PCGsRNA dataset and CAT-GTR model with two assembled data. Circles (ectosymbiotic) and pentagrams (endosymbiotic) denote symbiotic turbellarians, while unmarked individuals represent free-living.

Except for the topology produced by CAT-GTR model (both first and second round results), all other topologies were congruent: Catenulida was the sister clade to Rhabditophora. Within the Rhabditophora (turbellaria), Macrostomida was the earliest diverging (or basal) order, and the remaining species were divided into two major clades: 1) Tricladida and 2) a clade containing Rhabdocoela, Prolecithophora, Proseriata, and Polycladida. Within the latter clade, Rhabdocoela forms a sister branch to a subgroup consisting of Prolecithophora, Proseriata, and Polycladida, with these orders diverging sequentially. In the CAT-GTR model topology, Rhabdocoela diverged immediately after the earliest-diverging Macrostomida, Prolecithophora clustered with Tricladida, and Polycladida formed a direct sister clade with Tricladida rather than Rhabdocola with low support value (0.57) However, with the inclusion of the genomes from two Rhabdocoela species, this support value increased to 0.96 (Fig. 4).

Furthermore, there were also several differences observed at the family level within the orders Polycladida and Tricladida. Within the Polycladida, in the topological structures of AA-BI (Fig. S11), AA-IQ (Fig. S12), CAT-GTR (Fig. 4 and Fig. S13), and PCGsRNA-IQ (Fig. 2), Cryptocelidae diverges later than Euryleptidae + Prosthiostomidae clade, and earlier than Leptoplanidae + Discocelidae. However, in the PCGsRNA-BI (Fig. S14), its divergence occurred after both of these clades. The position of *Planocera reticulata* (Gray, 1860) varied across different topologies, as it did not cluster with closely related species, resulting in the paraphyly of Planoceridae In the AA-IQ, PCGsRNA-IQ, and CAT-GTR (1 and 2), *P. reticulata* forms a clade with Stylochidae. In the AA-BI analysis, it is placed within a branch that diverges later than Cryptocelidae but earlier than Stylochidae. In the PCGsRNA-BI analysis, it is placed in a clade that diverges earlier than the Euryleptidae + Prosthiostomidae clade, but later than the Leptoplanidae + Discocelidae clade. (Fig. 4-5, Fig. S6-S7). Additionally, Planoceridae and Hoploplanidae share the closest phylogenetic relationship because they clustered in all topologies. Within the Tricladida, in the AA-IQ, PCGsRNA-IQ, and PCGsRNA-BI analyses, Uteriporidae is consistently found to diverge earlier than Planariidae. However, in the AA-BI analysis, Uteriporidae clusters with Planariidae, and in the CAT-GTR model, it diverges later than Planariidae. Dugesiidae forms a sister clade with Geoplanidae, with Dugesiidae diverging earlier than Geoplanidae.

### Selection pressure

Selection pressure analysis indicated that, compared to the free-living species of Rhabdocoela, the symbiotic species experienced relaxed selection (k = 0.96). When the free-living rhabdocoels and the symbiotic rhabdocoels were each marked as the foreground branch, we observed that both groups underwent relaxed selection. Our reverse validation revealed an enhanced selection for free-living species, further supporting our hypothesis. Additionally, we also found that Rhabdocoela experienced selective relaxation in turbellarias (Table 3).

**Table 3.** Comparison of Selection Pressure Analysis Results

| Foreground | Background | Relax/intensification | p-value | K | LR | $\Delta AIC-c$ |
| --- | --- | --- | --- | --- | --- | --- |
| Rhabdocoela | rest of the tree | relaxation | 0 | 0.51 | 108.90 | 215.80 |
| symbiotic rhabdocoels | rest of the tree | relaxation | 0 | 0.98 | 150.41 | 298.82 |
| free-living rhabdocoels | rest of the tree | intensification | 0 | 0.63 | 112.67 | 223.34 |
| symbiotic rhabdocoels | free-living rhabdocoels | relaxation | 0 | 0.96 | 143.75 | 285.48 |
| free-living rhabdocoels | symbiotic rhabdocoels | intensification | 0 | 1.8 | 47.24 | 92.47 |

## Discussion

### NCRs and *cox1* pseudogene fragments

The complete mitochondrial genome of *C. pedum* exhibits multiple indications of frequent sequence duplications, comprising five large NCRs, within which we identified remnants of coding sequences, such as *cox1* fragments and a duplicated *trnT*. A direct repeat was also found in NCR3 may function as a control region contributing to the mitochondrial genome rearrangement. Our alignment of five NCRs showed signs of duplication among them. We speculate that the NCRs have undergone at least one complete and one incomplete duplication, which may be traces left by genome rearrangement. Within dataset, the mitochondrial genome of *C. pedum* is relatively large, though not the largest. The bigger mitochondrial genomes are found in several species of Geoplanidae (Tricladida), with the exceptionally long *cox1* gene and larger non-coding regions accounting for this observation [19-21]. Furthermore, a study found that the mitochondrial genome size in the sexual morph of *Schmidtea mediterranea* is larger compared to the asexual morph [67], and the sexual one with 27133 bp is the biggest mitogenome in used dataset. In some species of Neodermata, large mitochondrial genomes have been shown to be associated with genome structural rearrangements [68, 69]. Given those the genome of *C. pedum* is relatively large and exhibits a high rearrangement rate, along with evidence of non-coding region duplications and direct repeat elements in its mitochondrial genome, we hypothesize that the larger mitochondrial genome in this species may be related to mitochondrial genome rearrangements.

### Putative *atp8*

In turbellarians, *atp8* was first accurately annotated in the mitochondrial genomes of *S. sthenum* and *M. lignano* using transcriptomic methods, and its presence was inferred in several remaining polycladida species at the time [22]. Subsequent studies have identified *atp8* in the mitochondrial genomes of *S. echinorum* and *Nematoplana* sp.[13, 23]. The automated annotation software did not recognize *atp8* in *C. pedum*, and BLAST searches did not yield any significantly similar sequences. Following previously proposed workflows and standards [13, 22, 23], we found a putative *atp8* candidate based on the transmembrane region, hydrophobicity patterns, and signal peptide. A transmembrane region located between positions 10 and 32 of the amino acid sequence corresponds to such region in a putative *atp8* annotated on the basis of transcriptomic data in *M. lignano* [22]. The discovery of a signal peptide between positions 1 and 28 further supports the potential presence of *atp8*, a feature not observed in putative *atp8* of *S. echinorum* and *Nematoplana sp*., despite the 40% likelihood. The amino acid sequence does not begin with the common “MPQL” motif, found in other metazoans [70], but with “MNLC”. This unconventionality is also observed in other inferred species [13, 22, 23] as summarized in table1. It remains unknown whether this is a highly derived *atp8*, so transcriptomic data are needed to identify it with more confidence.

### Gene order and gene box

In our comparative genomics results, Rhabdocoela generally exhibited higher GORR and Brl values, a finding further confirmed by our Kruskal-Wallis Test. Those results support previous findings that mitochondrial architecture (gene order) is relatively rapidly evolving in turbellaria [24], and very rapidly in Rhabdocoela [13]. Between *C. pedum* and its closest relative in the available dataset, *B. personatum*, only two gene boxes were shared: *nad4L*-*nad4* and *nad6*-*nad5*, whereas the rest of the genes were completely rearranged between the two species. Furthermore, consistent with the observations of Monnens et al. [13], two endosymbiotic species from the family Umagillidae, *Syndesmis kurakaikina* and *Syndesmis echinorum*, lacked any conserved gene elements in comparison to other Rhabdocoela species, but their PCGs order are identical.

The *nad4L*-*nad4* box, considered an ancestral trait of bilateral animals [71, 72], has also been established as the ground pattern for Flatworms [22]. This gene box is considered to be functionally linked to bicistronic mRNA [73, 74]. In our phylogenetic dataset, the *nad4L*-*nad4* box is commonly observed, partly supporting the “ancestral trait” views.

Three arrangement patterns of *nad4L* and *nad4* in our dataset had been identified: 1) overlap between *nad4L* and *nad4*; 2) insertion of a non-coding region (NCR) between *nad4L* and *nad4*; 3) separation of the *nad4L*-*nad4* box, with the insertion of one or more genes. We found that the overlapping sequences are highly conserved within the order and share certain similarities between orders. Notably, the 16bp overlap in Rhabdocoela and Polycladida aligns well with the first 16bp of the 32bp overlap in Tricladida, suggesting that the *nad4L*-*nad4* overlap is a shared ancestral feature of them. Compared to Tricladida, Rhabdocoela and Polycladida are more closely related, as evidenced by all of our topologies, except for the CAT-GTR model.

The phylogenetic tree suggests an evolutionary track for the *nad4L*-*nad4* box in Polycladida. As species diverged, the *nad4L*-*nad4* box transitioned from overlap to the insertion of an NCR, and ultimately to complete separation. Since none species in Rhabdocoela exhibit an NCR insertion in the *nad4L*-*nad4* box and Tricladida does not show significant separation between *nad4L* and *nad4* in its most recently diverged species, we cannot make the same assumption for them, although they appear to follow a similar trend. For Macrostomida and Catenulida, traditionally considered basal taxa (also in our result), the *nad4L*-*nad4* box was separated. We speculate that the species included in our analysis may represent more recently diverged species within Macrostomida and Catenulida. The separation of the *nad4L*-*nad4* box does not challenge the basal position of these two orders. Our hypothesis is constrained by the available data, and we anticipate further data will be needed to validate.

Additionally, we sought to investigate the mechanism by which *nad4L* and *nad4* transition from overlap to separation across different orders. We compared the overlapping sequences from each order with *nad4L*-*nad4* box NCR insertion pattern. However, the results were the opposite, as the sequences aligned with internal regions of the genes. Three possible scenarios arose: 1) The overlapping sequence is retained at the tail of *nad4L* and the head of *nad4*; 2) It is retained only at the tail of *nad4L* or at the head of nad4; 3) This sequence is completely lost. Our results suggest that the overlapping sequence aligns with the internal regions of the genes, which does not support any of the three scenarios. This could be because overlapping sequences are expected to evolve more slowly than non-overlapping sequences due to stronger purifying selection constraints, as they must maintain the functionality of both genes. After the separation, however, the shift in selective pressures may have caused these sequences to mutate in different directions.

### Nucleotide diversity and dN/dS

The sliding window and evolutionary rate analyses yielded similar results. Both consistently showed that *cox1* is the slowest-evolving and least variable gene, and *nad4L* as the fastest-evolving. This is congruent with evolutionary patterns across many bilaterian lineages, as well as monogeneans [18, 75]. In addition, *nad4L* is recognized for its relatively short length and high variability in flatworms [76], and even putatively absent from some rhabdocoels [13].

### Mitochondrial phylogeny

Mitogenomes were susceptible in phylogenetic reconstruction because of compositional heterogeneity, substitutional saturation, and several other potential sources of error [77, 78]. To evaluate the suitability of mitochondrial genome data for phylogenetic analysis, we assessed two datasets (PCGs and AA). The AA dataset showed moderate substitution saturation with a linear regression (R^2^ = 0.814), while the PCGs dataset demonstrated a better fit (R^2^ = 0.871), indicating lower saturation. These results affirm the reliability for phylogenetic studies of the mitochondrial genome data used in this study. Additionally, we did not include the parasitic Neodermata lineage in the phylogenetic reconstruction. This decision was based on the fact that distantly related taxa with faster evolutionary rates can lead to long-branch attraction, which may result in incorrect inferences regarding the relationships among species [79].

The early morphology-based studies placed Macrostomida and Polycladida, along with Catenulida, among early branching lineages within the phylum Platyhelminthes [80], and they got support by some subsequent molecular evidence [2, 3, 11]. However, results inferred using CAT-GTR model of BI algorithms in Phylobayes and ML algorithms based on transcriptomic data [2] and hundreds of nuclear protein-coding genes [3] differ from this using Bayesian mixture model of Bayesian Inference (BI) and MCMC model based on 18S and 28S rRNA and partial mitochondrial DNA sequences [11] in positioning Macrostomida as an earlier-branching lineage than Polycladida. Our results support transcriptomic data and nuclear protein-coding data in this aspect, as well as in resolving Catenulida (class) as the basal radiation of Platyhelminthes.

The Rhabdocoela, Proseriata, Prolecithophora and Tricladida would be gathered into a single clade to the exclusion of the Polycladida, Macrostomida, and Catenulida in our topologies if the Neoophora clade was a monophyletic clade. In addition to earlier transcriptomic [2] and nuclear genomic [3] evidence, recent mitochondrial genome data-based analyses using ML, BI and CAT-GTR [12]and using ML and BI [13] have also raised questions regarding the monophyly of the Neoophora clade. The paraphyly of the Neoophora clade was also confirmed in our study by the insertion of Polycladida into the Neoophora clade. Unlike the clustering of Prorhynchida and Polycladida (Non-Neoophora clade) [2, 3], forming a sister clade with a branch contained Rhabdocoela and Tricladida [12], placing Rhabdocoela, Polycladida, and Tricladida in a trichotomous relationship causing the paraphyly of the Neoophora [13]. With more confidence from more mitochondrial genome data and different algorithms our all results except CAT-GTR placed Non-Neoophora clade, Polycladida, into a branch contained Prolecithophora, Proseriata. And this branch formed sister relationship with Rhabdocoela. Moreover, in this branch, Polycladida diverged lasted, Prolecithophora was resolved as clade that diverged earlier than Proseriata, a finding different from transcriptomic [2] and nuclear genomic [3] evidence, but partly supported by previous morphological analyses [80, 81]. In CAT-GTR, Polycladida (Non-Neoophora clade) diverged after Rhabdoceola, Prolecithophora as an earlier branch clustered with Tricladida which has also been observed in nuclear protein-coding genes evidence [3], 18S and 28S rRNA [82], and 18S and 28S rRNA along with partial mitochondrial DNA sequences evidence [11]. Despite high branch support, the analysis of Proseriata and Prolecithophora is limited by the availability of only one species from each order. However, for Rhabeocoela, with a low support valve (0.57) in main node of Polycladida and Tricladida, it diverged immediately after Macrostomida before Polycladida and Tricladida than diverged after Polycladida before Tricladida in transcriptomic data and nuclear protein-coding gene data [2, 3]. Subsequently, the addition of two mitogenomes of Rhabdocoela did not alter the first round CAT-GTR structure. Conversely, increased the support value to 0.97 for this major node, which further reinforcing the reliability of this divergence sequence. This finding is distinct from any previous studies. we believe our results are more reliable, as our sampling includes species representing the three different lifestyle types within the Rhabdocoela order. Nevertheless, this could also be an artifact resulting, at least in part, from uneven sampling across different orders. We also argue that in phylogenetic analyses of mitochondrial genomes, including species from diverse lifestyle types provides a more robust framework for interpreting the phylogeny of a given group.

Within Rhabdocoela, two main clades are identified, with one containing Promesostomidae, Graffillidae, Provorticidae and Umagillidae, and the other Temnocephalidae and Typhloplanidae. The relationship of Umagillidae Graffillidae and Typhloplanidae is consistent with earlier mitochondrial genome analysis results [13]. Temnocephalidae has previously been considered closely related to Typhloplanidae based on morphological evidence [83, 84] and 18S rDNA and partial 28S rDNA data using BI and ML algorithms [26]. The first sequnced mitogenome for Temnocephalidae, *C. pedum*, and our phylogenetic reconstructions, consistently support this relationship, as Temcephalidae and Typhloplanidae were sister clades with high support across all topologies.

For Polycladida, our analysis resolved Planoceridae as paraphyletic. In the AA-IQ, PCGsRNA-IQ, and CAT-GTR analyses, *P. reticulata* forms a clade with Stylochidae. In the AA-BI analysis, it is placed within a branch that diverges later than Cryptocelidae but earlier than Stylochidae. In the PCGsRNA-BI analysis, it is placed in a clade that diverges earlier than the Euryleptidae + Prosthiostomidae clade, but later than the Leptoplanidae + Discocelidae clade. The paraphyly has also been observed in a analysis based on rDNA data using ML and BI algorithms [85, 86] In contrast to rDNA data evidence, which clustered *Planocera multitentaculata* (Kato, 1944) with Stylochidae [85], our results placed *P. reticulata* within Stylochidae and consistently positioned *P. multitentaculata* within Planoceridae. Our results also support the monophyly and sister-relationship of Cotylea (Euryleptidae + Prosthiostomidae in our analysis) and Acotylea (all other families excluding Euryleptidae and Prosthiostomidae), consistent with the ML and BI analyses of mitochondrial genome data. In contrast, within Acotylea, Hoploplanidae clustered with the newly included Planoceridae than Stylochidae in our analysis [12].

In Tricladida, our results show that all families are monophyletic, and it indicated multiple independent evolutionary transitions from marine (Uteriporidae) to freshwater (Dugesiidae) and then to terrestrial (Geoplanidae) environments [82, 87]. Dugesiidae formed a sister clade with Geoplanidae which and is consistent with studies based on 12 mitochondrial PCGs using ML [88] and BI [89] algorithms. For Uteriporidae and Planariidae, in our AA-IQ, PCGsRNA-BI, and PCGsRNA-IQ tree, Uteriporidae diverges earlier than Planariidae and serves as the basal lineage of Tricladida, consistent with previous studies based on 12 PCGs data [89] and 18S rDNA and 18S rRNA using ML and NJ algorithms [90], also partly supported by the idea that the marine clade (Maricola, in our dataset is Uteriporidae) of *Tricladida* is likely the earliest diverging group, from which the freshwater (Paludicola, in our dataset are Planariidae and Dugesiidae) and terrestrial (Terricola in our dataset is Geoplanidae) clades have evolved [87]. In contrast, in our CAT-GTR tree, Planariidae diverges earlier and is considered the basal lineage. In AA-BI tree, Uteriporidae and Planariidae cluster together as the earliest diverging branch of Tricladida.

### Evidences for faster mitogenome evolution

Previous study had suggested that some endosymbiotic species exhibit relaxed selection in the *atp8* of the mitogenome, and this, along with their genome plasticity [13], provides indirect support for our hypothesis: that symbiotic tubellarias exhibit accelerated mitochondrial genome evolution compared to free-living tubellarias. Using mitochondrial whole-genome phylogenetic reconstruction as a backbone, we aimed to find evidence supporting our hypothesis through three main directions: GORR, Brl, and selection pressure. Our results show that symbiotic planarians have higher average Brl values and GORR, and that relaxed selection has occurred on the their mitogenomes. A similar association between parasitic adaptation and accelerated mitochondrial genome evolution has been observed in parasitic flatworms [24].

Beyond average values, we conducted a more precise comparison of GORR and Brl differences between free-living and symbiotic tubellarias using Permutation Test, which strengthened the evidence for Brl. However, the evidence for GORR was weakened in this test because p-value = 0.054. Despite a p-value of 0.054, its marginal nature prevents a definitive rejection of higher Brl in symbiotic turbellarians, suggesting that more data is needed for clarification. Additionally, through the Kruskal-Wallis Test, we compared GORR and Brl across different orders, and the results further enhanced the credibility of these two pieces of evidence.

## Conclusions

The mitochondrial genome of *Craspedella pedum* exhibits some uncommon characteristics Besides, with longer branch lengths and higher GORR, which suggest rapid evolution and extensive rearrangements of its mitochondrial genome. Our mitochondrial genome-based phylogenetic analysis, incorporating more mitogenome data, further confirms the paraphyly of the Neoophora clade and provides new insights that differ from previous studies on the phylogeny of tubellarias. Furthermore, we propose a trend that the *nad4L-nad4* gene box in turbellarians may have evolved from an overlapping state, through NCR insertion, to a separated configuration, correlating with species divergence. Our hypothesis that symbiotic tubellarians exhibit accelerated mitochondrial genome evolution compared to free-living tubellarians is supported by evidence from Brl, GORR, and selection pressure. However, the limited data available somewhat weakens the strength of the Brl evidence.

## Abbreviations

Brl: branch length
GORR: gene order rearrangement rate
*18S*: 18S ribosomal RNA
*nad4*: NADH dehydrogenase subunits 4
*12S*: 12S ribosomal RNA
*cox2*: cytochrome c oxidase subunits 2
ORFinder: Open Reading Frame Finder
PCGs: protein-coding genes
AA: amino acid
PCGsRNA: nucleotide sequence of 12 PCGs, 22 tRNAs, and two ribosomal RNA genes
AA: amino acid sequence of 12 protein-coding genes
ML: Maximum Likelihood
BI: Bayesian inference
NCR: non-coding region
tRNAs: transfer RNA genes
rRNAs: ribosomal RNA genes
*rrnS*: small ribosomal RNA
*rrnL*: large ribosomal RNA

## Declarations

### Ethics approval and consent to participate

All animal experiments were approved and conducted in compliance with the experimental practices and standards of the Ethics Committee of the College of Ecology, Lanzhou University (ethics approval form No. EAF2024012).

### Consent for publication

Not applicable.

### Availability of data and materials

The datasets for the conclusions of this paper are included in this paper and its supplementary material. The mitochondrial genome of *Craspedella pedum* has been deposited in GenBank (accession number: PV670564).

### Competing interests

The authors declare that they have no competing interest.

### Funding

This work was supported by the National Natural Science Foundation of China [32422089, 32360927]. The funding body had no role in the study design; in the collection, analysis and interpretation of data; in the writing of the report; and in the decision to submit the article for publication.

### Authors’ contributions

D.Z. and I.J. designed the study. and. conducted the experiments. J.Q.W. conducted the data analysis. J.Q.W. wrote the paper. All authors read and approved the final manuscript.

## Acknowledgement

The authors would like to thank Rong Chen of the BT-Lab (Wuhan, China) for helping with mitogenome sequencing and annotation.

## Authors’ information

J.Q.W. is a student in the Top Talent Program for the First-Class Discipline of Ecology (2022 Cohort) at the School of Ecology and Environment, Xizang University.

## Notes

### Competing Interest Statement

The authors have declared no competing interest.

