## Supplementary for "Mitochondrial Genome-Based Phylogeny of Turbellarians and Evidence for Accelerated Mitochondrial Evolution in Symbiotic Species": Fig. S4 direct repeat.pdf

### NCR3

AATTTTACTTTTACCTACTTACCCTTATTATAATGAAGAGGTACGAGAGATACCTCTATAG
TGGTGTGGGAATAACGAATATTGTGAAGAGGGAGAAGTTATAATTTGTGAGGTGTAAAT
TAAAATGGTTATTAATTTTCTTATTTATTTATCTTCTGTTTTATATACTATAATAATTAAGTG
TCTCTAAAATTCTTTTAAAATATCCTATGGTTTACCCTTATTATAATGAAGAGGTACGAGA
GATACCTCTATAGTGGTGTGGGAATAACGAATATTGTGAAGAGGGAGAAGTTATAATTT
GGGCCATCCTTTGTAAATATTTTTTGTGTATTTTTTTGTGCTTCCATTATACAGTTTTTTAA
TTAA

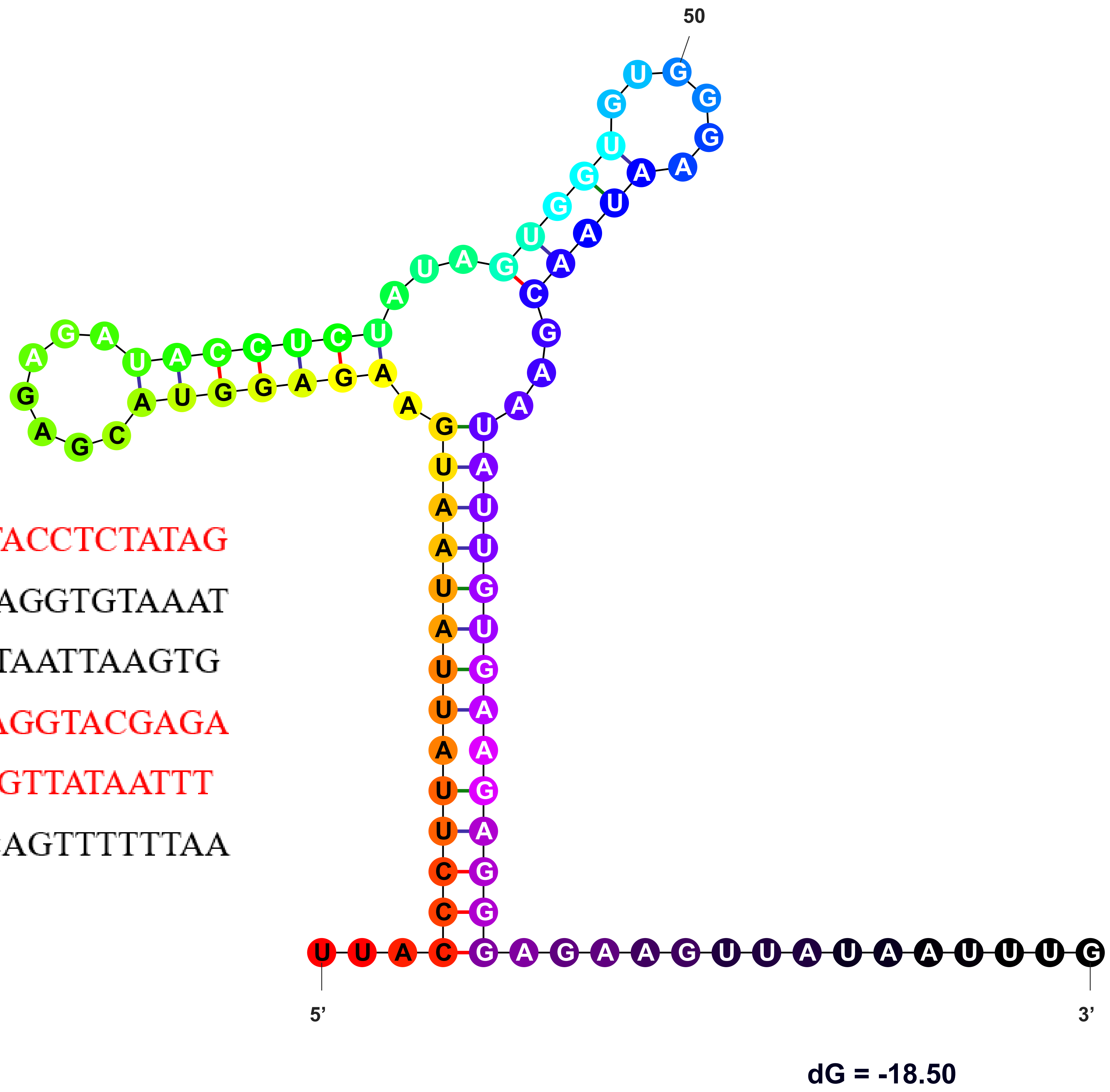
