## Supplementary for "Mitochondrial Genome-Based Phylogeny of Turbellarians and Evidence for Accelerated Mitochondrial Evolution in Symbiotic Species": Fig. S7 similarity_distance_matrix.pdf

| | NCR5 | NCR4 | NCR3 | NCR2 | NCR1 |
|---|---|---|---|---|---|
| NCR5 | | 0% | 75.000% | 72.000% | 90.361% |
| NCR4 | 0% | | 0% | 61.359% | 40.863% |
| NCR3 | 75.000% | 0% | | 0% | 75.000% |
| NCR2 | 72.000% | 61.359% | 0% | | 45.011% |
| NCR1 | 90.361% | 40.863% | 75.000% | 45.011% | |
