## Supplementary for "Mitochondrial Genome-Based Phylogeny of Turbellarians and Evidence for Accelerated Mitochondrial Evolution in Symbiotic Species": annotationBothromesostoma personatum.docx

ATTTTTTGTTTTAGTTAAAACCTTATACATAAAAATATTAGGACAATATATTTTACGGAAGTTTGATTTGGTGTCCTTGGAACATCATTATCTTTTCTTATTCGGTTAGAATTAGGTCAGCCTGGAACTTTACTTTCTGATGGTCATGTATATAAATGTTTAATTACTGCCCATGGTTTAATTATGATTTTCTTCTTTGTTATGCCTGTGTTAATTGGTGGATTTGGAAACTGGTTAATTCCAGTTCATTTAGGAATTCCAGATATGGCCCTTCCACGATTAAAAAATCTTAGATTTTGGTTATTACCTTCTTCAATGCTTTTAATACTGTTAAGTATGTTTTTAGGAGATGGTGTTGGTGCAGGTTGAACAATTTACCCCCCCTTAAGGAGTAATATTGCCCATGAAGGGATTAGAATGGATTTAGCTATATTCTCTTTACATATTGCAGGTTTAAGTTCAATTTTAGGTTCTTTAAATTTTAAAACCACAATAATAGTAAATTCTCTATTTTTTGGCTCTAAAGGTCGTTGAAGAGAAATTAGGTTATTTATTTGATCTATGTTTGTTACTGGATTTTTATTAATCTTTTCTCTTCCAGTATTAGCTGGTGGATTAACCATGCTTCTTACTGACCGAAATTTTAAAACTACGTTTTTTGACCCTATTGGAGGTGGTGATCCAATTCTCTTTCAGCATATATTTTGGTTTTTTGGACATCCTGAGGTATATGTTTTAATTCTACCAGGGTTTGGAATTATTTCTCATATTATGATAACTTATTCAGGAAAGTCTCAAACATTTGGGCATATTGCTATGGTCTTTGCTATTATTGGAATTGGATTACTTGGTTTCGTAGTTTGAGCTCATCATATGTTTACAGTGGGTCTAGACTTAGATACACGTGCTTATTTTACGAGGGCAACAATGATTATTGCTGTTCCTACTGGTATAAAGATATTTAATTGATTATTTACTGTGTATGGACGTCCTGTTTCTTTATTAAAGGATTATGTTGCACTTTTATGAGCTATTGGTTTCATATTTTTATTTACCTTAGGAGGAGTTACGGGTATAATCTTATCTAAAGCAAGGATTGATATAAGGCTTCATGATACGTACTACGTTGTAGCTCATTTTCATTATGTGCTATCTATGGGTGCTGTATTTTCTATTTTTGCAGGATTTATTCACTGGTGACCACTAGTTACAGGAAATGTTTTTAAATCAGACCTTTTATTTTCTCATTTCTGAATTATGTTCGTAGGGGTAAATTTAACCTTTTTTCCTCAACATTTTTTAGGTTTAAATGGAATGCCTCGGCGTTATATTGATTATCCTGATACATTTGGGTTTTGAAATCAAATTTCAAGTTTAGGATCATTAATGTCAATAATTGGGGTATTTTTCTTTTTCTATATTGTTTGGGAATCACTAATTTCTAATCGAGGCCTAGTTTATTTTTCTGTTTCTAAAACACAGATAGACTGAAGTAGTAATTTATTCCCACTTAGAAATCATACTTTTAGGGAAGTTAATGTTCGTTTTTT**GCTTA**AATAGCTTAAAAAAAGTGTAGTGTTGAAGCCGCTAAGTTGAAAACCCTCTTTAAGCAA**GGTTTTA**ATTACATATTATTTTGAAAATAATATAAGAAATCAAAAGTTTTCTAAAACCTTGATTCCTTGATTTTGGGAGTAAAAGTTCTATATACTTTCTACTATAGAAATCTCTTATAAAAATAGTTGATTATATTTTAAAATCATTTTCTAGTAGTTAAAATTTTTTTCTTATAAAATAAGAGAAAGGGGTATATATTTAGGTGATAAATCCAGGTGCCAGCAATCGCGGTTATACTTGAATACTTAATTTTTTTTTGGAACAAAGATTTAAATTATATTAAAATAAAAATTTAATCATATTGTTAAAAAAGTTTAATTTTTTATATATTTTAATTATTTAATTTTAATTTTATTTTAAAAAAAAAAGAAGATTAGAGACCTTCGTATTTTATAAAATATTTTTAATTTCACTTTAAGGGAAAAAATTTGGCGGTTTATAAAATTCCACCAGGGGAGTGTGAGGATTCATAAGATGGTCCACTAAAAAGTTTGCTAAATTTTTCTAGTAAGTGTATATCCGTGTTAAGGAGGTTTTGAAAAAATTTTTAAATTTCCAAAATTTAATATCTTAAAACAGGTCAATATGCAGCTTATAATTTAGGTTTCTTGCTCTACTAAATAAAACTTTTCTGAGAAGATCTTTTGGAATAATCTTACAGTAGAACTTGGAAGTAATTCTTGCTAGTTAGCAACAATGAAAGGGCTCTTTTTTTCGTACACACCGCCCGTCAATTTTTTTTTAAAAAAATAAGTCGTAACATGGTAACCTTAATAGAAATTTAGGTTAAAT**TCCTTTATAA**TATAATTGGGTATATTGATTTTGTAAGTCAAAAGAAAATGGTAAATTTTAGAGGAA**AATACAAGGA**CCGCATTAGTGGGCTACTCTGATATAGTAGACTGTAGAGGTATCTCTCTGTATT**AAGAATTTAGGTTA**TTAAGACCATTTATTTTCAAAGTAAAAAGAAAACATAAGTTTAATTCTTG**AACTTTATAGTTTA**TTAAAAACGTTTAATTTGCGTTTAAAAAAAAAAACCCTTTTAAAGTTAGAATTCGGAATTATTTAAGTATAGCTTCTAACTATTACAATGGTATTTTAACCTAGTTCCTAAAAGCTTATACTCCAAATTATTAGATATTTAATATTCTTTTTGTCTCACGAAATTTTTAAGATTTTATATATCTTAACCTTTCGAAATTTCTTTGATAAAAATTTCTTTTGTTTTGAACTAATTTATTGTAGTGAATTCTTAAAAAAAAAGAAATTTAGTTTTTGATAAATCTTCGTAGGGAATCTTCTAATTCAAAAATAAAATTTTCGAAAAATCATTTTTACTTTAAGGTAAAGATGACAACTTACTTATTATTATTTTATTATTTTTTGATTTTAATATTGAAATTCAAATAAAACTTTATTTTAAAAAACTAAAAAATTTTTAGTATTTATTATTTTTGTAAATCAAACTTTTAAAGGTTAATAATAAAATTATTAGTATAAAAAAAAACTAAAAGAGAACTCAATTTATATTTAACTGTTTTTCAAAAACATTTTTTCTTTATTAAGAAATATTCCCTGCTCCATGAAACTTAAATAGCCGCAGTATCTTGACTGTGCAAAGGTAGCATAATTGCTTGTCCTTTAAATAAGGTCTTGTCTGAATGGGAGATTAAAATATAATATTTTCTTTTAATTGAAATAGAAAATTTATTAAAGGTAAAAATACCTTTGTTTTTTAGAAAGACGAGAAGACCCTAAAATTTTTGATAAAAAAAATACTAATTATTTTAAATAGGGCATTTATTTTTTGTTTAAGAATTTTTTGAACTTAAAAAAATAGGTAAAATATTACTTTAGGGATAACAGGGTAATAAAAATTTGAAGTACATATTTGAATTTTTGATTACTACCTCGATGTTGAATTAGTAAAAAATCAAATTAAAGCAGAATTTTTTTATGAAAGCCTGTTCGGCTTTTAAATTACTACATGATTTGAGTTAAAACCGGTGCAAGCCAGGTTGGTTTCTATCTTCTTTTGATTATTTTTTGTACGAAAGGAACAAAATACTCTAAGCATT**TAAAGGTTAGTTTA**AATAAAAAATTTTTAATTGTCGGTTAAATAATGAAATTCTTTTCACCTTTAAAAAATTGTTCGAAGAAAATTAAAATTTTTTTATTGAGAGGATTCAAAGCTTTGGGTTTATATTTTATTTTTTTTATTTTGTAATTTT**ATGAAATTTTGTATTCTTTCTGCTGAG**AATATTAAAATTATGATTGTGTATTTACATTTAAATACATTTTTTTCTATACCCCTAATTTTTTTGTTTAAAAAAAATTGGAAATTACTAATCAACTATTTTCTTTTTCAAGTATTAGGAAGCATTTTCGTTGAAGTTGCATTTTGTTTTGATTATTTTATCTCTGATTATTTAGCCATTTTTGGTTTATTCTTAAAGCTTGGAATTTTACCTTTTTTTTGGTGATATCCATATTTTATAAAGGACAGAAATTGGTTTAATCTCTTTTTTTTTAATACTTTAAATAAGATTCCAGTTTTTTACCTATTAGGATACTGAGTACATGCTGAAGTTCTACTTATGGTAGTAGTTTTACTTTTTTCATTAATAGGGCTTTTTTTTTCAAGTTTATATATCTTTTGTTCAAAAAAAATTAAGTTTTTTTTAGGGTGAAGTTCACTTATTGATTCAGTTTTTTTTGTTTGATTGGCAAGTTATGATGATAAATCTTTTTTTTATTTATATATAATGTATTGTCTTATTTATTCTATTATTGCTTTTTGTTTTTTTCTTGCTTCTGAGCAAAAAACTACCTTAAAACCAATTTTATTAAAAACATCACGTCACCAATCAATTTTTTTTTTTCTAGCTTCCTTTTTTATTTTATTGTTAATTGGAATTCCACCTCTTTTTCCTTTTTTAATCAAGGTATTTAATATCTTTAGAGTTTTTGGAATAAAAACCTATTTTTTTATTTATTTTTTTTTTTTAGTATCACTAATACAAGGGTTACTTTACTTAAAGCTTTTTTTTTCTACTGCTAGTAGATTTAGGTTTAATTGGTCTTCATTAGTTTTTTCTAGCTTTGTTTTTTTTTTTTTGTTAATTTTTTTTTTCTGTCTTCCATTTTTCTTTTTAAAAAGAACCTAATTGTAGGTTACAGTTTCGACCTGTAGTATGAAATTTTATTCCTTTTTAAATTTAAAATGGCAGAAAAATGTGTCAGGTTTAAGCTCTGAAGATAAAGTTTACTTTTTTTTGTAAAAAAACTAAAAGTGTTGAACTGTAGATTCAAAAATGAAAAATTTTTCTTTTACAAATGTTTATTTTAAAAATTTTCCTTATTTCTTTTATTATGTTTATTATATTTAGAATTTCATTTTTTTTTTTACAGAGAAAAAACCTAAGTTCTGGCTTAGTTTCTCATAAACAGTTAAATTTAGTTTGGGCATTTTATTAAAAA**GACTAGATAGTATACA**AATTACAATTAATTGCAGATTAATAAATATCTTTTTAGGTTTTAGTTTTATAAAGAGGTGTTTTTGCATTCAAAATTTTGGTTTTTGAGGTGAAGTTTTCCTTTAT**TTAAAAGTAGTTTA**AATTAGAATAATAGTTTTGGGGACTTTTGGTCAAATATTTTGCTTTTAAAC**ATGAATTTAGATTTGTTTTC**TGCTTTAGGTTTAAAAAACTTAATATTTTTATTTCCTCTATTTTTCTTTACGTATTTTATATGACTAATCTTAGTTCTCCTTATTAAAACAAAAAATCGATTTTTTGTTTTAAATTCAAATATAATTTTTCTTTTCTTTTCTAATATTCTAAGGTCTATCTCTTACAAGATTCAATTTTCCGTTTATCTAATTAGCTTATTTTTTTTATTCTTACTTTACAGGAATTTATGAAAATTAATTCCTTTTACTATTAGCTTCTCTTCAAAAATAACATGAACACTTCTTCTTGGTTTATTTTTTTGATTTTTAATTAAATTTTCAAGTTTCTTCAAAAACATTTTTCAATATATCTCCCATTTTACTACACTAGGAAGTCCATTTATTTTAGTTCCCTTTATAAATATTATTGAGGTAATTAGCAATGTAATTCGACCATTAACTCTAGGTGTTCGGTTAGCAGTAAATTTACTCACAGGCCATTTACTTTTATCAATGTTCAGGAATTTTCATTCATCTTTATTATTTAGAAAATATTTTCTCTTTTTTTTTGTCTTTTTCTTTGGAATCTTTATTTTTTTTTATGAAAGATGTGTTTCCTTAGTACAAGCTTTTGTTTATGGGTTAATGATTAGTCAGTATTTTGATGAGCATTCCAATAATTAATTTTAAATGAGTTTTATTTCTGTTTGAATTGTAACAATTATTAGTGTTATTATTATGATACCATTTCTTACTTTAGTTGAACGAAAGGTGTTAAGATACATTCAGCTTCGAAAGGGACCTAAAAAGGTGGGTGTTTTAGGCGTTTTACAACCTATTTCTGATGGGGTAAAGTTAGTTTTTAAGGAAAGTGGTCCCACCATTCGCGTAAATTCATTTCTTTTTTGAATTACTCCACTTTTAAACTTTGTTTTAATGCTAATTTTATTTATAATCTTTTCTCCCTTATTTCCCTCTTATTCAATAAATTTAGGTTTATTAGGTTATTTATGTATATCTTCATTACTAGTATATTCAATTTTATTTTCAGGTTGGAGCAGAAAATCAAAGTACTCTTTCTTAGGGAGTTTACGTGGAGCGGCTCAGGTTATTTCTTATGAAATATCAATGTTAACTTTAATATTCTTCCCCAGTGCAATTTCTTCTACTTTTAATATCAATTTTATAAATATAGGTTTTCATTATTCAATATTAGTTTTTGTTTTTGTTTTTATTCTTTGGTTGATTACAATTGTTGCCGAAACTAATCGATCACCTTTTGATTTTGCTGAGGGTGAGAGTGAAATCGTTTCGGGATTTAAAACAGAGTATAGGAGATTTATTTTTGCCCTCTTATTTTTAGGTGAGTACGGAAAAATAATATTTATTAGGTTTTTAACAAGATTTATATTTTTTCCAAAATTTCCAGTCTTATATTTTATCATACCAATATTTATTATATTTTTATTTCTAGTTTTCCGCGGCACTTTCCCCCGATTTCGTTATGACCTATTAATGAATCTTGCCTGGAAGGTTATTCTTCCTTGCAGGTTAATTTTTTTTTGAATTTTATTTTTGTTTT**TGAAA**AATAGTGTTAAGCATATTAGGCCGTTAACCTAAAGGTGTATATTTTACTTTTTCAGGCCA**ATAGAAGAAGT**TTACAAAATATTAACTTGTGGTGTTAAAGAGGAAGAATATTCCTTCTATAATGAATAAGTTTTTAATTTTTCGGGTGGACAAAAAAATATTTTCTTCGGTAAAAAATCTCCTTATAGATTTACCTTGTCCTATAAATATCTCTTTTTTCTGAAATTTTGGTTCACTTTTAGGAATTACTTTTATTTTTCAACTAATTACTGGAATATTTCTTTCAATGCATTTTATCTCGGACGTAAACTTAGCGTTTGCATCTGTTGATCTTATTACACGAGAAATTTCAAATGGTTGAATTCTACGATTTTTGCATATTAGAGGTGCTTCTATGTTTTTTATCTTTATGTATTTTCATATAGGCCGAAACATTTATTTTTTTTCTTTTACTCTATGAAAGACCTGAAGAACTGGCGTTGTAATTTATTTTCTTTCAATGGCAATTGCTTTTTTAGGGTATGTTTTACCTTGAGGCCAAATGTCTTATTGAGGTGCTACTGTTATTACTAATTTTCTTTCGGCTGTTCCTTATCTTGGAAATGATCTAGTTGTATGAATATGAGGAGGATTTGCTGTTGATTACCCTACACTTACCCGATTTTTTTCTTTTCACTTTATATTTCCTTTTATATTATTAGTTTTTATGATTTTACATTTAATATTCCTTCACGAAAAGGGCTCTAATAATCCTCTAGGTTTAAAATCAAAAAATGACAAGGTTTTATTTTTTCCTTTATTTGCCTTAAAGGACATCTTTGGATTCTTCCTTTTATTTATTTTGTTTTTTTTTTTTTTTTTTTCTCCTAACAGCTTTTTAGAGTATCAAAATTTTTTAGAAGCTAATGCTTTAGTAACTCCTTTACATATTCAACCTGAATGGTATTTTTTACCACCTTATGCAGTATTACGGTGCATACCTAATAAGTTAGGTGGAGTTTTAGGTTTATTTTCTTTTATTTTTATTTTATTTTTACTTCCAATTTTAAAAAAAACAAAAATAAATAAAAAAACTTTCTTACGGAAAAGTTCTTATAATTTTATATTTCAATCTTTATTTTGAATTTGAATCATAAAATTTTTTTTATTAATGTGACTAGGTGCCTCTCCTGTTGAGTTTCCTTATCTTCAAATTTCTCGTGTATGTTCACTAATTTATTTTTTCTTTTTTTTTTGTTTTGCTTTTTTTTAATTTGGAATAGTGAATTTCTCTTTTTTTTTGTTTTTGGAATGATTATATTTAGAATGTTAACTAAAACCAAACGGTTTTTACATTTTTTAGTTTATCTAGAAGTTTTTGCTGTTTTAACTTTCTTTTTTTTACTGTTAGACGTAGCCTTGCAAGGCTATAGTGTATTATTAGTTTTATTAGTCATGATTTCATGTGAAAGTGCTATAGGTTTATCACTATTAGTAGTATTTGTGCGGAGCTTTAGAGTCGATAAAATTTCTGCTAAAAAAATTTCAACATGTGAGGGGTTTTAATAACTTTAATAGCTTTAAAAAATTTCTCTTTTTTCTTTTATTTCTTATTTTTTATTTTTTCTATCTACATAATAATACTTTTAGATTTTTCGTGGCCTAAAATCTTTTCATATTTTTTTTTAGCTGACAAACTCAAAATATTTTTAATATTTTTAACTATATTTGTCTTCTTTTTTATTTTATTTTTTACTTCTAATGAATATGTCTTAAAATTTTCTTTTTCAATTTTAACCATTTGCTTAATTTTATGTTTTTTAAGTTCAGAAATAATAATGTTTTATATTTTTTTTGAACTCTCTCTAATACCAACTCTTTATATTATATTTTCATGAGGCCTTCAACCTGAGCGAGTTAATGCTTCCTTATATTTTCTAATGTATGCTTTAATAGGTTCATTTCCTTTTTTAGCTAAAATTATTTATATTAGTTTATCTTTAGGGGAATTTAAATTTTTTATTCTAGGAAATTATAATTGTGACTATTTATATTTTTTTTGCAGAAAAAACTTGTTAATTTTTTTTTGAGTCTTTTCTTTCCTAATAAAGCTTCCTATTTATGGGGTTCATCTCTGGTTACCTAAGGCTCATGTAGAAGCACCTGTTTATGGTTCTATGGTTTTAGCTGCTATTTTATTAAAGTTAGGGGGTTATGGCTTAATTCGTTTAAGCATTTTTCCATTAATAAAAGTCAAAATCTTTTTTTTGTGAATAATTTTTAGTGTAATTTTATTAGGTTTTATTTCCTTTCGAAAAATTGACTTAAAGTCATTAATTGCTTATTCGTCAATTTGTCATATGGCTTTTTCAATTATTTTTTTTTATTTACTATTAAATATTAAAGTGTTTGGTGCATTAGTGATATTTGTAGGACATGGTTTTATTAGAAGGTGTCTATTCTTTAGATTCAATTTAATTTATTTAATTTCTAAATCACGAAAAATTTTTTTAAAAGAAGGTATAAAAAATATTTCTAAAATTTTTTATTTTTTTTTTTTCTTTATACTTTGTTTAAACTCATCACTCCCAATGAACTTAACTTTTTTTTCTGAAATTATCGTAGGTGTTAATCTTTTTTTAATTAATAAAATTTTTTTATTAGTTTTTCTTTTATCCGTTTTTTTAGTTGGTTTATTTAAAATGAAATTATTTTTATTTTCAAATCATGGGAATTCTTTAAAAAGATTAAATAATTCAATTTTATATTCACTTAATCTATTAAACCTTTCTATTATGTTCTTTCATATTATAATTAATTTTGTTTCGGTTTATTTTCTTTTTAGATGATTCTTTTAAATTTAAAAGTAGTATAAATAATACATTAGTTTTAGGGGCTAAAAATAAGGAATTCTTCTTTTAATAAAAGTAAAACATTTAGATGTGCCAGATTCTTACTCTGGAAAAGTAAATAATTTCTACTTTTAAATGTTTATTTTATTAGGGTCAGCTTTTAGTTTATTTTTTTTATTTTTGATTTTAGGAATTGGTTGAGTTTATTGAAATAAGGGTAAGAATTTTTTTAAAAATCGTGAAAAGAGAAGGCCTTTTGAGTGTGGGTTTGATCCAAACAATAATTCTCGTTTACCCTTTTCTTTGCGGTTTTTTTTACTCTTAGTTTTTTTTTTAATCTTTGATATTGAGGTAATTTTACTTATTGAGCTACCAGTGGTATTAAAAATCTTTTCTTTTAAAATTATTTTTGTAGTAAAATTTTTTCTAATTATAGTCTTCTTAGGATTAATAGAAGAATGGCGCCGAGGAATTTTAAATTGAAAGACATAATTTATATGATAAATAAGTTTCCATTCCATAAAGTTGAAAAGAGGCCATGACCAATATTTGTTTCTTTTATTCTATTTGGGTTTTTACTAAAAACTGTCTTTTTTTTTCATAATCTTGTTTCTTCATCTATAATTTTTATTTTTTTATTTTTATTAATTATAAGCTTATATTTCTGATGAAAGGACGTTATAAATGAGTCTAATTTAGGTTTTCATAAATATTTTATTATAAATAAATTTTATGTAGGTATGTTAATTATGATTTCTTCTGAAATTTTTTTTTTTCTAAGGTTTTTTTGGGCATTTTTTCATAATTGTTGATTTCCTGGTGTTGACCTTGGCAGTATCTGACCCCCTGCTGGGTTAGAGTTTTTAGTTGTAGACTCTTTTTCTGTTCCATTTTTAAAAACTATAATTTTATTATCTTCTGGAATATCAGTCACTTGAGCACATTATAGGTTAATTGCTGGAAAATCAAATAATTTACTTATAAGGTTAGTTTTAACCGTTCTCTTAGGTTTATTATTTTTATCTCTCCAAATTTTTGAATATTCTTCATGTCAATTTTCTTTTAAAAGTTTAGTATTTGGCAGATGTTTTTATATGCTTACAGGTTTTCATGGTGCTCATGTAATTGTAGGTACAATTTTTTTATTTATTTGTCTTCTTCGAGCTATTCTATTCCACTTAAAAAGCAACCACCACGTGGGCTTTGAGTTAGCTATATGATATTGACATTTTGTTGATGTAGTTTGATTATTTTTGTTTATTTTTGTTTATTGATATAGAAATATTTCTATTTAAAAGTGCTGGGAAGATTTGCGTTTATAATTTTTTTTTTTTTTTCTTTAAGGGGCCTAGCTACTATGGTATTCAAATTATTTGTCACAGCTTCTTTTATT**AGAATCTTCTG**TTTAATTTGAATTTCTAGTTGGCTTAGAATTGTTTTCTTTCTTATCTATGTAGGTGCTGTTTTAGTTTTAGTTTATTACATATTTTCCCTATCCTCAAAAGCAATTTATTCTTTTAATCCAAGAAGAGGAATTTTTTTTTTGTTACTCTTTTTTCCCACCAATCTTTGGTTAGAAGAAACCATGGTTGTAGAAAAAACTAATGATTGAGTTTTAGCCAGTTTTGATGGTTTTTTTTTTTTTTTTCAATTATATTATTAATTAGTCTTTGGGTTGCTTCAAAGTTGCAGCTTTTCAAATTTTTGGCTTTACGTGTATTTTTTTAAATT**ATGAC**AAGCTTTTATTTCTTATATTTTGTTTATATATTTATTTCTTTTTATTTATGTTCTTATTTAAGCAATTTTGAAAAATCAAATGTTTTAATTTATTTTTCTTTTTTGGATAAAACTAATTCATTGAGGTTTTTTGATTTCATTATTTCTTTAAATTGACTTAATATTCTCTTTTTGAGTGTTTTATTGTTAATTTCAGGATCAGTAGGGATATTCTCCATTACTTATATGGCTGAAGACCCAAATGCTAGACGATTTTTTTACATTTTAAACTTGTTTATACTCTCAATGATAGTTCTTATAATTGTACCCCATTATGTATTTTTTCTGTTAGGGTGGGATGGTCTGGGTTTTACTAGTTTTTTATTAATAAAGTATTATAATTCTAAGATTTCCTGATTTTCAAGTTTTAAGACTTTTCTTATTAATCGGTTAGGGGATGGTATATTATTAAGATCAATGGCTATTTATCTTTTTCAGGGCCATTTTTGATTTTTTTTTGAATTTTTTAATCACTATATACTATTATACTGTATTTTAGGTTTATTAACTAAGAGTGCACATGTGCCTTTTTCCAGGTGATTGCCTGCTGCTATGGCAGCGCCTACTCCAGTATCAGCTTTAGTTCATTCCTCTACTCTAGTTACTGCTGGTATTTACATTTTAATTCGTTTATCTTTTTTTTTTTCATCATTTTTACTATTATTAATTCTTTTTTTAGGAATATTAACAATTTTGTTAGGGGGCTTAAGTGCTTTAATCTCATTTGATAGCAAGAAGGTCGTGGCTTATTCGACTCTTAGAAATTTAGGATTTTTAGGAGTAAGTGTCGGATTAGGTTGTATTGGAGTTGCAATATTTCATTTATTTACGCATGGAATTAGAAAGGCTCTACTATTTATTTCAGTTGGAAATATGATGGAAACAACAAACCATAACCAAGATTTGCGCAATTTTTCTAAAATTTCTAAAAAAAATTTGTTCACTTTGGTCTGTCTTGTTTGGAGAATCTTGTCTTTATTAGGGGCTTTTTTTTTGTCTTGTTTTTATTCAAAGGATTATCTTTTTGAGGTTCAACAAGCTTCACCCAGGTTAAACATGATAATTTTTTGAGTTTTAAATATTACGGTTCTTATCACTTATCTTTATTCATTTCGTTTAATTTCTTTTTTTAGAAGAAGAAAGAATAAAATGTTTAATAAATTTTTCCCTTTCCAGTTAAAAATTTTTAAATTTTGTTCTAAGATTTTTTTATTTCTAGGGGTGCTTTTTTTTGGATTTTTTTTTCAGTATTTTTTCTTATTTGATTTTGAATCTTTTAGGATTTCTAAGATAATTTTTTATACATTATTTTTTTTTACTTTTGTAGTTCAAATTAAAAGCTTAGATTGAGTATCAGCATATCATTATTTTAATGGTGGGTTTTGGTTTGGTTTTGTCCATGTTGGTGATTTTATTTTTAATATGTTTGATTTAGGTTTATTTTCAAAAATAAACAGATCTATTCTAAAGGTATTATATCAATCATCCCAGTTACTAAAAAATTTTAGGTTTGGAAAATATGTTTTTTCTTTTTTTAATTTAGCTTTTATTTCAATTTTCATTTGGTATTTAATATTTTAAAT**TTAAAGGTA**AGTTAAAAAGACTACAGGACTCATGTTTCTGAAGTAGGAAATTCCTCTTTTAAAATCTAACTTTATTATTAAGTTTATGAAAAATTTTTTCTTTAAAAAAATACTTACATTAAATTTTCCACATGGGTCAACCCCTGTA**ATGGGA**AACATTATTAATTTTCATGAAAAGTGTTTACTAATTATATTGTTTGTCCTTGTGGTAGTTTTAACTCTTTTATTATATAAAATTCTTTTTTCACTAAGGCAGACTAATGGTTTGGAGAATACCTTATTAGAGATTTTGTGAACAGTAATTCCAACTTTAATTATAATTACAATTATGGTTCCCTCTTTAGAAATATTATATTTCTCAGAAGAGAGGGGAGGTTCTTTTTTTAGTAAAAACCTAAAGGTAGTAGGTCATCAATGATATTGAACATATGATGAATTTGAAATTTCGAATCAAGCTTTATTAAACTTTGACTCTTATCTGATTAAAGAATCAGATTTAAATTTAGGAGAATTTCGAAATAATGAAGTTGATCGTCCAGTAATTTTTGAGGTAAAACGTGCTACCCGATTGTTGATTACTTCAGCGGACGTAATTCATTCTTGGGCTGTACCTGAGTTTGGAGTAAAGGTGGATGCTGTACCGGGGCGATTGAATCAAATTGTAGTTTTTACAAGAGAGCCTGGGCGGTTTTTTGGTTATTGTTCTGAGTTATGTGGTGTAAATCATAGGTTTATGCCAATAGTTGTAGAGGTTATTAATATCTAGAAGAATTTGTTT**CTAAAACAAG**TGTTCTGCATATTAGATTTTCTCTCTAATGGTGAAATTTATTTTTCGTTTTAGAACTAAAATTTTTATAGTCAGAGTAAATACCACTGGGCGGCTAGGCAGAAGCTGATATATATATATATATATACGGGGAAGAAGATATCTAATAATATATTAAATAGGAAGTGAGAGTTCCCTCCCCTTCCCCCCTTTTCCGATTCCCAGATTTGAAGTCCTTTCAAAAGAAAGATAATTAGGCTTCGTTAAAAAAAAACACGAAAGCAAATTCTTTCATGCCCGGTTTCGCATTTTCTTAATTCTTTTATACAAAAAAAATCGTCTTTGGTTTCGAGATTATTTCCCCTACCCATCCCTGAAATTTTTTTTTTGGGAAAAAAACGAAAATTCCTGTTTCTGGGGGCAGAGATT**GCGACCCA**AGTTTGCTAAACACTTTCCTTCCACGAAAGAGTCTCCTTAGGTTAGGAGGGTTGTACTTTTCTT**ATACTTTTAGTTTAT**AAAAAATTAATTGTTTACACCAATTAGAAATCTCTTTAGGTAAAGTATATTCCTTTCTATCTAGATAGTAATTTTAATTAATCATTTGTTATATATAGGCACCCTGTGCCATTATTAAAGAATTCCTGTTTTACTTATAAATTATTGGGTTATAGATATTTAGAATAGGCAGGGTTATTAGTACAGTCTTTTTGAAGGAAAAAAAATATTTACATAATTTCCATGCTATTTATTTTCTCTATTGCTGCATAGTTTATTCATTGATGGCTATTAGTTAGAAGAAATGGTTTAATTTAAACATCTTGTTATTTATTTCTATTTTATTGTTTGTACGGTTTAAGTTTAAGCTTTTCAATTTTGTCCAATTTGTTTATGTGTCCCTTGCTCTTCAAATGATAGCTGGAATACCTCATCATATATAGCAATTTAAATGAAAATTCTATTTAGATGCTTCTTGTTTAGCTTTTTATCCTTATGAAAAACAATTATATTCATATTAAGGATCAAAGCTTTGCCAAGCTTTGCTCTATAAAAAATTAATTGTTTAAATGAATTAGAAAATTTTCTAGTCAGAGTAAATTAGGCGAGGAGCTAAAAATATAATATTTATACATAGGTATATCTAATTATAAATTTTATATAGGAAGGGGGAATTTTTCTAACATTTTCTTTGATATAAA
