## Supplementary for "Mitochondrial Genome-Based Phylogeny of Turbellarians and Evidence for Accelerated Mitochondrial Evolution in Symbiotic Species": annotationCryptocelis alba MF993331.docx

ATGACACCAGTCATAGGACTTGAATTTCGCGGTGACTATATTCTACTAAACATAAGGATATAGGAACTTTATATCTTATATTTGGTATATGAGCAGGCCTAGCAGGTACTGCATTTAGTTTTTTAATTCGTTCTGAGCTCGCACAACCCGGTAGAATTCTGAAAGACCCCCAACTTTATAATAGAATCATTACTGCCCATGGATTAGTA**ATGATCTTCTTCT**TTGTAATGCCAGTATTGATAGGGGGATTTGGGAAATGATTAATACCTTTATATTTGACAGCTCCAGATATGGCTTTTCCCCGATTAAAAAATATGAGTTTTTGGCTTTTGCCTCCTTCATTTTTTTTATTATTAGGGTCTTTCGTAGTAGAGTCTGGTGTGGGTACTGGTTGAACAATATATCCTCCTTTATCAGGTAAAATAGCGCATAGTGGGCCAAGAGTGGATATGGCTATTTTCTCATTACATTTAGCAGGAGTTAGATCAATATTAGGTTCTCTAAATTTTATTACAACAATGGTAAATGCTAAGTTGCAGATGTTATGAAGAAACTTTCCTCTATTTTTATGAGCAATAATCATAACGGCCTATATGTTGGTATTATCCTTGCCTGTACTGGCAGGGGGCTTAACTATGCTTTTAACTGACCGTAAATTTAATACCACGTTTTTTGACCCAGGAGGGGGGGGAGACCCCATCTTATTTCAACATATATTCTGATTCTTTGGGCATCCAGAGGTATATATCTTAATATTACCTGGCTTTGGGATGGTGTCCCAAGTCGTGACCTTTTATAGAGGAAAGGATAGAGCCTTTGGTCATATGGGAATGGTGTATGCAATTTTGGGAATAGGACTTCTAGGATTCATAGTATGAGCACATCACATGTATACTGTAGGATTAGACATAGATACTCGTGCCTATTTTACTGGAGCCACTATGATTATTGCTGTACCTACAGGTATAAAGATATTTAGATGGTTGGCAACTTTTTATGGCCGTCCTTTACGAGAGTTAATAGATAACACAGGACCAATGTGGGCTACAGGCTTCATTTTTCTATTCACTTTAGGGGGCTTAACCGGGGTAGTTCTAGCTAGAGCAAGTTTAGATATAAGCCTACATGATACTTATTATGTGGTAGCTCACTTTCATTATGTGCTCTCTATGGGAGCCGTTTTTTCTATTTTTGCTGGGATAGTTCATTGATGACCATTGTTCGTTGGAACAGGCTTAGATTCTAAGATGTCAATGGCCCAATTCTGAATACTGTTTACAGGGGTTAATTTAACTTTTTTCCCCCAACACTTTTTAGGTCTTGGGGGTATGCCTCGGCGTTATACGGATTATCCTGACGGGTTTGGTTATTGAAATGGTATTTCTTCTTTCGGTTCCTTAATTTCTGTAATAGGAGTAGTTGCTTTCTTAGTAATTATATGAGAATCAGTGTTAAGTGAACGGGTGTTATTATATGTTGTTTCTCCCAATTCACAATCCGAATGGACTGGGTCTTGGTATTTTCCTTTCTCTTTCCATACAGGAGATTCAGTAACAGTAATGTATCTTGAATAAATCACTTCTTTTTAGTTTAAAAAGAGAATGTTTGCCTTCCAAGCAAAAGGTCCTAAAAAGGAAAAGAAATACATTATAATTTACGTTTTGGTTCTCGAAAATTAACTGGTAAAAGGTAAATTATTAATAATATTGTGTAGAGATAATTACCTTTTGAATCATGTTCTAATAAATACTATTAGTGAGGCTATATTT**CCGAAACTCTAGGATTATAAGTTCTT**TAGTTCAGAATTTAAATATTAAAAGGGCATTTATATTAAAAAAGATCTTAAGTACAGAAATTGTAAATCGGCTAGAGTCATATCTGATTAAAAGTACTGAATTTCATGATTCCTTTCCCACGTAAATCGTGGGAAATGTAGTTTTAGCTTAGAGTATTAATAAATTCATTGTTTAAGATGAGAGGTGTCTAAAATTAATTGAGTGTTATAACTTTAGTTAATCAAAATAATGCTAAATATATTAACTTTGAAATCAACTTATAATTTATACTGATAGAATTAGTATACTAAATAAAACCATGGGAACTCGATATTTTTACAACTGTTTATCAAAAACATCTCCTTTAGTAAATAATTAAAGGTAAGACCTGCTCAATGTTATGTATTAATAGCTGCGGTATCTTAACTGCACAAAGGTAGCATAATTAATTGTCTTTTAAATGGTGACGGGTTTGAAAGGTTTAATGAGAGAAATAATATCCTTGGTTTAATTATAAATTATATTTAGAGGTTAAAATACCCTTTTATATAAGAAAGACGAGAAGACCCTGTTGAGTTTTACAAATATATAAATATTTGTTTTGATGGGAAATTTGGATTCTAATTGAATACTTTGTTGATCCTGGAAATAATTTCTGGAAAAAGGATTAAATTACCGCAGGGATAACAGGACTATATTTACTGAGAGTTCTTATTGAAGTAAATGATTGTTACCTCGATGTTGAATCGAAAATTACTCACCGACGCAGCAGTTGGCTAAGAGTAGGTCTGTTCGACCTGTAGATTTTCACGTGATTTGAGTTAAGACCGGCGTGAGCCAGGTTGGTTTCTATCTTCTTTTGTTTTTTTACTGTACGAAAGGAACTAAAAAATTGGGATGACCAAATATTCAAAATGGCAGATAATATGCATTAGATTTAGGTTCTAGAGATGGGGAGTTTCCCCTTTTGAAATACTAGTATAAGTGTAATTTAGCACATTAAGTTTTCTCCTTAAAAGGGTTCTAATTAAGAACATATTAGAAAATAGATAA**ATGTCCTCTTTTTTATGGTTTGTCTTGATAGTCGCAAAATTTCTTTCGTTATTTTTTATTAATACTTTAGGATTGGCCT**TTTATATTTTTTTAGCTAGGGTATCTATTTGTTTACTGATGGCTTTCCATGGGAGCATATGATATGCTTTAATATTCTTTCTTATATATGTAGGAGGAGTATTAGTACTCTTTATTTACATATCCTCATTAAACTTCAAACCCGTCTTTCGAAATCAATCCATTTGAACAGGATGGGTAGGAAAAACAATCAAGTTTCAGCTTTTCTTTATTCTTTTTAGGGTTCTATTATTTAACCCTCATGAGGGTAAGGGAACAAAATGAATAGCGTTAACGTCTAAGGAGTATAGATATGATCTATTTCTTCAAACGGAAACTTTATTTTTAGTTAGTGTGGGGGCATTATTGTTATTTGTTCTGTGGGCAATAAGAAAGTTAACCTTTCGAACGCGAGCTAGACTTCGCCCCTTTTTTGATAATTTATAATTATATAAACCTAATCCCAAAGCTTAGAAGATCATTGGCTCTCCCAGAATTACGG**ATGGACTTAGTTAAGACTA**AATGTAGATTAGCAATAATCAGCTCCAAAATATTCTTCTTCCTTTCTATACTCTTTGGGGTCTTATATAGCTTTAATACAAAAAGTTATTTAATTACTCTAAAAATCTTAGATTTTTGTGGATTAAGAATAGATCTTAGGTTCTTTGTGGACTGGATTAGGCTGTTATTTTTTAGTGTGCTGTGCGTCATCGTGTCATGTGTATTGAAGTTTTCCGAAATTTACATGGCCAAAGATATTTTTAAAACACGTTTTACCTGATTAGTTCTGAGTTTTGTGTTTTCAATGTTCTGCTTAATTTTCTTTCCTCATTTCTTCTTTTTATTAGTAGGGTGAGATGGTTTAGGCATAACAAGTTTCCTACTAGTGATCTATTATTTAAGTGATTCTTCCTGAGCTGCGGGGATGAAGACATACTTACTAAACCGGGTTGGGGATAGATTCTTTATAATAGCTCTAGTATTGTTTTTACATAATGGAAGATGGGATATAAAAAGCATAGATAGGAAAAATTTGTTAGCCTTACTAATTGTACTAGGAAGGTTTACAAAGAGTGCACAGTTTCCTTTCTCTAGTTGATTACCTGCAGCAATGGCAGCACCTACTCCTGTATCTGCCTTGGTGCATTCTTCCACATTGGTTACCGCCGGGATATATTTGATGGTACGTTTCAGTAGAATTTTCCCAGATTGATTATTTTTCTTGATAGGTATAAGAGGTATGTGAACTTTATATTCCGCCAGTTTAGCAGCCTGTAGTGAATTTGATGCAAAGAAGGTCGTAGCTTTTTCTACGCTAAGACAATTGGGATTAATGGCAGTATCTATTTCTTTAAATCTCCCAATGGTAGCTTTCTTTCATTTAATAACTCATGCAATGTTCAAGGCATTAATATTCATATGTGTAGGTTATTTAATAAATAATAGAGGACATTTTCAAGATCTTCGCAGTTTGAAAGGTTTATGATTGACTAGTCCATTATTAGCAATTACTTTAATAGTAAGAAGCTTATCTTTAATGGGGTTTCCATTTTTAGCAGGCTTCTTCTCTAAGGAATTAATATTAGAAAATAATATACTGATAATTAACCAAGTTTTTCATCTTTTATTATTGTTTTCCCTACCCTTAACATCTTATTATAGCTGTCGTCTTGTGTTTAATATATTAAATGGGTGCAAATATAACAGAGTATCTTGTCAAAAAGATGATAATGTCTTATTATTCTCTTTGCTACCTTTATATTTAGGTAGAATACTGGTAGGAAGTGCGTTATATCCTCAATTTTATAGATTAAGTTCAATTTATCCGTGTCATATGATGAAGTTTTTTGTAGCTCTTTTTATATTCAGAGGAGTATTTCTTAGTTGAAAGGATGTAAAGATAAAAAAAAATTCATTTATATGATTTAGAAGCACCATCTGTTTTTTGGTACCTTTTAAAAGTTCTTTTTGGAATAACAAGTTTGGTAACGTAGGAAGTGACTTATACTATCTTTTAGACCAAGGAATTTTAAGGAAATTAATTTATCACTTAGATAATAAAATCCAAAAATATGGAAGATTTTTTACTTCTTCTTTAAGCCTTTTTGATACTGTTAAAGGGAAGTATTATGTAATGGGAATTCTAGGATTAGGTGGATTTATAGGCATCCTAACTTAGATCATTTAGTGTTGGCGATTTTCAAGAAAAATAATTTCGTTTTGTTTTGAAAGCAAAAAGAGGGGTTAAAACCCCGAAAATCTCAAAAAGTTAGTATATAATAATTATAATTGGTTGTCGGCCAATAGATGAGAGTTTACTCCCACTTTTT**GTGAACAAAAATAATAAACAT**TTACGAGGATTTCCTAAGGATATTTATAAAACTCCTTTCCATTTAGTAGAAGTTAGCCCTTGACCCTTATTTGCCTCCGTAAGAGCTTTAGGGCTAACGTTTGGGGGAATATTTTGGTGGCATTATAATAATACTACTATTATAACTTTAGGTCTAATTTTAAATATATTAATAGCTTTCTCCTGATTCTTTGATATTATAAAGGAAAATATGAGAGGCTTTCACAAAAGTAGGGTAATGTTTGGGTTTCGTCTTGGAATGATATTATTTATCTTGTCAGAAATACTTTTCTTTTTCTCCTTCTTTTGAAGATATTTTCACAACTGTTGAGGTCCAAACTCAGAAGTTGGGTTTTTCTGGCCCCCTTATGAATTTGGGACTATTATAATAGATCCATTTTCAATACCATTATTAAATACAGTCATATTATTATCTTCTGGGGCGAGAGTTACTTGAGCTCATCACTCCTTGGTTAACCAGAGATATAATGATGCTTTATTGGGTTTATTAATTACCGTTTTTTTAGGAAGATATTTCTTATTTTTGCAAGGAACAGAGTATTTCTTAGCTGAATTCTCTTTAAAAAGCACAGTATATGGAACAGTCTTCTTCATGTTGACTGGATTCCATGGATTTCATGTAACTATGGGAACTGTATTATTATTAGTTTGTTTTCTCCGGCATTATTATATGCACTTTTCTGAAAATCAACATGTAGGGTTTGAGGCTTCTGCATGATATTGACACTTCGTAGATGTAGTTTGATTATTCTTATATTTCTTCATTTATTGATACGGATTTATGGTTTAGATAGGTTAAAAAAGTTTAAATTATTAAACACTACACTTAACTGTAGAGATGAGTATTTTACTCTTTGACCTTTGCTGTTGGCTATGATAGTTTCTCTGAGAATTTCTTTATTTTTTGTTGATGGAGGGGCAATTATAGCTTTACTAATGGTTCTGATATTATTC**ATGTGCCCATATTTAAGAGGGTTTCACTCACATTTTATCTATTTTCACTCGTTTGGCCTGTGTCTGGATAAAATGTCATCTTTC**TTAATTATATTGACTTTTTGAATTACAATGTTAATGATACTGTCGATGTGGGAAGCTGATAAATTGAAGTGACTATTATTCTGTTTTACATCTTTAAATCTAATTTTAACCCTAGCTTTTATGAGCAGAAGGTTACTAGGATTTTATATATTCTTTGAGTTATCTCTGATACCTACTTTGATTATAATATTAGGTTGAGGAGTTCAACCCGAACGAATTCGGGCGGGGAGTTATTTAATGATCTATACATTATTAGGGTCTCTGCCTCTACTAGGATGCCTAGTCTTCTTAGATAGTCATTGTGGAAGGGTGAAGATATTTTCCCCTCTAGTAGACATATTTTTCTTTTCCTATAAAGATTACAGACTCTTTTGTCTGTTATGAATTTTGGCCTTTTTAATAAAGATGCCAATTTATGGTGTACATTTGTGGCTGCCGAAGGCCCATGTAGAGGCGCCAGTAGCCGGGTCTATGGTTTTAGCGGGTATTTTACTAAAGTTAGGAGCATATGGCCTAATACGATCACTTCGATTCTTAGATTTAGATTACTGTTTTTGAAGAGACTTCTTCTTTATATGAGCCTTATTTAGAATGTGCTTAGTAGGATTAATGTGTTTCCGGCAATGTGACTTAAAGTCACTTGTAGCTTACTCCTCTGTGGCTCATATGTCTTTGGTATTTGCTGCTTGTTTCTCTTGTGATGTAATAGGAATGAAAGGGATAATGAGAATGTTAGTATCTCATGGATTATGTTCCTCTGGGTTGTTTTTCGGCGTGCAATGCTTGTATGAAAAAAGTGGCTCCCGTAGAATATATTTAAATCGAGGAATGATAAGCTTATCCCCTTTATTCTGTTTCTTTTGATTCCTATTATGTGTAGGGAATGCTTCAGCTCCCCCAAGATTAAATTTATTAAGAGAATTTATGCTAATCTCAAGAATAGTGAGATATGGGGGTCCTATAGCTGCTTTTCTTTGTGGGGTATCTGTATTTTTAGGAGGACTATTCAGAATATACTTATATGTTTCCGTTTGTCACGGTAAGTGATCCACATTAAGAAAATATTGAATGCCTTTTACTCTCCGTCATCAATTAGTTATGTTCTTGCATGTACTACCTCTTTATGGATTAGTTTTTTTAAGAAAATATATTTTCTATTTTTAAAAAAAATCTGTAGTTTAATAAAAACATATTGCTTACACCAATAAGATGGGGTTTACTCCCAGATTTAAT**ATGGAATTAGAAGTTTTACTTCCTTGAGCAATAGTAGTGATAAATATGTTGATAGCTATACCTTTTTTAGTAATGTT**TGAGCGAAAGGCATTAGGTTATACCCAAGATCGGAAGGGTCCAAAAAAGGTAAGATTTTTAGGTGTTCTACAGCCTATTGCCGATGGTGGTAAGTTAGTTTTAAAGGAATTCGGTTCCCCCAACTTAGGAAAAATAGTTCTCCTATGAATCAGGCCAATTCTGTCTTTTGTATTAATGATTTTGGCCTGATTCATATACCCTACTGAAAACTTAAGGTTTAGATTCAGATTAGGAATGGTATTCTTTATTTGCATATCTAGAATGCAAGTATATACTCTTTTGGGCTCAGGTTGAGGCTCTAACTCAAAGTACGCTCTAGTTGGGGCTGTACGAGGAGCCGCCCAAACAATTTCTTATGAAGTCTCTCTTATAGCTTTGATTTTTTTGCCTTGTGCACTTCAACATAGTTATAATCTTTATGAGTTCGTAAATAAGACATACCCTTATGTACTAATATTCCTCCCTCTAGTACTTGTATGATTTGTATCCTGCACAGCCGAGACCAATCGAGCACCTTTTGACTTCGCGGAAGGTGAAAGTGAACTGGTGTCGGGATTTAATATTGAGTACTCTGCGTTCGAGTTTGCTTGTTTATTTTTAGCAGAATATGGTAAAATATTATTAATGAGTTTTCTGACAGCCATTTTATTTTTTCCTGCAATTAAAGGAAGATTTTTTTATTTTGGTGCGTTTTTTACTTTTTCATTCGTATGAGCTCGGGCATGTTTCCCCCGATTCCGGTATGATTTCTTGATGAAAATCGCCTGGAAGACATTCTTACCAATAGCTCTCTGTTATTTATTTTTAGTAATATTATAAAAGGGATTACAATTTTAACAATGTTAAAAATGAGCTAAAGCTCTCTCTTTTAA**GTGGTTGGTCTGGAATT**CTTCTATTTTCTATTTTTTTTGAAATTGTTATTGGTAGTTTTTCGTTTAAAGAGCATCTTAGTCATTTTATTTGCCTTTGAATTTATAATAGTTAAAATATTCTTCTTATTGGCTTTTCTTAGAACTCCCCTTGATAGTTCATATGTTCTAATATTTTTAGCAATCATGGCTTGTGAAGCCAGTATAGGTCTAACCTTAATGGTATCAATATTACGACAAGGAGGCAAAGATAAAGTTAAAGTATCTTCTAGTCTGGCTATAGATAGTTAGGAATTTCGTGAGAAATTCATAAGAAGAATGGTTTCGGCCCATTATTTGAGTAAACCAGTACTCTCTCATGTATATGCCACAAATGTCTAAAGTACCCTTTTTATTAGTAAGGATTACAATCTTTCTTTTCTTTATTTTATGAATAGTAAATGTGTGATTTAACATTTTTTCTTTGTCAGAAAAAAGCTCACAATCAAACCAAAGGAATAGAGTTTCTTCTGTTTGGCCTTTGAAATAAATATCTTCGATAGCTAAAGGATAGCGTAGTCTTGAAGCGGCTAAGGTGATAATTTTTATTCTCGAAGAATAAAGTGGGGTATTCTAAAATTTTAAGAGGTTGGATTGTAGCTCCGAAGATGAATAATATATCCCCCACAAAGGAACTAATAGTATAATTATTACGTTGAATTGCAGATTCAAAGGTGTTATAAAACTTAGTTCTT**AATTTATTTGGTTTTGGTAAGATAAAGGATGGACATTAAAACGAGCATAAAAA**AATTTTAGAAGTTTATAACTTTAAGTTTTAGGTTATAGAATTTAGTCGTGAATTTTTATTAATAAAAACGAAAGTTTGTTTAAGTTGATGTCAGAAAGGGGCTAATAGTAGGTGCCAGCAGCTGCGGTTAAACCTATTCTTATCTTTATTTTCGGGTTAAATACAAATAAATTATGGTTAGTTTGGGAGAAAATAAATTATTTCGGTTAAATGGTAATAATTTTATTATTCAAACCCCAAATATGTAAATTGTGAAAATCACCACAAAAACTAAGATTAGATACCTTATTATTGGTGAAGGTAAATAAACATACCCAGAGTAGTAAGTGTTTTGAAACTCAAAAGACCTGGCGGTGGATGCTTCCTACCAGAGGAGCGTGCTAGGTAATAGATAGTCCGCTAAGTAATAAACCTTTCTTATTTTAGTCAGTGTACGGCCGCTTGCAGGTATTTCTTATTGGAAACATAAACTCCCTATAACATACTACTAGCTAGTTAAGACAGGTCAATGTGCTGCTGATAGAAAGGGTCAGAAGGTTCGCTATTTTAATATTAGGCAAAAGATATTACGCTGTAAAGTTTGATGGAATTCAGATTTGGAAGTAACAAGATATAAAATATGTATCTGTGAATAAAAGGTAAAGCTCTGTGTACACATCGCCCGTCGTTCTTTTCCAACGAAAAGAGAAGTCGTAACATGGTAGCCTTAATAGAAATTGAGGCTTAAGGGATAGTATAATAATATTATATAAAGCTGTTAACTTTATGATGTGCTTAAGCACTCTCTTAGTGAAATGGTTCTAGAAGCGATAAAAATTTAATAGCAGCAGATTCTTACTCTGTAGGTGGATAATTTAATTAATCCCTGAGCCACCATTTAAAAGTAGTTTATATTAAAATGCTTACTTTGGGGGTAAGCGAACAAGGAAAACCTTGCTTTTAA**ATGTCTATAGGT**TTATTGATTGCTGTAATAATATCGAAAATTTTAGTATTGCTAATAATAGCTTGAATGGTATGAACTACTCGATTTAATTATGCTAGGTGAAAGGAAAAGGCGATAGCCTTCGAATGTGGATTTGACGTAAAGGACTTTGCTCGTATGCCATTTTCCCTCCGATTTTTTCTACTAATAATATTATTTCTGATTTTTGACGTGGAATTATCCTTTTTATTACAACTCCCTTTCTATTACGAAGGAGAATATAGAAAGGGACGAATAGGCCTATTAATTTTTACCTGAATTCTATATTTAGGAGCTGTCGAAGAATGGCGTCGAGGGATACTAAGTTGAAAGATTTAGTTTGAGGTGGTCAAATAAGATAGTAAAAGCTGCTAACTTTAGCTAGAGGACATTGTACCCTTCCCACCTCTATTGCTAACTAGAGTAAACAACCGCGCGTATTTTTTACCCCGATTTTGGCTTAGAAGATTTTTATTA**ATAGTCGCGGCCAG**TTTCGGGGTTTTTATATCTATATTTAGAAGAAATATATTCTTAATCTGGTTTGGATTAGAAATCAAAATGTTCAGCATAATTCCCTTTATTAATATAAGGGCTGAAGATGAAAAAGAACGACAATTAACTAAGAGTAATCAAGTAAAAGTGTCATTTTTCTACTTCTTTGTACAAGTTCTTGGAAGCCTCTTATTTGCCTGGGGGAGAGTCGCGAGAGATGCCTTCTTCTTAGGTTTAATAGGCCTAATAATTAAGATGGGAGTAGTACCTTTCTTCTGATGAGTACCCCCTCTAATTTCTCGCCTTGACTGACTATCTATAGGAGTAATGAGTACTATACAAAAGATACCTGGATTATTTCTTTTTCGCCTGTTATTTGATATATCTTTCAATATGTGTATTGCCATAGGTTTAATAGGTCTAATAATCTCTGCTATAGGGATTAATCTATCTTATAAAAACTTCAAGCAATTAATAGCCTGGTCATCTATAAGTAATATGAGAATACTATTCGTACTAGTTGTGATTAATAGAAGATTAGGTCTAATCTACTATATATTTTACAGATTAATGATTTTAATGTTTTGCTGTATTCTAGCTTATTCCGATAAAAAGGAAATAAGTCACTCCTTTACTCAAGGTAACACCACTATAATAGATATGTTGGTAGGAAGACTATTATTAATATTTTCAGGTCTCCCTCCCTTTATAAGATTTATACTTAAGGTGTATTTTCTAAGAGGTATCTATTCCTTAGATTCCGTGTTCGTATTAAATGATCTGGAATTGAAAAGTAGAAAAATGACATTCTTTTATTCATTAAGAAAATCTTTAAGAAGCTGAAACTTAGCAATTATATTTTTATTCTTAATGGTTTTACAATCTGTAGGTTATATAAAGGCTTTTATTAATATGTATAGTTCCAAAGAGTCTCGATTATTAAATAGATGCAAGGGACCCTGAACATTTTTAAGATGTTTTACCTCAATAAGCCTATTAATATATTTATTTAGAGTATTCACAATATTATTTTAGTGAGACGATACCTGCTTAAAAGTCAAAAATGCGACGTAAGTCTTGTAAACTTAAGATTGAATCTATTGATTCTTAAGCAAAACATTATAATTATTTATTTCTCCTAGTAGTTTATTCAAAATATTAGTTTTGCGTACTAAAGATGGTGAGGACCCTTGGTGGCATAACGTGAGGACTGCGAATGCAGGTCACTGTGATATAGTGAACTTTAGATTAGAATCTCCACGTTATTTTG**ATTTATGATTTATTTAC**TAGTGCGGATTTTTGTTATTCCAATTTTTCATTACTGAACATAGCTGTTTGATATTTAGGTTTAGTATGATTATTTTGAAAAATAAATTTGTTTTTTTACTCAAGCTCTCGGTTAAATAGATTATTAGCCTTTTTA**ATAAATTGAC**ATTATGAAAAAGTAAAGTCTTATTTTTTAAAAAAGGTAAAGGGTTCTTATATTATTATTTCTGGTTCTTTTCTAATTATACTAGGATGTAATATATGGGGTCTATTCCCTTATATATTTGGAATTACTACCCAAATGGTCTTAACCTTTACGATGTCAATGATAATATGATCTTGCATAGTATTTTCAAGTTTAGATTACTCCTTTATAGGCTTCCTTTCTCACTTAACTCCTCAAGGTTCTCCTGGATACTTAGCCCCTATTTTAAACTTAATAGAGCTAGTAAGCAAAATTATTCGGCCCCTAACTTTAGCCTTACGGTTAAGAATTAAAATTACTACAGGTCATGTATTTATAAGTTTAATGGGAACAAGAGGAACAGCATGTTTATTTCCTTTTAAGATCTTCTGAATACTCTTTGTGATTTTAATGAGAGGCTACTTATTATTCGAGATAGGTATATGTTTTATTCAAGGTTTTGTTTTCAGTTTATTAAGAGTACAATATCTAAGAGAACATACGTAAAGAATTAGTAGTTTAAATTAGAATTCTGCTTTGTGGGTGCAGGGGTATACCTTTAGAGTATCATTCTTCTGTTTCTTTCTAAGCTATATATAAATTCAATACGTCGAGATCGGTCT**GTGTTAAAGAGAGTTAACGGAGGAGTTTATGATTTACCCTCCCCAAAGAAAATATCATACTGATGGGGATTTGGTTCTCTATTAGG**CTTATTTTTAGTGATACAAATATTAACTGGATTATTCTTATCCATGCATTATGTTTCGGACTTAAATGTGGCTTTTCTCTCTGTAGACTCAATCAGGCGAGAAATTGATTTTGGTTGATTAATACGAAGTGTACATGCTAAAGGAGCATCTTCGTTTTTTCTATTTTTATACTTACATATTGGGCGCGGAATTTATTATGGTTCTTATTTATTTAAGCACACCTGAAAAACAGGAGTAGTTATATATATATTGTCTATGGCTACTGCTTTCCTAGGATACGTATTACCCTGAGGTCAAATGTCCTATTGAGGCGCTACAGTAATAACTAATTTTTTCTCCACAGTACCTTATGTAGGAACAGATTTAGTGCAATGAATCTGAGGAGGTTTTGCTGTAGGATATCCCACACTAACACGCTTTTTCTCTTTACATTTTCTATTACCTTTTGTAATTACAGGGCTAATAGTAATACATTTAATATTTTTACATGATACTGGGTCTAATAATCCTTTAGGATTAAGTTCTGCTGGAGATAAGATCCCTTTCCACCCATATTTCACGGTGAAGGATATTTTAGGATTTGGAATCGTTTTTTTCATTTTCTTTTTAATTGCGATAGGTACTCCAGATTATTTCGGAGATCCTGAAAACTATATTGAAGCCAAACCTTTGGTAACTCCAATCCATATTCAACCAGAATGATATTTTTTACCAGCTTACGCCATTCTTCGAGCCATTCCAAATAAGTTAGGAGGGGTAGTTGCTCTTTTGATGTCCATCCTGGTATTATTTTTATTCCCTGTAGTTTCTTCTTTTTGTAATCGAGGATATTTTTATTCTGTAGGACATCAGCTATTATTTTGAAGGTGAGTAAGAGATATACTAATATTATTATGAATAGGAGCTCGACCTGTGGAAGAGCCCTATATCTCAATAGGAGCTTTCTGTACTCTATATTACTTCTTATTTTTCCTGCTAACTCCAATAGTTAGACTAATTTATGGTAAGCTATTCTCAAAGGGAAGCCTAAACGTAAGATATGTTACCGTTTCAGAAAATCCTGAAAGAAAGAGGTTAATGTTGAGTTAAAAATAATCTGGAGTGGCAGAAAATATGCGCTGGATTTAAGCTCCAGAAATGAACAATTTGTTCCTCCAGAGTTAAGTATAAGGCGGTGTTACTAGCTGCACGTAAGTTTTTGGTACTTAAAGAAAACGTAGTTTCCTTATAAAATTATAGGCCAATTAAACATAGCCGTTTACCTCACTTAAAATGAGCTTAAATAGAAGTCTAATTTAATCCTTGGCCTATAAGTATATATACCCTCCCTCTATTAATATAATTAAAGAAAAAAATTTCTTCCAAAAAATTTTATTAAAATATTATCTGATTGAGTAAATAAAATTATAGGCCAATTAAACATAGCCGTTTACCTCACTTAAAATGAGCTCCCTCATTTCACATAAGGGGACCCCCTTATGTGAAAAACAGGTCCTCTTTATTGACTTAGATATTTACACCAAAAATAGATAATAAGTTATAATTAAACTATAGGGCTCATGCCCCTAAAAAGATCCAATCTTATCTA**ATGAAGATCGGTAGATTTGTCGGAAGTGGTTTTATTT**ATGGTATGAATTTACAAGATGCTTGCAGACCGGTAATGCGTGAAGCGGTATTTTTTCATGACGAAAAATTAGTGGTTTTAATTGTAATAGCTTCTCTAGTAGGAGGAAGCATAGCAATGTTATGAATTTTAGATAGAAGATCCTGTGATTTTATTGATCATAAGTGATTAGAAGTAGGGTGGACATTATTCCCAGTGGTTGTATTATTAGTACTAGCTCTCCCCTCTTTAGAACTTTTATATTATAGGGATAGACCTTCAACTATTACACCTCTGGCCACATTAAAGGCAATAGGTCGTCAATGATATTGGTCTTACGAAGTTAGAGTTCGTGGAAGCGAAGGTCCTACAGGTTTCTGTTATGATTCTTATATGCTACCTCAGGGACAAGAAAAAGAAGCTGAGGGAGACTTATCAGGCTTTCGTTTATTAGAAGTTGATAACCCTATATTCTTACCTCGAGGTGAATATATTCGTCTCCTTGTGACGGGAGGGGATGTAATTCATAGATTTTGTGTACCTAGTTTAGGTGTAAAGATCGATGCAGTACCTGGCCGGCTAAACCAAACCTTTTTTTTCCCATTAAATTTAGGAAGATTTTTTGGCCAGTGTTCAGAAATATGTGGAGCTAATCATAGTTTTATGCCTATTAATATTGAAATAATAACTCCAGATGTTTTCTCTCGCTTTTTTACCTAGACTTTTAGGTTAATTTAGATCATTAGCCTTCAAAGCTAGGGGAGAGAAACTGGAATTCTCAGAGTCTGTGT
