## Supplementary for "Mitochondrial Genome-Based Phylogeny of Turbellarians and Evidence for Accelerated Mitochondrial Evolution in Symbiotic Species": annotationDiscocelis tigrina MF993333.docx

A**TGAATATCTCGTT**GGTTATATTCTACTAATCATAAGGATATAGGAACGTTATATTTAATTTTCGGTATATGATCCGGGTTTATAGGAACGGCTTTTAGTTTTTTAATCCGTTCCGAGTTGGCTCAACCGGGAAGTATATTGAATGACCCACAACTGTATAAAAGAATTATCACCGCACACGGTTTAATAATGATTTTCTTCTTTGTAATGCCAGTACTTATAGGAGGTTTCGGAAAATGGTTAATACCACTATATTTAACCAGTCCAGATATGGCTTTCCCCCGACTCAAAAAAATGAGGTTTTGATTATTACCACCCGCACTGTTCTTACTACTAGGATCTTTTATTGTTGAGGGTGGTGTAGGTACCGGCTGGACAGTCTACCCCCCATTATCAAGAAATATAGCTCAAAGGGGACCAAGAGTTGATATGGCTATTTTCTCTCTACATTTGGCAGGAGTAAGATCTATATTAGGTTCTATAAACTTTATTACTACAATGGTAAAAGCTAAGGTTCAAGTTTCTTGAGGACAAATGCCATTATTTTTATGGGCTGTGATGGTAACAGCTTATATGTTGGTATTATCACTACCAGTCTTAGCTGGTGGATTAACTATGTTATTAACTGATCGAAACTTTAATACCACCTTCTTTGATCCGGGGGGTGGGGGAGACCCTATATTATTTCAACATATATTTTGATTTTTTGGACATCCAGAAGTATATATCTTAATCCTCCCCGGATTTGGTATGATTTCACAAATAGTTACATATTACAGTGGAAAGGAAAGTGCTTTTGGTCATATGGGTATGGTATATGCTATTTTGGGAATAGGGTTTCTAGGATTTATCGTTTGGGCTCATCATATGTATACTGTCGGTCTAGATATAGATACACGAGCGTATTTCACAGGAGCTACTATGATTATAGCTGTACCTACAGGTATAAAGATATTTAGTTGGTTAGCTACTTTTTACGGCCGACCCTTAGGGGAAGGTATGGATAGCGCAGGTCCAATATGAGCAACAGGATTTATCTTCCTATTTACTTTAGGTGGTTTAACTGGAGTGGTCTTAGCTAGAGCAAGTTTAGATATAAGACTACATGATACATATTATGTGGTTGCACATTTCCATTACGTACTATCAATGGGAGCTGTGTTCTCTATATTTGCTGGGGTAGTACACTGATGGTCTCTGTTTACTGGAACAGTACTAAATACAAAGATGGCTATAGCTCAGTTTTGAGTACTATTTACAGGGGTAAATCTAACCTTTTTCCCTCAGCATTTCTTAGGATTGAGAGGGATGCCCCGACGTTATACAGATTATCCGGATGGTTTCGCATATTGAAAAAGTGTATCGTCATTTGGTTCATTAATATCAGTTATAGGTGTAGTATTTTTCGTAGTAATTATATGGGAAGCTATGTTAAGAGAACGTTTAGTAATTTATATAGTCTCTCCTAGCTCACAGGTTGAGTGGATTTGTTCAGACAATTACCCATTTAGTTTCCATACAAATGAAGTTGTTAGAAAGGTTTATGTAAAATAAATATAAAAGATTCTTTCTAGTTTATAAAGAATATTTGTCTTCCAAACAAAGGGACCTTAATATAAGGGAAAGAAAAAAAATTACACTTGAGATTTCGTAATGTCTAAAATCTTGATAAAGATATTATATACACTTATTATATTGGTTAAATCTTACCTTTTGTATCATGATCTAATAAATTTTTATAAATATTAATATA**CCCGAAACTTTTGGATTATATA**TCATCTAGTTAGGAACAAATTTTTTCCAGGGTTTTGTTAAAAAAAGATGGAGAGATAAGAATAAATATTATAGTCGGCAAAAGTTATATCTGGTTAGAAGATTAATAATTTCTTTTTAAGATGTGGGGTTCAAGTTTCACATTTTATATATAGGTAAGATTATATAATTATTGTTTTAGATGAGAAGTGTCTATTAACTAATAAGTATTATAACTGTAATACGTTTTATTTTATACCTTAGTTAGAGTTTTCTAAATCAACATATAAATATATTGTTAGAATAAGTAAAATAAATAAATTCATAAGAACTCAATGTTTTTACAACTGTTTATCAAAAACATTTCCTTTAGTAAATATTTAAAGGTATTACCTGCTCAATGTTTAAATATAAATTGCTGCGGTATCTTAACTGCACTAAGGTAGCATAATTAATTGTCTTTTAAATGGTGACTTGTTTGAATGGTTATATGAGAAAAGAATTTTTTTTGAATTAATATGAAATTATATTCTTTGGTCAAAATACCAAGATATAAAAGAAAGACGAGAAGACCCTGTTGAGTTTTACAATACATTTATTGTTTAATGGGAAATTTGATTTCATTTTGAAATCTATTAGATCCTAAATTTTTTATTTGGAAATAGGATAAAGTTACCACAGGGATAACAGGATAATATTGACTAAGAGTTCTAATCGAAGTCTGTGATTATTACCTCGATGTTGAATCGAAATATCTATTTAATGCAGCAATTAAATAAGAGGGTCTGTTCGACCTTTGAAATTTCACGTGATTTGAGTTAAGACCGGCGTGAGCCAGGTCGGTTTCTATCTTCTTTTGAATCAGTTTTGTACGAAAGGAATTGATGATTTAATACTTTTTAAATTCGGGGTGGCAGAGTATATGCGGCAGAATTAGAATCTGCAGATGAGTGAAAAACTCCCCTGAAAGATCTTTAAGAAGTTTAGTTCGTTTTGTTTTGAAAACAAAAAGAGGGATATATATTCCCAAAGATCTCCCTTATTTTAGTGTATAACTGGCACATTAAGTTTTCTCCTTAAAAGAGTTCTATTAGAACAAATGAGATTAACGATTGTCA**ATGAGCATTTGATTTTTATTAT**TAGTTGTTAAATATTTTTGCCTTCTTTCCTTAAAATCTTTAATTTTAGGCATTTTCATATTTATATGTAGTTTGTGTGTTAGACTATTAATAAGAATATATTCTTTATCCTGATACTCTTTGATATTTTTCTTGGTTTATATAGGTGGGTTGCTAGTATTATTTATATATATATCATCATTGAAATTCAATCCAGTATTCTATTTTTCTAAGTTATCTCCTCTTGCTAAAATATTATTAAAGGCCAATATATTTTTTTTCTTATTTTTTATAATAACTCAAATCACGTGAAATTTCAGAACCATTAGGTGGAGAAATAGAGATATGAAAAATTTTAGTTTAAATTTATTTAATGAAACCGAAGTACTATTTCTAATAAATGTGGGAATATTATTATTAATTGTTTTATGGGTAATAACAAAGCTGTCTTTCCGAAATCGGAGAGCACTCCGACCTTTTTTTGGTTAATATT**ATGGAACGACACTATACAGTGAATGTAATTAATTACAAT**TTAAAGTTTGGATTTCCTTATACTTGTTCCAAAGTATTCTTTTTATTAAGAATTTTATTCAGCTTGTTATTTATGCTTAATAAAAAAAGTTATTTATTGATATTAAGTATATTAGATACTTCATCATTAAACCTAGATTTTTGTATCTTAGTAGATTGAATTAGATTACTTTTTTTTGCCACTTTGTGTATTATAGTATCTTGTGTTTTAAAGTTTTCATGTGTATATATGGAAAAAGATCCATTTCAAGTGCGGTTTACATGATTGGTCTTAAGTTTTGTTTTATCAATGGCATGCTTAATATTTTTCCCCCATTTCTTCTTTTTACTGATAGGGTGGGACGGTTTAGGAATAACAAGATTTCTGTTAGTAATATATTACTTAAGGGATTCTTCTTGAGCTGCAGGAATGAAGACCTATTTAATTAATCGAGTGGGGGACGGATTTTTTATAGTAGGCTTAGTATTATTTTTATATAAAGGTCATTGAGATATAAAAAGTATAAGAGAAAATAATATCTTAGGCATTATAATAGTTTTAGGGTGTTTTACAAAGAGGGCTCAATTTCCCTTTTCAAGATGGTTGCCAGCCGCAATGGCCGCACCCACCCCCGTGTCATCTTTAGTGCACTCATCTACATTAGTTACTGCTGGTATATATTTAATGATTCGATTTGCTGATATATTTTCACAAGAAGTATTCATCCTAATTGGGATATGTGGGTTGTGAACTCTTTATTCGGCCAGGTTAGCTGCTTGTTCTGAATATGATGGTAAGAAGGTGGTTGCATATTCTACTTTGAGACAGTTAGGCTTAATGGCAGTTGCAATTTCATTAAATCTTCCTATATTCGCTTTCTTTCATTTAATAACTCATGCAATGTTCAAGGCTTTAATATTTATATGTGTAGGATATTTAATTAATAAAATGGGACATTTCCAAGATCTCCGGAGCTTAAAAGGGTTGTGGATTACATCACCAACATTAGCTATAACTTTAATGGTTAGGAGATTGTCCTTATTAGGCTTCCCATTCTTGGCAGGCTTCTTCTCCAAGGAGTTAATATTAGAGAATCAAATTTTGGTAGTAAATAACCTTTTTAATTATTTGGTGTTACTATCATTACCTTTAACATCATATTACAGCTCTCGGCTAACATTTAATATATTAAATGGGGTTAAATATAAAAGAATAAGCTGTAAAAATGATGGAAAAGTATTATTATTTTCTATATTTCCTCTATATATGGGGTGTATCTTTATAGGGTGTGCTATATCCCCCACAATACCCAGTTTTAGTGGGGTTGTATCCGCCTGTCATTTAAGTAAGTTCTTTACAAGGTTCTTCATTTTGTCCGGCATAATATTATGTTGGTTTGACGTAAAGATAAAAAAAAATAGTTTTATTTGGTTTTGCAGCTCAATAAGTTTTTTAACCCCATTTAATAGGAGGTTTATAAATGATAAGCTCAGAAAAACAGGTAGAAATTACTATTATATATTAGATCAAGGAATATTACAACAAACCATAAGGTCATTAGATAAGGGGTTAACAAAAAGAGGTAGTTGATTATCCAGTTCATTAAGATACTTTACGTTTCCTCGAGTTGGCTATTATGTGATAAAAATATTATCTTTAAGATTTTTACTAGGAATTGTAACTTAAAGATTCAAAGAGTTAGTATATATATTATAATTGGTTGTCGGCCTTTAGAAGGAATATTTATTTCCACTTTTTGTTGAAAGCATCTATTATAAAACGAGTAAACATATCAAAGAGTAGATATAGAACGCCA**TTTCATTTAGTTGAGGTAAGCCCATGG**CCCTTATTCGCTTCAATTAGTGCATTGGGCTTAACTTTCGGAGGTATTTATTGGTGACATTATAATAAAACAAGAATAGTTATATTAGCATTTATATTTAATTTAATCATCTCCTTCTGTTGATTTGGAGATGTAATAAAGGAAAAAATGACTGGTTATCATAATAGGGTAGTAATGTTTGGGTTTCGATTTGGGATGATACTATTTATATTATCAGAGGTTCTTTTCTTCTTTTCTTTTTTCTGAGCTTATTTTCATAATTGCTGGGGTCCACAGAGAGAGCTTGGGTTTTCTTGACCCCCTTATGATTTTAATAATATAGTTATTGATCCATTTTCTATACCTTTATTAAAAACGGTAGTACTATTGTCCTCCGGAGGGAGAGTTACTTGAGCACATCATGCTTTGGTTAATCAAGATCATGATCAAGCTGCTATAGGCCTTTTTGCTACAGTATTTTTGGGAGCTTATTTCTTGTTCCTCCAAGGAAAAGAATACTTCTTAAGAACATTTTCTATTAATAGAACAGTATATGGAACAGTCTTTTTTATGTTGACTGGATTTCATGGATTTCATGTAACTATAGGCACTATTTTATTGTTAGTATGTTTTTTTCGACATATTTTAGTACATTTTTCTAAAAACCAACATGTTGGCTTTGAGGCCTCGGCTTGATATTGACACTTTGTTGACGTGGTTTGATTATTTTTATACTTCTTCATATATTGATATGGTTTTAATTTATAAATATTTTAGGGTGGCAGATAATATGCACTAGATTTAAGCTCTAGGGATGAGTATAATTACTCCTCTAAATATGGGAGAGTGTATTCAGCACGTTAGTTTTTGGTACTAAGAGTAAACGTAGTTTTCTCATAAATTTGAAAGCGAATAATAGCAGCGGATTCTTACTCCGTAGGTGGGTTGCATTACCCTCAAATTTTAGGGGTAGTTTAATAAAATATTAACTTTGGGAGTTAAAGATCGAAAGTCCTATTCGTCCCTAAATT**ATGTTATCAA**TGGGAGTAATTTCTATCTTAATATTTATGATATTCAGTTTATTAGTAATATTATGAATTGTATGGATGACCCGAAAGAAATTTTCAAAATCACGGGAAAAGGCAAGACCTTTTGAATGTGGTTTTGATCCGAAGGATAAGGCACGAATACCCTTTTCCCTCCGCTTTTTTTTAATAGTGATACTATTCCTTATTTTTGATGTAGAATTGTCATTACTATTACAATTGCCTTATCAATTAGATTATGAGCATTTTAAGGGACGAATAGGAGTGATAATATTTACTTGGATTCTACTAATAGGGACCTTAGAGGAATGGCGACGGGGAGTTTTAAATTGAAAGAAATAATTTTAGGGAAGTTCAAATTGATTCAAAGAGCTGCTAACTCTTTAAAGAGTAATTAATAATTATTCACTTCCCTTAA**ATGTCAGTTGC**AAATATTCGCTTTCTGCCAAGAATGTGAGGAAGATCGGTGGGATTAATATTTTGTTTATCTATAGGAATATTAATATCTCTAGTTAGGAATAAAATATTTTTGGTTTGATTTGGACTTGAATTAAATATGTTTGGTATTATACCATTTATTAATTCAAGTCCAAATCAAACCAAAAATATTTTATTCCTAACTCCTCAGGAAGTTAAAGTATCTTTCTTTTATTTTTTTGTACAGGTGATAGGAAGGCTACTTTTTGCTTGGGGGGGAATATTAAGGGATTGGTATATAGTAAGTATTATAGGTTTAACTATTAAGGTGGGAGTTGCTCCCTTCTTTTGGTGAGTACCACCTTTATTAACTCGTTTAGATTGATTGTCTATAGGAGTTGTTAGAACTATTCAAAAGATTCCTGCTATATTATTAATTCGGTTGGTGTTTGATTTAAAATTAGAAATATGCTTACTGTTTAGTGTTTTGGGGTTTTTGGTGGCTACGGTAGGCATAAAATTTTCTTCTAAAAAATTGAAGCAGTTAATAGCTTGATCATCAATAAGGAAAATGAGGATATTAGTAGTTTTAATAATCCTAAATAATAGGTTAGGATTATTATATTATGTTTTTTACAGAGTATTAGTTTTGATTTTTTGTATTTGTCTTAAATTTACTAATTCTGACAATCTTTGCAATTCTTTTATTAAAGGGGAAAAAGAAAAAATCAAAACTATTAGAAGTCTACTTCTCCTTGTTTTCTCCGGCCTCCCACCCTTAGTAAGTTTCCTTTTAAAGATCTTTTTCTTAAGCGGTTTCTTTTTGAAAGACTGTTCCTTTCTGATAATGGATATTGATATGAAAGGTAAAAGAATCTCCTTCTTTTATTTATTAGGCAGGTATTTAAATAGATGAAATATAGTGACGCTTTATATAAGATTAATAATTCTACAGTCTATAGGATATATAAAGGCTTTCATAAGTATAAAATCTAGTAAGTATTCACGTCTTAATATAATATCAAATTCTGTAAAAAAAACAAAAAAGATAATTTTAATATCTTTAGGTATATTATATTTTATCAGTCTTATATTGATATGGTTATAATTATTGCTTCAAAGTCAAAATAGAGACGTAAGTCTTGTAAACTTAAAATTGAATATTATCTTCTGAAGCAAAATTAAATCAAACGGTAGTTTAAAAATAAAACGCCAATTTTGCATGTTGGTAATTGGCCAATCCACTGTTTGAAATGTGAGGACTGCGAATGCAGGTTACTGTGATATAGTAAACTTTGGATAAAAATATCTCCACATTAATGGTGTATGATTTATTCACTAGAGCAGATTTTTGTTATTCTAATTTTACATTTATAAAAATAATTGTATGGTTTTTAGGACTATTTTGATTATATTGAAGAATAAATTTATTCTTCTTTTCCTCATCCCGGTTAAATACTATTATATCTTATTTACTTACATGACATTATGAAAATATTAAGTCTGACTGTTTACAAACAATTAAGGGATCTTATATATTGATTTCCGGCCTGTTTGTAATTATTTTAGGTAGTAACATATGAGGCCTTTTCCCTTATGTCTTTGGGGTTACTACTCAGATGGTGCTAACTTTTACACTATCTTTAATTATATGATTAGCTATAGTAGTTTCTTCAATAGAGTACTCATTTATGGGATTTTTATGTCATTTAACTCCACAAGGAGCTCCTGGTTACTTAGCTCCCATATTAAATTTGATTGAATTAGTAAGTAATTTTATTCGTCCCTTTACTTTAGCTTTACGATTGAGAATAAAAATGACCACAGGTCATGTATTAATTAGTTTGATGGCAACTTCAGGGGTAATATCTTTATTTACATTTAAGTTTTTTTGAGTTTTGATGACTTTATTAATTACAGGTTACATACTTTTTGAAATGGGTATATGTTTTATACAGGGATTTGTATTTAGCTTATTAAGTACTAATTACTTAGGAGAACATACTTAATATATAAGAGTAGTAGTTCAATTTTAGAATCTTGCTTTGTGGTTGCAAAGGTATACAATTATATGTATCTCTCTTCAATGAAAAGTAATAATTTATCTATAAATTCTTTTCGACGGGATAAGACCGTTCTTAAAACAGTCAAAAAAGCAATTTATGATTTACCTTCTCCCAAGAAAATATCGTATTTGTGAAAATTTGGCTCCCTATTAGGTTTATTTTTGGTTATACAAATATTAACTGGTTTATTTTTAGCAATGCACTATGTTTCGGATTTAAAAGTAGCCTTCCTATCAGTCGACTCTATAAGTCGAGAAATAGATTTTGGTTGGTTAATACGAAGTATACATGCTAACGGGGCATCCTCCTTTTTCTTATTCTTATATTTACATATTGGTCGGGGTATATATTATGGCTCCTACTTATTTATTCATACTTGAAATAGAGGTGTAATGATATATATATTATCCATGGCTACTGCCTTTTTAGGCTATGTGCTTCCTTGAGGACAAATGTCCTATTGAGGGGCAACGGTTATAACTAAATTTTTCTCTACTGTTCCTTATGTTGGAAAGGATTTAGTGCAGTGGATTTGAGGGGGGTTCGCCGTTGGTTATCCCACTTTAACCCGCTTCTTCTCTTTACATTATATTTTACCTTTGGTGATAGCCGCATTTGTGGTTATTCATCTTGTGTTTTTGCATGACACGGGCTCTAATAATCCTTTGGGATTAAGTTCTGCAGGAGATAAGGTTCCTTTCCACCCCTATTTTACAATAAAGGATGCCTTTGGGTTCAGTATAATATTAGGCGCGTTCTTTTCTATTGCGGTTATGGCACCTGACTATTTTGGTGACCCAGAGAAATATATAGAGGCTAACCCTCTGGTTACTCCTGTACATATTCAACCTGAATGGTACTTTTTACCCGCTTATGCTATATTGCGAGCTATACCTAATAAGTTGGGTGGGGTGGTTGCACTATTAATGTCAATTTTGGTGTTATTTCTATTTCCTATAGTATCTTCATTTTGTAATCGGGGATACTTTTATTCATCGGGCCATCAAGTATTATATTGGCTATGAATAAGTAAAATATTAGTACTATTATGGATTGGGGCTCGGCCTGTTGAAGAACCTTATGTTAGTATTGGTTGATATAGGACAGTAATATACTTTATGTTTTTTTTGTTAACCCCTTTATTAAGTCTGAGGTACAATAATATATTTAGAAATTCAGATACTTCTTCTTTGAATCTTACCCCAAAATGGAGCTTAAAACCTCATT**TGGACCGGAAAT**CCAAGATTCATATA**ATGTATAGGATATTATTACCAATTATATGCACAA**TTATATTTAGGGACTTAAAATATATTGTAAGTTTATTTAGGATTCTTATCATACTATGTGTACCTTTTTTAAGAGCCTCCAAAATAAGGCCTTTCTTTCATTCATATGGATTGTGTTGGGATAACTTATCGGTTTTTTTAGTAATATTATCTATATGGATAACCTTATTAATGTGTATTTCTATGAAAAAAGCACCCAGGTTAAAAAAATTATTTCTATGTTTCACTAGTTTAAATACCATTTTAGTCTTATCCTTTTTAAGCAGGAGCCTTTTGGGTTTTTATATATTTTTTGAATTATCTTTGATACCCACCCTTATAATAATTTTGGGATGAGGAGCTCAACCGGAACGTATTCGGGCTGGAAGATATCTTATGATTTATACGTTAGTAGGGTCCCTCCCCCTATTAGGGGCCATAATATATATGGATTTTCATTGCGGGAGGGTAAAGATGTTTAGGCCCCTAATAGATATATTTTTTTTTAGTAGATTCGACTATAGGTTGTTCTGTATACTTTGAATCTTAGCCTTTTTAATAAAGTTACCAATATACGGTGTTCACTTATGATTGCCTAAGGCTCATGTAGAGGCTCCAGTAGCAGGCTCTATGGTTTTAGCAGGGGTACTGTTAAAGTTAGGCGCTTATGGGTTAGTTCGCTCTTTGCGCTTTCTTAGATTGGATTTATCATTTTGAAGGGATACCTTATTTATATGAGCTATTTTTAGTATGTGTTTAGTGGGAATAATGTGTTTCCGTCAATGTGATCTAAAGTCTTTGGTTGCATATTCTTCTGTAGCACATATGTCATTAATACTTGCATCTTGTTTCTCATGTGACATTATAGGTATTAACGGTGTTATGGGTATGTTAATATCCCACGGATTGTGTTCTTCTGGCTTATTCTTTGGCGTCCAATGTTTATATGAAAAAAGAGGATCCCGGAAAATTTTTTTAAATCGGGGAATATTGTGTATTTCCCCCCTCTTTTCCTTCTTTTGATTATTATTATGTGTGGGTAATGCCTCCGCACCCCCCTCATTAAATTTATTAAGGGAGTTCTTTTTAGTATCAGGAATATTAAACTTTGGTGGTAGACTATCCGCTGTGTTTTGTATTTTATCGGTCTTTTTAGGTGGGCTATTTAGGATATATTTATACGTATTAATATGTCATGGTAAGTGAAAATCATCAAATAAATATTGATCATTACACTCTATGCGAAAATATATGGTATTATTTCTACATTTCTTTCCTTTATACGGTTTATTGTTTTTGAGAAATTATATATTTTTTTAATCATAGATTTATATTTTATTTAAAATATGTTGCTTACACCAACAAGAAAGATACCTTATCTTAAATCTAAAT**TTGAATCCATTCAT**ATTATTCCCCTGAGTATTAATAATATTAAATATATTTATAGCGGTACCTTTTTTAACCTTATTAGAGCGTAAGATATTAGGCTATGTCCAATCTCGAAAGGGCCCCAAAAAGGTAAGTTATTTAGGGATTTTACAACCCATAAGGGATGGATTTAAGTTAATATTGAAGGAATTAGGTACTCCTAAAATGGCAAAAGCCTTATTTTTTTGGATAAGTCCTGTAATATCATTATTATTAATGATCTTGGCTTGGTCTTTATTTCCCTCCCCCTATAGGTTCTATACTTTTAATTTAGGAATGGTATTTTTTCTATGTATAGCTAGCTTACAAGTATATACATTGTTAGGTTCGGGGTGGGGTTCAAAATCAAAGTATGCCCTTCTAGGGTCTGTGCGCGGAGCCGCGCAAACAATATCTTATGAGGTATCTTTAATTTTTATTATCTTTTTCCCTTGTAGATTGGAATTTAGTTATAATATACATACATTTTTAGACAAGAGTATTCCCTATCTTTTTTTCCTATTTCCCATATTTTGCATGTGGTTTATTTCATGTTTGGCTGAAACTAATCGAGCTCCCTTCGACTTTGCTGAGGGGGAGAGTGAGTTGGTATCAGGTTTTAACATAGAATTTTCTGCGTTAGAATTTACTTGTTTATTTTTATCAGAGTATGGCAACATTCTTTTAATGAGATTTCTCACGGCAATATTATTTTTCCCCACCACTAATTTATCATTTATAGCTTTTGGTTCGCTCTTAGCTTTTAGTTTTGTATGAGTGCGGGGCACTCTCCCCCGACTCCGCTATGATTTATTAATGGATATGGCATGGAAGATATTTTTACCATCAGTATTATTAACGCTGTTCATTATATTAGTTTAGTTCAAGATTGGATATATTTTGGTATTTAGCCCCATCTTATTTTGTTTAGTTTGGATATCTTTTTTTTTATATTTATA**ATGAAATGCC**TATTACTTACATTTCGACTTAAGAAAATGCTAATAATATTATTCGCTTTCGAATTTATGATGGTAAAAATGTTATTTGCTTTTGTATACTTAAGTATACCGTTTGACCCATTATCTTTATTAGTCTTCCTGGCGGTTGCGGCTGGAGAGGCAAGAATGGGATTAAGTCTATTAGTATCTATTTTACGACAGAGAGGAAAAGACAAAATTACTAGATACTCTCTGAATTCTTTTGAGGGCTATTAAATCACGAGAAATTCAAAAAGCGAAGAATGGTTTCGACCCATTTATTGGGTACACATACCCTCTTGTGTTTGCCACAAATGTCTAATATTCCATTTATTCCAGTTAGAGGTATAATTATGATTACTATAGGATTGTGAGTTTTATTAGTATTTTTTACTGTTATAGTCCCTTTCGAAGATATTAAGAATGGGGGAGCAAGCTCCTCATCCCAAAATGGTTTATCTAATTGACCTTGGAACTAAATTCAAGTTTTAAATAATCCAGATGTACGTAGTAAAATAAATATTTTAATGGAACTGGGTGACTTTTAGTCTAAACAAATCTTTGATAGCTAAAATTTTTATAGTGTGGTCTTGAAGCGGCCGAGGTGAAGATATCTTCTCAAAGAAAATGGGGTAGTCTAATTATGAAGATATTGGATTGTAGCTCTAATGATGAGAATAAATCTCTCCCATGAGGGCTGGTAGCATATAAAGTGTATTGAATTGCAGATTCAAGGGTGTTTA**GAAACCCAGCC**CTAATAGTTATTTGGTTTTGGTTAGATAAAGATTTGATGTTAAAACCAGCATAAAAGAATTATAATTATTAAGATAAACACAATATATAAATTATCATATTTAGTGGTGGATTTTTATAAATAAAAGTGAGAGCTTGTTTAAGTTAATATCAATGAGGGGCAAATATTAGGTGCCAGCAGCTGCGGTTATACCTATTCTTCTCCTTTATTTTTGGGCTAAATATAAAATTTTATTTTCAGATTAAAAGATCTTAGTTAGGTTAAATTTTTAAGATATTTTTTTATATTCTGAAAAACTAAATTATCAAAATTATTATAAAAACCAAGATTAGGTACCTTATTATTAATAAGAGTAAATATGAATACCCAGAGTAGTAATATTTTGAAACTCAAAAGACCTGGCGGTGTTAAGATTTCCTACCAGAGGAGTGTGCTAAGTAATAGACAATCCGCTAATTAATAAACCCTTAAATGATTTAGTCAGTGTACGGCCGCTATGGTCAATTTTATTAGAATATCGAAATTGGCACTATATTTCTACTTTAACTAGTTAAAACAGGTCAATGTGCTGCTGATGTAAGGGGTTGAAGGTGCGCTATTTTAATTAAGATAAAAGATTATATGTTTTAACACATATATGAATTCAGATTTGGTAGTAACAAAATATAAAATACGTTTTTGTGAACATAGGTAAAGCTTCGTGTACACATCGCCCGTCGCTCTTTTTAAAAGAAAAGAGAAGTCGTAACATGGTAATCTTAATAGAAATTGAGGTTTAAAAGGTAGTATAAAAAAAATTATATAAAGCTGTTAACTTTGTGATGTGCTCTACCAAGCACCCTTTTAGTTTAGAGGATAAGTTATTTATAAACTATAGGGCTCATGACCCTAAAAAGATTTTGATCTCCTCTAATTGAAAACAAAAGGA**ATGTTTGGTAGAGGACTGATATATG**GTATCAAATTACAGAAAGCCTGTAGACCAGTATCTCGTGAAACAGTATTATTCCATGATGAGAACTTAATAATATTAATAGTGATAGCTTGTTTGGTGGGGGGAAGTCTATTTATGTTATGATCTATTAAGTTCTCATCATTTGACTTTGTAGATCATAAGTGGTTAGAGGTAGGGTGGACCCTATTACCTGTTTTTATTTTAATAAGTCTAGCCTTACCCTCCTTAGAATTATTATATTATATGGATAGACCATCATCAATTACACCTTTTGCTACTTTAAAGGCAATAGGCCGCCAATGATATTGATCATATGAGATAAGCGTTTTAAGCTCAGATGAGGTATCCAGTTTTAGATACGATTCATATATGTTACAAGAGGGTAGTGAAAAAGATATAGATACTGATTTAGCAGGCTTCCGTTTATTGGAAGTGGATAATCCTATTTTCTTACCCCGAGCTGAATATATTCGGTTATTAGTAACCGGGGGGGATGTAATTCATAGTTTTTGTATTCCAAGGCTGGGTATAAAGGTAGATGCTGTACCTGGTCGTTTAAATCAAACATATTTTTTCCCGTTAAATCTAGGCAGATATTATGGCCAATGTTCCGAGATTTGTGGAGCTAACCATAGTTTTATGCCTATTAAAGTAGAAGTTATACCTACAAATATATTTTTAAATTTCTTTAATTAAAGATTCTTAGGTTATATAGACCATAAGCCTTCAAAGCTTATAGAGAAAAGTTTTTTCAGATTCTGATTGAAAGAATATTTTAAAAGTGG
