## Supplementary for "Mitochondrial Genome-Based Phylogeny of Turbellarians and Evidence for Accelerated Mitochondrial Evolution in Symbiotic Species": annotationEurylepta cornuta.docx

ATGGAGAAAAGGGGCTTGTAGCCTGGACTAAGAAGTGGTTTTACAGTAGAAATCATAAGAACATAGGGACTATTTACTTAATATTAGGT**GTGTGGTCAGGTTTAATAGGG**ACAGCGTTAAGTTTCATTATTCGTTTAGAATTAGGGCAGCCTGGGAGTCTTCTACAAAACTCTCATGTTTATAATACTGTTATTACTGCCCATGGTTTAATAATGATCTTTTTTTTCGTAATGCCGGTTATGATAGGGGGATTTGGTAATTGGTTAATTCCTATTTATGTTGGGGCACAAGATATGAACTTTCCCCGACTTAACAATTTAAGGCTTTGGTTATTAATTCCAGCTATAGCCTTGATGCTAAGGTCTTTAGTTATAGGTGAAGGGGTGGGGGGAGGATGGACTATTTATCCTCCTTTATCAAGTAATATGGCACACAGGAGGAGAAGAGTGGATATTGCTATTTTTGCACTCCATTTAGCAGGTGCTAGTTCTATTTTAGGTTCAATAAAATTTATCAGCACAGTAGGTAAATCTAATAAGGGCTCAGTTAACTGAGCCCGGCATTCTTTATTTATCTGGGCTATGATTATAACAGCCTATATGTTAGTATTGTCCTTACCTGTGTTGGCGGGGGGTATTACAATGCTACTAACCGATCGCAAATTTAATACTAGGTTTTTTGACCCGGGGGGAGGTGGTGATCCTATTCTCTTCCAACATATTTTCTGGTTTTTTGGTCACCCTGAGGTTTATATTTTAATCCTCCCTGGGTTTGGAATGGTTTCTCAAATTGTTACTTTTTACAGAGGTAAGGATGAAGCTTTTGGACATATAGGTATGTTATACGCTATCGTAGGTATTGGTGTATTGGGGTTCGTGGTTTGAGCTCATCACATGTACACGGTAGGTTTAGATGTGGATAGACGAGGTTACTTCACAGGAGCTACTATGGTAATTGCAGTACCTACGGGTATTAAGATTTTTAGATGACTCTCTACTTACCACGGACGCCCATTACGGGACTTATTTGAGGTACCTGCAGCTCTTTGGTCTTTAGGGTTTATTTTTTTATTTACTATTGGAGGTCTAACAGGGGTTGTGTTAGCTAGGGCTAGTTTGGATGTATGTCTACATGATACTTACTATGTGGTAGCTCATTTTCATTATGTACTCTCAATGGGGGCAGTGTTTACTATATTTGCTGGGTTAGTACATTGGTGACCTTTATTTACAGGTGTGGCTTTAAAGTCCACCCTTGTAGTTGCTCATTTTTGGGTTATGTTTATAGGGGTTAACGTAACATTTTTCCCCCAACATTTTTTGGGTCTTAGGGGGATGCCCCGTCGGTATAGGGATTACCCTGATGGATATGCTTATTGAAATGCTGTGTCTAGGTATGGGTCTTTACTCTCTGTTGTAGGCGTTACCTTTTTTATAGTAACAATCTGAGAGGCTATAGTGGCAGAACGAAAGATCTTATTTGTAGTTGGGCCCAGAACCCACCCCGAGTGGACTAATTCTTCATTACCTTACGGAGAGCATAGTTTAGATTTAAGTTTAGCTCATGTAGTTAAGGTAAAATAAGGAATTCTCTTTAGTTTAAAAAGAATAGCTGCCTTCCAAGCAGTTGGTCCATTTTGGAAGAGAAAAGAAAAAAGAGTTTTGAAGGGAAAAAATAGATTAGGTAGTAAAAGAGTTTAAAATTTGGTTTAATTCTCTTCCTTTTGCATCATGATCCAGTAAATTAGGGGTTTATACCATAATA**CGAAACTATTTGGTAAACTTTTCTTTAGCCAGGGGCTTATAAATAAAAAGGTTTTTTTTAAAAAAGGTTTAGTAGTATAGATAATAT**TATCGTATATAGTAATAGCTTATAAAGAGGGAAGCTTTCGAGTTAATATGAACGGGTAAAATCAGTTCTTTATTTATAATGTATTTGTATTTTAATACGAAAATTAAAATTTGAGGTGAGAAGTTTCTAAAATAAAAAAGTGTTTTTACTTTAGTCCAGGCTTACAAGCATTTTTATAGCTCTTCTTAGTTAATCGCAAAAAATTAAACTATGCTGGAATTAGTAAATGACTAAATATACAAAAACTCGAGATTAGCCCCAACTGTTTAACAAAAACATTTCCTTCTTGTATATTTTGAAGGTAAAGCCTGCTCAGTGGGTATATTAAACAGCCGCGGTATTTAAGCTGCGCGAAGGTAGCATAATTAATTGTCTTTTAATTGGTGACTTGTTTGAATGGCTTAATGAGGGCTAATCTTTTTGTATGCTTAAATGAACTTATTTTCTAGGGTCAAAAAACCTAGATTAATAAAAAAGAAGAGAAGACCCTGATGAGTTTTACAATTTAATATTGTTTTAATGGGGCCTTATAGTTTATTTCATACTATATTTGATCCTACCTTGAGGAAAAAGGGATAAATTACCACAGGGATAACAGGACAAAATTTACTAAGAGTGCATATCTAAGTAAGAAAATGTTACCTCGATGTTGAATCAAAGTTAACTTTTAGGCGCAGAAGTTTTTGGAGTAGGTCTGTTCGACCTTTAAAACTTTACGTGATTTGAGTTAAGACCGGCGTGAGCCAGGTTGGTTTCTATCTTTTTAAGTGGGAGGTTTGTACGAAAGGAACACTTTATATTATTTGTTACACATACATTTGGTAAGTAATAATTTAAGATGGCAGATAAATGTATCAGGTTTAGGACCTGAAGATGAAAATTTCTCTTTTAGGGTCATCCTTGGGGAAGCTATCCTTTCTAGATAGTAATAGTATATTTTATACACTCTAAATATGAGA**ATGGTAAGTTGGGGGCTTTTATTAGGTATAAATTATCTTTGTATGTACGCAAAAAAAACATTAAAGTTGGGGGGTATGATTTTG**ATATTAAGGATAAATATTAGAGTTATAGTAAGATTAAATTATTTATCTTGGCTTTCTTTGATTTTTTTTATCATCTATATAGGGGGTTTGTTAGTACTTTTTTTTTATCTGTCTTCTTTAAACTACAAACCTATATTTTCTTTAAGTAAGTTAAGCTTAACCAGGAGTATGTTGGTGAAGATTAACAGGGGGTTGGTAATACTTTTAGGCAGGCTTAGTTATACTTTTTCTGAGGGTAGGTTCAGATTTATTACAAGCAAATTCAAAAATTTTAGAAAAAGGCTATTCAATGAAGAGGAGTTTAATAGCCTGATATTGGTGGGGGTGATGTTATTATTAGTTTTATGGCTAGTAACAAAGCTAACCTACAAAACACGAGGGGCTTTACGTCCAACTTTTTAGACTTAAGACTCTTAAGATTTAGATCATTCAATTTTGAAAGTTGAAGAAGGGGGTATCCCCAGGAGTCTATGAAGTCACATTTGCATTATGTTAGTAAATTATTTTTACTACTAGGGGTTGTGGTGTTTGGGTTGACTGGTAAAAGATTTATGTTTAGATGAAAAGTAGCAGATTTAGGAAGGTGTAGCCTTGATTTACTAGTATTAGTGGATTGGGTATCACTAACTTTTTTTTTTACTTTATCTATTATAGTAAGATGTGTTTTAAAGTTTTCAAGAATATATATGGAAAAAGAAAAGTTCTTTTTACGATTCCATTTTCTAGTCTATAGGTTTGTTTTTTCTATGTTTTGTTTAATTTTCTTTCCTCACTTTTTTTTTCTTTTAGTTGGATGGGATGGATTAGGGATTACAAGCTTTCTATTAGTGATTTATTATTTAAGTAGGTCTTCATGGGCGGCTGGAATGAAGACCTACTTAATAAATCGGTTGGGGGATGGGTTGTTTTTAGTAGCGTTAGGTTTACTTTTAATACAGGGACATTGGGACATAAACGGTATAGCTAACAACCAAGTGTATTGAGGGTTTATTCTTGTGCTGGTTTTAGGTTGTTTTACCAAGAGGGCTCAATTTCCATTTTCTAGTTGGCTCCCAGCTGCAATGGCAGCACCTACTCCTGTCTCCTCTTTAGTCCATTCCTCTACTCTAGTTACCGCAGGTATATACTTATTAATACGCTTTAGAAGTATTATACCTAATTGGATGTTTTTTTTTATAGGGGTAATGGGTTTATGGACTTTATACGGAGCTAGGTTAGCGGCTTGTTTTGAGTGTGATGCAAAGAAGATAGTGGCGTACTCTACTTTAAGCCAATTAGGGTTTATGGTTGTTTCCATCTCTAGAGGGTTGGAGAAATTTGCGTTTTTTCATTTAATAACCCACGCAATGTTTAAGGCGATGATGTTTATTTGCGTGGGTTATTTCATGGTAAAAAAAAACCATTTTCAAGATTTTCGGAGATTGTCGGGTATGTGAAAGTGTAGGCCTTTTGTTAGAATTAGTTTGGTGGTTAGAGGGCTTAGCTTAATAGGGTTTCCTTTTATGTCGGGATTTTTTTCAAAGGAATTGGTCTTAGAGTCTAATTTATTCTGTGTTAATGAAGTATTCTTTAATCTTTTACTTCTTTCATTACCTCTAACTTCTTTTTATGCAAGCCGACTAATTTTTCAGTTATTTAAAGGTAAAGGCTATAAAGTGTTTAGGCGATGTAAAGACAAAAGCACGGTTCTAATGTCTCTTATACCTTTATACCTAGGTAGAATACTAATAGGTAGGCTTGGGAGGTCTTTTTTTTTAAGTAATAGATTTTTATACCCAAGTTCTTATTTTAAGCTTTTTGTAAAATTTTTAATTTTTAGGGGGGTTAGTTGAGCATGAATTGGAGTCAAAAAAAACAGGCCTTCATTTAAATGATTCTATTCAGAGATTCTTTATACGTCTCCCTTTAATGGTAAATTTTGAGTGGATGGAATAAGTGATCTTGGAGAGGACATAAACTATCTTTTCGACCAAGGCAGGTTAAGAAAATTTATATTCAAAATGGATTTCGGAATATTAAAGGCTGGGGGATGGGTTAGAAAAATGTACTATTATTTTCTAAGTCCAAGGGTTGGTTTATATATAATTTTAGGTTTTTTGGTTTTAAGCCTAAGTTGGTATATCTAAGAAGTTAGTATAATAAATTATTTTTGGTTGTCGGCCAATAGAAGAGAAATTTCTCACTTCTTATATTTAAGGTGGCAGATAAATGCATTAGATTTAAGCTCTAAAGATGAGGTATTCCCTTAAAAGTTTTGTATAAGAGGGTGTCCCCGCACGTAAAATTTTGGATTTTAAAGTAAACGTAGTTTTCTTATATTTAGGGGTAGTTTATATAGAATTATAGCTTTGGGAGCTATGGGACGATTTTTTCGCTCTTAAAAATGGCTTGGAAGCTTTGTAGAAGCTGTAGATTCTTGATCTAAAGAAGGGGGTTCCTCCCAAGCCATTGAAGTCTATAGGAATAAGTATATTTGGTTTAAGAGGA**GTGTTAATAGCTTTGATTATGGTTTGGGCTATTT**GAAGTAGACGAAAGAATTATATTAATTCTCGTGAAAAGAGATCCCCCTTCGAGTGTGGATTTGATCCAAAGGACAAGGCTCGGATCCCTTTTTCATTACGGTTTTTTATTATAATAATTCTTTTTATTATTTTCGATGTAGAATTATCCCTACTGCTACAACTTCCTCTTCAGTTTGAGAAAGGATATTTTAAGAGCCGGGTGTTGATTTCTCTTTTTATCTGAATACTTTTATTAGGTACATTAGAAGAGTGGCGGCGGGGGGTTTTAAGTTGAAAGGACTAGGAAGTATCTTTAAAAGTATGAAAAGCTGCTAACTTTTTATAGAGTATTTAGTATTCATATTTCTATGATATCCTTTCTCAAACTGAATAAAAAAAAGTGGTTAAGAAAATTATGTCTTTTACTAGTTATGTTGCTAGGAGTATTAATTTGTGTTAGGAGGTCTAATTTATTTATTTTATGGTTTGGGCTTGAGCTCAATATGTTTGGAGTTATACCTTTTATAGTTTTAAACTCCAAACCAAAAAATCAAAAAAAGGACTTAAAAGTAGGTATATACTATTTTATAATCCAAGTGATTGGAAGAATTCTTTTTTCTTGGGGGAGGGTAGTGGGAAGTAGAAGCATTTTGGGTCTTAGGGGTTTACTTATTAAGATGGGGGTAGTTCCCTTTTTTTGATGAGTTCCTTCAGTGCTTAGACGAGTAGACTGAGGTAGATTCTTACTATTAAGCACACTACAGAAGATTCCCTCTGTCTTACTAATACGTATCAGGTTTGATTTATCCTTTAAAGTCTGCTTGTTTATATGCCTAGTAGGCCTAGTGGTAAGGGTTATTGGGATAAAATTTTCTTCAGGTAAATTTAAGCTTCTGTTGGCGTGGTCTTCAGTAGGAAAAATGAGCTTGGTGGTAATATTAGCTACCCTTTATTTTAATTTAGGTAGTTTATACTTTACCTGTTATGCTATTAGAGTTCTAGGATTTATTTTATGCATCTCCTCCCCAAAAAGAGGAGATATAAAAAACAAAGAAAATAAAGGGGGAGTTAATAGTCTTATAAAAATTTCATTTTTTTTACTCATTTTTTCTGGCCTTCCACCTTTACTTGGTTTCATAACAAAGGTTGTGTTGTTTAGGGGGTTAAGTATTTGTGAAAGAGATATGATGATAAAGCAGATTAGAGTTATTGACAATAATAACTATATGGTTGATTACCCTATCTACAAAGTTATTGGTAGGTGAAGTCTAAGAATTGTTATAAGCTTTATTTTAATTTTACAAATAATAGCTTATATTAAGGTATTTATAAGGGTGTACACCTCTATTTCTATTAAATTACAAGGAAGCTCACGAAGTACAAATTCAAACTATTTGGAAGCAATGTTAGTAGTCGGCCTATTAAGAACACTTCCGCTGGTACTGTTGTAGTAAAGGGTAGTATAGTTATGTACGTAAAGCTGTTAACTTTAAGGCATGGGGTTACTTTCTCATTCCTTTACAAAGTATGAGGACTGCGATAGCAGGTTACTGTGATATAGTAAACTTTGGGGCCACCCCCATGCTAAATGATTTTTGATTTGTTTTCTAGGGCAGATTACTGCTATTCTAAATTAGGTTTGTTTAAAAATGTAATTTGGGTTTTAGGGTTTATTTGATTATTTTGGAAAATAAACTTATTTTTTTATAGTTTAAACCGGTTAAGCCAATTTTTTTCTATACTTTTATCTTGGCATTACGAAAAAATTAAGTCTTCTAATATCTCAAATATTAAGGGAAGATATATAATTATAGGAGGGAGATTTTTTTTAATTTTAGGAAGAAATTTATGGGGGTTATTCCCTTATGTGTTTGGGCTAACAACCCAAATGGTTTTAACTTTTTCGGCATCATGGATTATATGACTAAGAATTGTGTGGTCTAGTGGAGAATATTCTATCTTAAAGTTCTTTTCGAGTTTTACTCCTTCTGGCTCACCAGGATATTTGGCCCCTGTACTTTCTATTATTGAGATTATTAGCAATGTTATTCGGCCTTTAACGCTTTGTTTACGGCTTAGGATAAAAATTACTACTGGGCATGTCTTTTTAAATTTAATGGGAGTAGGAAGAAATAGTATTTTGTTATTGGTAATCTTAATGTGAGGCTATCTACTGTTTGAAATGGGTATAGGATTTATACAAGGTTTTGTGTTTAGTTTATTAACCTCACAGTATTTAGGAGAGC**ATGTTTAAATTTTTAATTTTCTTAATA**CCTTGGGCTTTCATTTTTATAAAAGTAATTATAGCCATCCCTTTCCTAACTTTGTTAGAGCGTAAGGTAATAGGTTATATACAAAATCGAAAGGGACCAAATAAGGTTAGGTATTTAGGCCTTCTACAACCAATTAGGGATGGGGGAAAGTTAATTTTTAAGGAGTTAGGGACCCCAAGTTATGCTAAAATAATTTTATTTTGGTTTAGCCCTATTGCTTCTTTAATGCTAATGATTGTGGGATGGGGTATTTTCCCAAGATATTTTAGTAGGTATAGCCTCAACTTAGGGGTGGTATTTTTTTTGTGTGTAGCAAGACTTCAGGTCTATACTCTTTTGGGGTCTGGATGGGGAAGAAATTCAAAGTACGCTCTTTTGGGGTCTGTTCGGGGGGCTGCTCAAACCATATCCTATGAGGTGGCGCTAATATTAGTTATCTTTTTACCTTGTTGTCTTGAAAAAAGGTATTCTTTTTTAAAATATTTGTACAAGTGTTATCCTTATTTTTTTATTTTATTTCCTATTTTGTTAATATGAGTTATTTCTAGGCTAGCAGAAACAAACCGAGCCCCTTTCGATTTTGCAGAGGGGGAAAGGGAGCTTGTGTCAGGGTTTAATGTTGAATACTCAGCTTTTGAATTTGCGTGCTTATTTTTATCTGAGTATGGAAAAATTTTATTAATGAGGTTTTTAACTAGGTTATTATTTTTTAAAAGAGGTTTGTTTATCTACATTTTAATAGGGTTAAGAGTAACTTTTTGGTTTGTGTGGGTTCGTAGAGCTTTACCTCGTTTTCGGTATGATCTGCTAATGAATTTAGCCTGAAAGGTATTTTTACCTATTTCACTATTTTTTATTTTTTTAGTAATAACTATATGGCTTTAGAAGCAAAATTTGCGTCGGTTTTGTAACCCGGAGGTTGATGTCAATCCAAAGCCAAAAAAGATAGTTTAATAAGAATTTTGCTTTGTGGTAGCAAGGGTGTACAAGTTGCTCTTTTTAT**ATGTATAAAGGAAATTTAA**ATTTAGTTTTTTCTCCCCGTCGGGATGAGGCAGTAATAAATTTTACAAATAGGTTGCTCTACGACCTACCCTCCCCCAAGAAAATTTCTTTTTGGTGGGGTTTTGGGTCTCTGTTGGGTTTGTTTTTAATTATTCAACTAGCAACAGGGCTGTTTTTAGCTATGCATTATGTATCAGACTTGAAAGTTTCATTCCAATCTGTTGATTCATTAAATCGTGAGGTGGGTTTGGGTTGGTTGATACGAGCTATTCATGCTAAAGGAGCCTCGGCTTATTTTTTGTTTTTATACCTTCATATCGGTCGAGGGGTTTATTATAGTTCTTACCTACTAAAAAAGAGCTGAAATACAGGGGTTTTAATTTTGTTTTTATCTATGGCAGCTGCCTTTTTAGGTTATGTACTTCCTTGGGGACAAATGTCTTATTGGGGGGCTACTGTTATAACTAATTTTTTTTCAACCGTCCCCTATGTAGGGCTTGAGTTGGTTGAGTGAATATGAGGAGGATTCGCGGTTGGTTATCCTACTCTAACTCGGTTTTTTGCATTACATTTTTTGATACCTTTTGCAATATTAGGGTTGGTTTTAGTACACATAATATTTTTACATGAGGGAGGGTCTAATAACCCTTTAGGACTAGCTTCAAATGGGGATAAGATCCCCTTCCATCCTTATTTTTCTATAAAGGATATAATGGGGTATAGGGTAGTAATACTATTTTTTTTGTGTGTGGTGGTATACTCCCCAGATTTTTTTACAGATCCTGAAAATTTTATAGAGGCAAAAGCATTAGTTACTCCTGTACATATAATGCCTGAGTGATATTTTTTACCTGCTTATGCTATTTTACGTTCTATACCTAAAAAGTTAGGAGGGGTAATTGCCTTAGTCCTAAGGGTAGCAATATTATTTATATTCCCTTTAATATTTTGTAAAGATAAGCGGGGGAGTCAGTATTCTGTAGCTTTACAGGTTTCTTTTTGGGTATGACTAAGAAGAATTTTAGTCTTACTTTGGATAGGTTCTCGACCTGTTGAGGATCCTTATATAATGATAGGACAGTTAAGGACAATAGTTTATTTTAGATATTTTGTAATCTTACCTGCTATTAGTGTAGATTTGTTTAAGGGTA**ATGGTTAGTAGGTTAATTTTT**CTTTTGTTTATTCTAAAAGGAGGGTTATTAATTGTTCGGGCAAAGAGGATTTTAATCATGCTATTTGCTTTCGAAGTTTTAATATTAAATATTTTTTTAATATACTCATTTAGCAACAGCCTTAGTGACATAAGGTCCTGCTTAATTTTTCTTTCTGTAGTAGCAGGGGAGGCTAGGTTAGGCCTTTCTTTACTAGTTTCAGTAGTGCGTAGATATGGAAAGGATAACAAAACCAAAAAAAACATAAA**ATGTGAAGGATATTAGGTTTAGGGGCTAGACTA**CTATTTATAAAAGAGATAAGAGTAATAACGTTTGTGGTTAGGGGGTTGGTAATTCTCATCACAAAGTATTTAGTAATTAGGTTTTCTCATTTTATCTCTTTTCACTCTTATTTTGTATTTTTTGATCAGATATCAAGATTCTTATGTTTGTTAAGGGTATGGATAACTCTTTTAATGGGGTTAGCTATGCTAAAAGCAGAAAATAAGAAGTTTTTATACTTTAAATTCGGAGCTTTAAATTTAATTCTTTTATTTGCTTTTACAGTTAAAGACTATATTAGATTTTTTATATTTTTTGAGTTATCTTTAATTCCTACTCTTATTATAATTCTAGGTTGAGGAGTTCAACCCGAACGGGTTCGGGCTGGAAGGTACTTAATAATATACACTGTAGTAGGGGCTATACCCTTATTACTATGCATAATATACAGTTACTATAAAAGGGGGTGTACAAAAATTGTTTATAGAAGTATAGGAAATTTATTTACCATAACCTTAGAGTATAGCTGGATAGTTATTTTTTGGGTGTTTGGGTTTTTAATAAAGCTCCCAGTATATGGGGCGCACTTGTGACTCCCTAAGGCTCATGTTGAGGCCCCAGTTAGAGGTTCAATGGTTCTTGCTGGGGTTTTACTTAAGTTAGGAGCCTATGGCTTAATCCGAGTTAAAAAGTTTCTCGCAATTTCTGGGTGTTTTTGGAGTAATGTTGCATTTATTTGGTTTTTGTTTTCTATGTGTATGGTTGGGTTTGTTTGTTTTCGGCAATCAGACCTTAAGTCTCTAGTAGCTTATTCTTCAATTGCCCATATGTCTTTAGTGGTGTTAGGAGTTTTTTCTTTTGATTATTTAGGATATATAGGAGTTTTAAGAATGCTGGTAGCTCATGGAGTGTGTTCATCAGGTCTTTTTTTTTGTGTTCAGTGTTTTTATGAACTAAGAGGTACTCGCAGGGTAGTCCTAAACCGAGGGTATTTAAATTTATCCCCCATTTTATGTTTTTTTTGGTTTTTGCTTTGTGTGGGTAAAGCCTCTGCTCCCCCCACCCTAAACTTGTTTAGAGAATTTTTATTAATTTCTTCTATTTTAAACTACAGATTAATTAGTAGAGTGTTTCTTTTTTTGTCAGTGTTTTTAGGAGGATTGTTTAGTATATATTTATACACCTTCACTTCACATGGAAAGTGAAAAAAAATTAATAATTATTGAGGGGGAATTAGCCTACGGCATTGTTTAATTTTGTTTCTTCATGTTATTCCTGTTTATTTTTTAATTTTTAGGGGTAATTACTTACTACCTAGGGTGTAAAACCTTAAGGTCCAAAGAATTGGTTGATGGTTTCGGCCCATTAGTTGAGGTTTGTCCTCCCTTAAGAAGTGCCACATATGTCGAATATTCCATTTTTAATGGTGAGGTTTGTTTTGCTAGTATTTTTTTTGGTGTGGATTCTAAATGTATGGTTTAATTCAAAAAAAACACCAAAAAAAACAGGAGGTTTATATAAGAAAGGTAGAAATAAATGGCCTCTATAGTTTTTGATAGCTTAATTGTAAGAGTGTGGCCCTGAAGAGGCTAAGGTGAAAATATTCTCAAGCAAGGTGGTCTAAATTAAGGCGTCAGGTTGTAGCCCTGGAAATAACATTAGTTTCTTGCAGGGCTAATAGCATAAATATTATTGCATTGAATTGCAGATT**CAAAGGTGTAAAAACTTAGCCCTAAGGCTA**TTTGGTTTTGGTAGCATAAAGGACAATAGGCCTAACTAGCAGTAGATAATTTATTTAAATATTCTAGAACAAATTTTTGTTTAGATTTCGTTAGTAGTGCATATTTCTCTATAAGTGAAAGCTTGATTAGGAGGTCTATTAAAATAAGGGGCGAATACATAGTGCCAGCAGCTGCGGTTATACTTGATCTTTTCCTTTAAGCTAGGGTAAAATGGAAATAATTTGATATTAAGAAAATAAATATTAGCATATGGAGGAATAAGCTTATGAATCAAACATACTTTTTATAAATAAATTCCTAAAGTTAGATTAAAAACTAAGATTAGGCACCTTATTATTCTAAACATAAATTTTTAGACCCATAGTAGTAGAAGTCTCGAAATTAAAAAGACCTGGCGGCAGCCTTACTCACCGGGGGAGCGTGACGCATAATAGATAACCCGCGTTATAATTAACCTTTTTTACTTAGTTAGTGTACGGCCGTTTTCAGCCTAGCTCTGCAGAGCATAGACTTAGGTATTACTGTTGTAGTTAAGACAGGTCATTGTGCTGCTTATAAAAAGGTGTTAGTTTCGCTATTTTGATTGGAAAAAGGGAAAAGCTTTTGTAATAAGGCTTAGAATTAAGACTCGGCAGTAATGGGGAACTATTTAGTCCCCATGAAGGTAAATTTTTCTAAACACACATACTTACTATTTTAATTATGACTACTTGAACGAGAAAGCATAGCTTAGTAGCCGTAATTGGCAGATGGATTGTGTGTTTAGTAATTATCCTATCGCAAACAACAAAGTTACTTAAACGAGAAAGCAAAGCTTAGTGATTTTGTTTTAAGGGAGGGTAGTTCGGGGTTGTGTACATACCGCCCGTCATTCTTCTCTAAGAGAGAAGAAAAGTCGTAACATGGTAACCCCAGCTGAAGCTGGGGTTAAGATACTTAGAAGTTTTACAAACTTTGGGTTGTTATACCCAATAAAGATTATCCTAAGTTAACTAAAATACTCTAATTTTGAAT**ATGCGGGGCTGAATGCC**AGACTTGATAAGTGCTAGACCATGACCCAGAGCAGTTGTATTCGGGAGATTAAATCTTGCAGGGGGGTCTATTGTATTGTGGCACTATTTTAGGCTTGATATAGTAAAGTGGGGGTTACTCGCTTTAAGACTATTAGTCGTTTTTTGATGGTGGGATATGTACCGGGAAACCGGGGGTGGGAGTCATACTCGTGGGATTAGCTTTGCTCTCCGGTATGGTATGGCCTTGTTTATTAGGTCAGAAATATTATTTTTTTCTGCGTTTTTCTGAAGGTATTTTCATGGTTATTGACACCCTGAAGATGAAGGAGGGGATTGGACAGGTTCTGAGTATGGGTCTGTTATTCTAGATCCTTATGGGTTACCCTTTTTGAAAACTTTAATTTTACTAGTTTCTGGAATTTTTGTGACCATGTCCCATCATGGCCAAATTAGACGATGTGGCTTTACACGGATTCTAATTTTTATTACTATGATGATGGGAGTTTACTTTATGCTTATGCAAGGGGTAGAATATGAGTCTTCAGGATTTTCAGCTAATAGAAGTGGATATGGAACACTATTTTTTATTCTAACTGGGTTTCATGGAATGCATGTGAGTGTAGGAGTGGTGCTACTAAGATTTGCGTTCTTTCGACATGGTCGGGGAGCTATGAATGAATCAAAGCATGTGTTTCACGAAGCAGCTGCCTGGTACTGACATTTCGTAGATGTTGTTTGGCTAGGTTTATATTTCTTTATTTATTGATATGGAATGGACGGTTCTGGGCCTAGTGTCTAAAGCAGATTTTTTCACCGAGGATACGATATGGACATTCCTGGATCTAGTGGGTGATATAGACTGGTTTCAGGTGGATGTAGACGTAGACATGGAGTGGGACGATCTTTTTACAAGACACTTTGAAGAATGACTAGAGCTTGACAAGTGGATCTGATAATCGGGTAGATGTTAGGTTCGGGGTTGTCACTTTAGACTTTTATGGATTATATTTTTTTAGAACTTTAGTTTATAATCTGGAATTTTTGTGGCCATGCCCCATCATGGCCAAATTAGAAGATGTGGTTTACACATCTTCTAATTTTTATCACTATAATGATGGGAGTTTACTTTATGCTTATGCGGAAAATAGAGTATAAATCTTCAGGATTTTTAGTTGATAAAGGGGTGTGTAAACCTATTTTTTAAGCTGTTTAGTAGTATGTCTACTATTTAATTACTAAGCTTAAAAAATGGCTTACATGCTAGAGGATAGGTTAAATTAGACCGGAGGGCTCATGACCCTAAAGTGGTAAATCCTCCTCTAAT**ATGAAAAATCAAT**TTGGTACTGGGGTTATTTATGGCTTAAGTTTTCAAAAAGGGGTTAGCCCTGTAATGAGGGAGACTATAAAGTTTCATGATGAAAATATGTTAGTATTAATAGTTATAGCATGTTTAGTAGCAGGAAGTATTATTTTAGTATGGTGTAATAGGTTCAGTTCTTTAGAATTTATAGATAGAAAGGTTATGGAAATACTTTGGACTTTATTACCTATATTTATTTTAGTTACTTTGGCTCTACCTTCTTTAGAACTCTTATATTTTTTAGATTGCCCAAAGGATATAGATGGTGTTGCTACTATTCGGGCGACAGGAATGCAGTGGTATTGAAATTATGATATAACTTATGAACTTACTGGAGCAAAGTTTAATTTCGACTCTTATATATTGGACTCCAATAGAAAAGAAAAGGGATATCGCTTGTTGGAGGTAGACAATCCGGCTTTTGCCCCTCGGGGGGAGGTGGTAAAGTTGGTGGTTAGAGGGGGGGATGTAATTCATAGGTTTGCTGTCCCTTCTTTAGGGGTGAAGTTGGATGGAGTTCCAGGACGTATGAAAGAAGGAGGATTTATAGCATTAAAGCTAGGTAGTTACTATGGGCAGTGTTCCGAAATATGCGGAGCTAACCATAGGTTTATGCCCATTAAAATTGAGGTTATCCCTATAAAGGCCTTTGCTGAAGGGTTTAAATCTGCTATTAAACCTAATATTTTATTGATAATAAAACAAAGAAGTCATTCCAGATATACATTAAAAGGGTAGGAGATCTAAAAGGTTTACGGGCAGATAGCTTTAATTGATTCTGACCCCTAAACTTTTTATACATTTTGAAAAAGTTACCAGGGTTTAGGGGAATTTATTTCGGCAGAAACCTAGTAACTTTTTTAATAGGTAAGGGTACCCCCTTACACTTTTTTTAATTAAATAAATTCCCCTAAACCCCATCTATAAATCTGCTATTAAAACCTAATATTTTATTGATAATAAAACAAAGAAGTCATTCCAGAAACTAGTAAAATTAAAGTTTTCAAAAAGGGTAACCCATAAGGATCTAGAATAACAGACCCATACTCAGAACCTGTCCAATCCCCTCCTTCATCTTCAGGGTGTCAATAACCATGAAAATACCTTCAGAAAAACGCAGAAAAAATAATATTTCTGACCTTAATATTTTATTGATAATAAAACAAAGAAGTCATTCCAGATACATATTAAAAGGGTAGAAGGTCTAGCCCCCTACTACTTCGTAGTAGGGTTGCTTCAATATATTTTTTAGTAGTAATATACTTAGCATGCTTTTTTTAATTACTACTACTTAGTAGTTTATATAAAACTTTGGATTTGCATTTCAACAAAGGATTACATCCTTAAGTAGACCCCACAAACGTGGAAAAAAGGAGACTATATACTTATTACAAAGGAAGTTTGCGAAATTTAAATTTCCTCCCCACAAACGTGGAAAAAAGGAGACTATATACTTATTACAAAGGAAGTTGAGCCTGAAGTTCTGTATGAACACTTCATTTACGGTGAAATAGTGAGATTATCTCCAGGCTTATTACTAATAAAGTAAGGCATCGTCTTATTTTTTTTTAAGGGAGCTAGTTTATTTGAACGTAAACTTTCAAAGTTTAAGGAGAATGTATATTCGTTTCCTGTGAAATTCTTATTAGGCCCCATATTAAGGAGCAATAGAAAAGAGGGATCTAGATTTT
