## Supplementary for "Mitochondrial Genome-Based Phylogeny of Turbellarians and Evidence for Accelerated Mitochondrial Evolution in Symbiotic Species": annotationImogine fafai MF993335.docx

ATGAGTAAGAAGGGCTGGATAGCCCGATGACTTTATTCAACCAATCATAAGGATATTGGGACATTATATTTAATATTTGGTGTGTGATCTGGTTTAGTAGGAACGGCATTTAGTTTTTTAATTCGTTCTGAATTAGCTCAGCCAGGTACTTTGCTTAAAGATCCACATTTATATAAAGGGATTATAACAGCCCATGGACTCATTATGATTTTTTTCTTTGTAATGCCTGTATTAATAGGAGGTTTTGGTAATTGATTAATTCCTCTATATTTAACAAGTCCAGATATGGCATTTCCTCGTCTTAATAATATGAGGTTTTGACTATTACCTCCCTCGTTTTTTTTACTATTAGGTTCTTTTTTAGTAGAAACGGGAGTGGGAACCGGCTGAACAGTTTATCCCCCCTTATCCGGAAATATCGCCCAAAGAGGTCCAAGGGTAGATATGGCTATTTTCTCTTTACATCTAGCTGGGGTTAGGTCAATTTTAGGATCTATTAAATTTATTACTACTATAGTAAAAGCTAAGGTTCAGGTATCATGAGGACAGATACCATTATTTTTATGGGCTGTTATGGTTACAGCTTATATGTTGGTTTTATCTTTACCAGTATTAGCTGGAGGTTTAACTATGTTGTTAACGGATCGTAAATTTAATACCACTTTTTTCGACCCAGGGGGAGGGGGAGATCCGATTTTATTTCAACATATGTTTTGGTTTTTTGGTCACCCTGAAGTTTATATATTAATACTGCCAGGGTTTGGTATGATTTCTCAAATAGTAACTTTTTATAGTGGGAAGGATAGAGCCTTTGGACATATGGGGATGGTATATGCTATTCTTGGTATAGGGTTTTTAGGTTTTATTGTGTGAGCTCATCATATGTACACTGTGGGGTTAGACATTGATACCCGTGCTTATTTTACAGGAGCTACGATGATAATTGCAGTTCCTACTGGTATAAAGATATTTAGATGATTAGCTACCTTTTATGGGCGTCCTTTAGGACAGCAAGCTGATAAAGTTGGACCGTTATGAGCTACTGGGTTTATATTTTTATTTACTTTAGGTGGTTTAACTGGGGTTGTATTGGCAAGTGCAAGGTTAGATATAGCTTTACATGACACTTATTACGTTGTAGCTCATTTTCATTATGTTTTGTCTATGGGAGCAGTATTTTCTATATTTGCTGGTTTGGTCCATTGATGACCTTTATTTACAGGGACAGTACTAAATGGTAAGATGGCTATAGCTCAATTTTGAGTTTTATTTACTGGTGTAAATTTAACGTTCTTTCCCCAACATTTTTTAGGATTGAGTGGAATGCCACGTCGATATATGGATTATCCTGATGGTTTCTCTTATTGAAATAGAGTTTCTTCTTTAGGTTCTTTAATTTCTGTTGTTGGTGTAGTATTTTTCGCTGTTATTATTTGGGAAGCTTTAGTTAGTGAACGACTAGTTATATATATAGTTTCTCCAAATTCTCAAGCTGAGTGAATTAATTCAGATAATTACCCTGTGTCTTTTCATACTAGAGAGTCGGTAAGAAAGGTTTTTATCAAAGAATCTTAGATTGGGTTCTCTTTAGTTTAATAATAGAATATTTGCCTTCCAAGTAAAGGGTCCTGACAAAGGAAGAGAAAAGTAAATATTAATTAATATATTTGTGTTTTTATTAATTTTCGTAAGTTTTATAATTATTTTGTATATTGGGAAAACCTCCTTTTAGTATCACGGTCTAATAAATAAATAAATATATTTTTATATCCGAAATTTGGTGATTATATATTTTTTAGTTTTGAACATAATTATTTAGGGGTCCTATTAATTAAAAAAAATTTAAGAATAAAGATTATAGATCGGACCAAATTATATCTGATTTGAAAGAAATATTGCGAGATAAACCCAAAGGTTAATTTTTTGGGTTTGGTTAAAGATTAAACTATAGGTTAAT**TGTTTAAGATGAG**AAGTGTCTAAAAATAAAAAGTGTTACAACTTTAATATAATAGTTTGTTTTCTTGTAAAACTTTTTTAATCATAATTCCTTAATATATTGATAGAATAAGTATATAAAATGGGTCATATTAACTCGATACTTTTACAACTGTTTATCAAAAACATTTCCTTTAGAATATATTTAAAGGTATTACCTGCTCAATGTAGTATTATAAATAGCTGCGGTATCCTAACTGCACTAAGGTAGCATAATTAATTGTCTTTTAAATGGTGACTTGTTTGAATGGTTTAATGAGAAAAGGATTTTGTTTGGTCCAAAAAGAATTTATATTTTAGGGTTAAAATACCTAAATTGTAAAGAAAGACGAGAAGACCCTGTTGAGTTTTACATTTATATGTTAAATGTTTAAATGGGAAATTTAATTCCTTTATGGAATTTAATAGATCCTAAAAAAGTTTGGAAAAAGGATAAAATTACCACAGGGATAACAGGACTATATTTTCTGAGAGTTCTAATTGAAGAAAGTGTTTGTTACCTCGATGTTGAATCGAAATAACTTTTTGGTGCAGTAGTCAAATAAGTAGGTCTGTTCGACCTTTAAAATTTCACGTGATTTGAGTTAAGACCGGCGTGAGCCAGGTTGGTTTCTATCTTCTTATGATTGTTTTTTGTACGAAAGGAATTGAACGATTGATATTTTCAATTTTGGAATGGCAGATTATATGTGATAGATTTAGGTTCTATAGATGGAGATTTTCTCCTTCCAAATATTTTGTAATCTAATGTTGGTGTATCCCAGCACTTTAAGCTTTCTACTTAAAAGAGCTCTTTAGGGGCACATTAGAT**ATGTCC**TGTATACTATGATTTTCTCTAATGTTGATTAATTATCTTTGTTTATTTTCATTAAATTCTTTAGTGTTAGGATTATTGATTCTTTTTTGTAGGTTTTTTATAAGCTTACTTATAAGTATATATTCTTTATCTTGGTATTCATTAATATTTTTTATAGTATATATTGGAGGTTTATTAGTTCTTTTTATTTATATTTCTTCATTAAAATTCAAACCAGTTTTTTACTTTTCTCGTACTGGGGGATTCTTAAAAATTTTATTAAAGATAAAAGTAGTATTTTTTTTGGTGATAAGTTTAACTCAAATCACGTGAAATTTTAAAGGTTTTTTTTGAACAGAATTGGATACAAAAAAATTTAGATTTAAATTATTTAAAGAAGTTGAGATATTATTTCTGATAAAAGTAGGATTAGTTTTATTAGTAGTTTTATGGGTTATAACAAAGTTATCTTTTCGAAATCGTAGCGCCCTGCGCCCCTTTTTTAGGTAAATATTGGTACACTTCAATTTTGATACT**GTGATTAA**GCACAATGTTAAGTATGGTTTTAGTTATACTTGTTCCAAACTTTTCTTTTTACTGAGGATTATATTTGCGATAATCTTCCTTTGTTTAGAGAAAGGAATAATGGTAAAATTAAAAATTTTGGATGTTGGCAGGCTTAAATTAGATTTATGTTTTTATGTAGATTGAATTAGAATTCTGTTTTTTACAGTTTTATGTATTATAGTCTCATGTGTTTTAAAGTTTTCTTGCATTTATATGGAAAAAGATATATTTCAAGTCCGTTTTACTTGATTAGTTTTAAGATTTGTATTTTCCATGTTTTGTTTAATCTTTTTCCCCCATTTCTTTTTCTTGTTGGTGGGGTGAGATGGATTAGGAATTACTAGTTTTTTATTGGTTATTTATTATTTGAGAGATTCATCTTGGTCTGCAGGTATGAAGACTTATTTAATTAATCGGGTGGGGGATAGGTTTTTTATATTGGGTTTAGTATGTTTTATGGCTAAAGGGTATTGGGATATAAAAAGTATTACTAACAATAATATTTTATCTTTGATAATAGTTTTAGGATGTTTTACAAAGAGGGCACAGTTTCCCTTTTCCAGTTGATTACCAGCAGCCATGGCTGCTCCCACTCCCGTTTCTGCATTAGTTCATTCTTCCACTTTGGTAACTGCTGGAATTTATTTAATGGTACGTTTTTGTAAAATGTTTCCAGATTGATTATTTATAATAGTGGGTGTTAGAGGGATGTGAACTATGTATTCAGCTAGTCTTTCTGCTTGTTCTGAATTTGATTCAAAGAAGATAGTAGCCTATTCTACTTTAAGTCAATTAGGTTTAATGGCAGTATCCATTTCTTTAAAATTGCCATTGATTGCATTTTTCCATTTAATTACTCATGCAATGTTTAAGGCTTTAATATTCATTTGTGTTGGGTATTTAATTAAAAAAAGAGGACATTTTCAGGATTTACGGAGGTTAAAAGGTTTATGGCGAACTAGTCCTTTGTTAGGAATTACTTTAATAGTGAGAAAATTATCATTATTAGGTTTCCCATTTTTGGCAGGGTTTTTCTCAAAGGAGTTAATATTGGAGAAAAACATATTAGTTATAAATAGGGTATTTCACACTTTATTACTTCTTTCCCTCCCCCTAACTTCATATTATAGTACTCGTTTAGTCTTTAATATATTGAAAGGGGTAAACTATAATAGTGTATCTTGTAGTAAAGATAGAAAAGTGTTGTTATTTTCTTTACTTCCTTTATATCTTGGAAGAATATTTGTTGGGAGTATAATTTACCCTCATTTTTATAGATTAAGTTCTATTTATCCATGTCACATGATGAAGTTTTTTGTTTTTTTATTTATTCTTAGGGGTGCTTTTCTGAGATGAAAAGATGTTAAGTTCAAAAAAAACTCTTTTATTTGGTTTAGTAGCTCTATATCCTTTTTAGTTCCTTTTAATGGAGGATTTTGAACCAAAAATTTTGTAAATTTGGGGAAAGATTATTTTTGTTTAATTGATCAAGGTATTCTATCTTCCATAATGGGAAGTTTGGATAAGGGGGTAAATTCATCCGGAAGATGATTAGTAAGCTCTTTTGAGTATTTTAGAACACCACGGGCTAAGTATTATGTTAGGTCAGTTGTAGCATTTTCTATGTTTAGAGGTAGAGTTGCTTAACATAGACTTTCAAGAATTTGTGGTTCGTTTTGTTTTGAAAGCTTAAAGAGGGATAATAATACTTCCCGGAAGTCTTTTCAAAAGAGTTAGTATATTTTAATTATCATTGGTTGTCGGCCTTTAGAAGGGGTATACCCCACTTTTT**GTGAAT**AAACAGTTTTTAAGTTTAAATAAATTTTCTAAGGGGGATAAAACTACCCCATTTCATTTGGTAGAGGTTAGCCCGTGGCCAATATATGCTTCTTTGAGCGCTTTAGGTATAACTTTTGGAGGTGTATATTGATGGCATGTAGGAGAATCTTGAATTTTTTTAACAGGTTTATCATTTAAAATTTTGGTAGCCTTTTCTTGATTTTCGGATGTTATAAAGGAAAAAATGGCTGGTTTTCATAATAGAGTAGTTATGTTCGGATTCCGGTTTGGTATGATATTGTTTATTTTGTCGGAGGTTTTATTTTTTTTCTCTTTCTTTTGGGCTTATTTTCATAATTGTTGAGGTCCCCAAGCTGAGCTTGGATTCATTTGGCCTCCCTATGGGTTTAAAACTATTGTTATAGATCCTTTTTCAATACCTTTATTGAAAACGGTTGTACTCTTATCTTCAGGGGCTAGGGTTACTTGAGCTCATCATTCTTTAGTTAGTCAAGATTATGTGAATGGTTTAATTGGTTTAGGGGTAACTGTGTTTTTAGGAGGCTATTTTCTATTTTTACAGGGTAACGAGTATTTTCTTAGAGAATTTTCTATTAATAGAACTATTTATGGTACGGTATTTTTTATGCTTACTGGATTTCACGGATTTCATGTCACGATAGGGACTATATTATTATTTGTTTGTTTTTTACGTCATTATTTTAGTCATTTTTCTACCCATCAACATGTCGGGTTTGAGGCTTCAGCTTGATATTGACATTTTGTGGATGTAGTATGATTATTTTTATATTTTTTTATTTATTGATATGGTTATAATTTATAATTATATTTTTAGAGTGGCAGAGCATATGCGCTGGATTTAAGCTCCAGAGATGAGATTTATTTCTCCTTTAAAGGATATGGGGCAGTGTAAGAGGCACATAAGTTTTTGGTACTTAAGGAAAACGTAGTTTCTCCGTAATCTGAAAGCGAAATGGGAGCAGCAGATTCTTACTCTGTAGGTGGACTATTAAGTCCTCAGATTATTTTATAAAGAAAGAAAATATAAGGGAATTTCATTTCTAAAATTCCCTTATATTTTCTTTCTTTATAAAATAATGTAAGGGGACCCCCTTACATTATTTTATAAAGAAAGAAAATATAAGGGAATTTTAGAAATAGTTTAAGTAAAATATTAACTTTGGGAGTTAATGATCAAAAGTATCTTTTGTTTCTAAATTGTTTCCT**ATGTCTTTT**AGTTCGGTTGTATTTTGGAATATAGTTGTGATTTTGATTGGGGCGTGGGTGATATGAACAAGGCGAAAGAATTTTTTAAATTCCCGAGAAAAGTCTAGTCCTTTTGAATGTGGTTTTGACCCAAAGGATAAGGCACGGATTCCCTTCTCTCTCCGGTTTTTTCTAATTGTTATACTATTTTTGATATTTGATGTAGAATTATCTTTATTATTACAGCTTCCTTATCAGTTAGATTTTGACCATTTTAAGGGTCGGGTTGGGATAGTTATATTTGTCTGAATTTTATTACTAGGTACTTTGGAAGAATGACGACGAGGTATATTAAATTGAAAGGATTAGAGATATATTTTTATAAAAAAAGAGGTGTTCTCTTTGGATTTAAGAGGCTGCTAACTTCTTTTAGAGTAATTAAGAATTGTTCACACCTCTATTAATTTCTTTAGA**GTGCGGATAC**TTCCTAATATATGGAGAAGAACACTTTTTTTGAGATTTTGTTTTTCTTTTGGGCTTATTATATCCCTTATTAGAAATAAAGTGTTTTTGGTTTGATTAGGACTTGAATTAAAAATGTTTGGAATTATTCCTTTTATTAATTCAAGTCCTAATCAAACCAAAAACACTTTATATCTTAGCAAAGAAGAAATAAAAGTTTCCTTTTTTTACTTCTTTGTTCAGGTAATAGGTAGGTTACTTCTCGCATGGGGAAGAGTATTAGGGGGGTGATTTATTATTAGGATAATTGGATTGATAATAAAGATAGGAGCTGCCCCATTCTTTTGGTGGGTCCCGCCTATAGTTTCCCGATTAGATTGGTTTTCTATAGGTGTTATTAGAACAGTACAAAAGATTCCAGGAATCTTTCTGTTTCGGTTAATATTTGACTTGGATTTAGGTATATGTATGTTATTAAGAATAGTTGGGTTTAGAGTATCGGTTGTAGGAATAAAATTTTCCTCTAAAAATTTAAAGCAGTTGATAGGATGGTCTTCTATTAGAAATATGAGTATATTATTTGTTTTAATAGTACTTAAAAAAAACTTTGGGTTGATTTATTACCTTTTTTATAGTATTCTTGTATTAATATTTTGTTTTATATTGAAAGTGTATTCTATGAGAAATATTAGAAGAGCCTTTATAAAAGGTAAAAAAAGGGTTTATTCTCTTTTGGTTGGAGGTTTACTGTTAATATTTTCTGGATTACCCCCATTGGTTAGGTTTTTATTAAAGGTTTACTTTTTAAGGGGGTTTTATTTCTTTGATTGCATAAAAATGTTATTAGACTTGGAATTTAATGGAGTTTCTATTTCCTTTTTTCATTTATTAGCTAATGCTTTGAAAAGATGAAGTGTGGTTACTTTTTTTCTATTATTAATAATTTTTCAAGCCGTTGGTTATGTAAAGATCTTTATAAAGATCTTTACAAGGAAGTCCTCACGCGTTAATGGCAGAGTGAAGCTCTTAAAAAGAAAGATTAAGATATTTTTTAAGTCAGTTTTTCTATTATATTTATTGAGTATTTTACTAATTTTAGTATAAATGCTTTAAAGTCAAAATAGAGACGTAAGTCTTGTAAACTTAAAATTGAAGAAAATTCTTCTAGGGCAATAATATTTCTTCCGATAGTTTATTCAAAACGTTAGCTTTGCACGCTGGATATGGGTTAAAGCCCCTCGGGTGATATAAGGTGAATGTGAGGACTGCGAATGCAGGTCACTGTGATATAGTGAACTTTGGGTTTAATAACCTCCACATTAACTTTG**GTGTAT**GATTTATTTACTAGTGCAGATTTTTGTTATTCTAAATTATCTTTATTAAATATATTTGTTTGGGTTTTGGGTTTGGTATGAATTTTTTGGAATATAAACCTTTTTTTCTTTAGGAGAGGGCGGCTAAAAAGGTTTATATCTTATTTATTAAACTGGCATTATGATAATGTAAAGACTTATTTTTTATATAAGGTTAAGGGATCTTATATAGTAATATCAGGTGCATTCTTTATAATTTTGGGATGTAATGTTTGGGGATTATTTCCTTATGTTTTTGGTATTACCACTCAAATGGTATTAACTTTTTCTATGTCATTAATTATATGATTATGTATAGTGGTTTCTAGATTGGAATATTCTTTTTTAGGGTTTTTATCCCATTTAACTCCACAGGGGTCTCCTGGTTATTTGGCCTCAATATTGAATTTAATTGAACTTGTTAGAAATATTATTCGTCCTTTAACTTTAGCTTTACGTTTAAGGATAAAAATTACTACTGGTCACGTATTTATAAGATTAATGGGTACCAGGGGTAGCATATGTCTTTTTTCTTTTAAGTTATTTTGATTGTTTTTTGTATTCTTAATGATGGGGTATCTTCTTTTTGAAGTAGGTATATGTTTTATTCAAGGATTTGTCTTTAGTTTATTAAGCGTGCAATATTTAGGTGAGCATACTTAAAAGAATAGTAGTTTAATTAGAACACTGCATTGTGATTGCAGGGGTATACTATTAGTATCTTTCTTCTGTACCGTTCTAAATTAAATATTAAA**TCTTTTCGACGAGA**CCGTTCGGTTTTAAAGACAATTAAAGGTGGGGTATATGATTTACCCTCCCCCAAGAAAATTTCTTATTGATGAGGGTTTGGTTCATTATTGGGGTTATTTTTAGTTATACAAATATTAACAGGTTTATTTTTGGCTATGCATTATGTGTCCGATTTAAAAGTAGCTTTCCAATCTGTGGATTCGATTAGTCGTGAGATAGAATTTGGGTGATTAATACGTAGAATACATGCTAAAGGTGCTTCGGCATTTTTCCTATTTTTATACTTACATATAGGTCGTGGGATTTATTATGGTTCTTTTTTATACAGTCATACTTGGAATACAGGGGTAGTAATTTATATATTAGCCATGGCCACTGCTTTTTTAGGTTATGTATTGCCATGAGGTCAAATGTCTTATTGAGGGGCTACAGTAATAACTAACTTTTTTTCTACTGTACCTTATATAGGTAAAGATTTAGTTCAATGAATATGAGGAGGATTTGCTGTAGGCTATCCGACTTTAACTCGTTTCTTTTCTTTACATTATGTGCTTCCTTTTGTTATTGCTGCTTTTGTATTGATTCATTTAGTGTTTTTACATGATACTGGATCGAAAAATCCTTTAGGACTGAGTTCTGCTGGGGATAAGGTTCCATTTCATCCTTATTTTACTATTAAGGATATCTTTGGATTTAGTATAGTATTATTTTTATTTTTTTCTGTTTCTGTTTATTCTCCAGATTATTTTGGGGATCCAGAAAATTTTATAGAGGCTAATCCATTAGTTACTCCTATACATATTCAACCTGAATGATATTTTTTACCAGCTTATGCTATTTTACGGGCCATTCCAAAAAAGTTAGGTGGGGTGGTTGCTTTGTTAATGTCAATATTAGTACTGTTTATTTTTCCTTTGGTGTCTTCTTTTTGTAATCGAGGATATTTTTATTCTCATTTACATCAAGTATTATTCTGATTTTGAGTGGCGGATATACTAGTTTTACTATGAATAGGGGCACGTCCTGTTGAGGAGCCATATGTTAGTATAGGGGCAATAAGAACTATAGTATATTTTATGTTCTTTTTACTAACCCCATTATTAAGATATTTTTATAAAAATATATCCCAGGAGAAGGGTAATAAAGGTGACATATCTTAGATATAATAAACATTTATACTATATGTATATATAGCTAAATGTTATAAGAGAAAAAGATTTGAAAAAAGAGAGTAACTTAATATTACTTTTGAAGGGGTATACCCCTCAAATCTAGTCAGGGGATTAATATGTGCAACACTTTCATTAAGTTTATTTGGTAGGGTAAATTCATTAATA**ATGGGGTTAG**TAATAATTATGTTACTTTTTACTCCTTATTTAATGGGTATTCACTCCCATTTTATATTTTTTCATTCTTATTGGGCATGTTTAGATAAAATGTCAACTTTTCTTATAATTTTAACTCTTTGAATAACTTTATTAATGAGTTTAGCTATGAGAGATGCAAAGCAAAAGCGCAAGTTGTTATTGTGTTTTTCTTCGTTGAAATTTATATTGGTTTGTGCTTTTATAAGTAAAAGACTTTTAGGTTTTTATATCTTTTTTGAGCTATCCTTAATTCCCACTTTAATAATAATTTTGGGGTGAGGTGTTCAACCAGAACGGATTCGGGCTGGTAGATATTTAATGTTATACACTTTAGTTGGCTCTCTTCCTTTATTGGGTACTATTTTATATATGGACTATCATTGTGGCAGGGTAAAGATATTTATGCCATTTATAGATATATTCTTTTTTTCGAAGTGGGAATATAGATTTTTTTGTTTATTATGAATAATGGCTTTTTTAATAAAGTTACCTATATATGGGGTTCATTTGTGATTACCAAAGGCACATGTGGAGGCTCCAGTAGCCGGTTCTATGGTTTTAGCTGGGGTTCTATTAAAGTTAGGGGCCTATGGTTTAGTTCGTTCTCTTAAGTTTATTCAACTAAATTTATGTTTTTGGTCTGACTTCTTTTTTATATGGAGTTTATTTACTATGTGCCTTGTGGGTTTTATGTGTTTTCGACAATGTGATCTTAAGTCATTAGTGGCTTATTCTTCTGTTGCCCATATGTCTCTTATTTTGGCTGTATGTTTTTCTAGAGATATTATAGGTATTAAAGGGGTGATGGGAATGTTAGTTTCACATGGTTTATGTTCTTCCGGCCTGTTTTTTGGAGTTCAATGTCTTTATGAAAACAGTGGTTCTCGTAGATTATATTTGAATCGGGGTATAATTAGGTTATCACCCCTATTTTGTTTTTTTTGGTTTCTGCTTTGTGTTGGTAATGCTTCAGCTCCACCAAGCTTAAATTTATTAAGGGAATTCTTCCTTATATCAAGAATTGTTAGATATGGTGGTCCCATATCCGCAATTTTTTGTGGTTTGTCTGTATTTATGGGAGGATTATTTAGAATTTATTTATATGTACTGATTTGCCATGGAAAGTGGTCGGTTTTGAAAAAATTTTGAATGCCTTTTAGTTTACGTCATTATTTAATTCTTTTATTACATGTTATTCCGCTATATGGTTTATTGTTTTTAAGAAAATATATTTTTTATTTTTAACCATAATTTTGCGGAATCTGTAGTTTATGTGAAAACTTATCGTTTACACCGATATAAAAGATGGTTGTTATCTCGGATTTATATAAGTTTCTTT**ATGGTGATTCC**ATGGTTATTAACCTTTGTGAATATTCTGGTTTCTATACCCTTTTTAACTTTATTGGAGCGAAAGGTCTTAGGATATATACAGTCACGTAAGGGACCCAATAAGGTTAGCTTCCTGGGATTATTACAACCAATAAGAGATGGAGGTAAGTTAGTTTTAAAGGAATTGGGAACCCCAAAAATGGCCAAAATAATATTATTTTGGGTTAGTCCAATATTATCCTTTTTTTTAATGATTATGGCATGATCTGTATTTCCCTCCCCCTATCAGTTTTTTTCTTTTAAATTAGGAGTAGTTTTTTTTCTTTGTATTGCTAGATTACAGGTATATACTCTATTAGGATCTGGGTGGGGGTCTAATTCTAAGTATGCTTTATTAGGTTCGGTACGGGGAGCAGCACAAACTATTTCTTATGAAGTTTCTTTAATTTTTATAATATTATTTCCTTGTAGATTAGAATATAGGTATAAATTTTCATCTTTTTTAGATAAGGCTTTTAGATATATATTAATTTTATTTCCCATATTTTTGATATGGTTTACATCTTGTCTTGCTGAAACCAACCGTGCCCCTTTTGATTTTGCTGAAGGGGAGAGAGAATTGGTTTCAGGATTTAAAGTAGAATTTGCTGCTTTTGAATTTGCTTGTTTATTTTTATCTGAGTATGGAAAAATCTTATTAATAAGGTTTTTAACAGCGATATTCTTTTTACCTACAAAGTACCTCTTTTTTATAGTTTGTGGTTCTATGTTTTCGTTTTGTTTCGTGTGAGCACGAGGATGTTTACCCCGTTTTCGGTATGATTTTTTAATGAGTATGGCATGAAAGATTTTTTTACCTAGATGTTTGTTTGTATTTTTCATATTGTTGGCATAGATAATGGAGAATATTTTTTATAAAGTTTGATTAACTTTATTCTCTTGAAAAGGTTGGATATCTTTTATATTATTTTTTTATTAAAATGTTTATTACTTTTATTTCGGTTGAAGAAA**ATGCTCATAATATTATTTTCTTTTG**AATTTATGGTTGTAAAAGTATTATTCGTGTTTGTTTTTTTAAGAATCCCTTTTGATCCTATTAGATTGTTAGTATTCTTAGCGGTAGTAGCTGGGGAGGCGAGGTTGGGATTAACTTTATTAGTATCGGTTTTACGCCAAAGGGGTAATGATAATATACTTCCTATATCTTTTAATTCTTTTGAAGGGTTTTAAAGAAACATCCTCGTGAGAAATTCAAATAGAAGATTGGTTTCGGCCCAATAGTTGAGTGTCTTGAATGCTCTCTCATGTTTGCCTCAAATGTCTAATATCCCATTTTTATTAGTTAGTTTTACTATCTTTTGTTTTTTTGTTATTTGAATTTTGAATACTTGATTTAATATTTTTATATCCAAGGGTTCAAATAGTTCGGGTATTGGTAAGAATAAGGTGTTTACCTTATGGCCATTAAATTAAATATTTCTCCGATAGCTAAAGTTTTATAGTGTGGTCTTGAAGCGTCCGAGTTGATGTTATATCTCGGGGAAAGTAAAGTAGTCTAAATTAAAGGCATTAGGTTGTAGCCCTAAAGATGAGGAGGATCTCCTTTGCAAATGGGGAGTTAATAGCATAAGTTATAATTGCGTTGAATTGCAGATTCAAAGGTGTTTTTACTTAACTCTA**GATTAATTTG**GTTTTGGTTAGATAAAGAATTAGCATTGAAACTATCATAAAAATTTTGATATAGATTTTCTTTTTCAAAAAGAATTTTATAATTAGTGGTGAATTTTTCTAATAAAAATGAAAGTTTGTTTAAGTTTATGCTGAGAAGGGGCTAATAGTAGGTGCCAGCAGCTGCGGTTATACCTATTCTTATCTTTATTTTTGGGTAAAATACAAATTAAATATTTGAGAGAAACTGAAATTAATCCGGTTAAATGGGTTTAAATCAAGTATTCAAAATTCAAATAGTAGTATTGTTAAAGTTTATAGAAAGACCAAGATTAGGTACCTTGTTATTATAAACTGTAAATAATAACACCCAGAGTAGTAAAATAATTAAAACTCAAAAGACCTGGCGGTGGATGTTCTTCCTACCAGAGGAGGGTGCTAAGTAATAGATAATCCGCAAATTAATAAACCTTTATTTTTTAAGTCAGTGTACGGCCGCTTTCAGATAATTTTCATAGAATTATTATAATTATCTTTGTTTTCTAATTTTGGAACTAGCTAAGACAGGTCAATGTGCTGCTGATATTAAGGGTTAATGGTGCGCTATTTTTATTTTAGGTAAAGGATTCTCATCTGTAAGGTTGGGATGAATTGAGATTTGGTAGTAATGAGGTTATTTAATATGTACTCATGAATTAAGGTAAAGCTCTGTGTACACATCGCCCGTCGTTCTTTTTGAAAAAAAAGAGAAGTCGTAACATGGTAGTCCTAATAGAAATTGGGGCTAAATAAAGGATAGTATAAGGTTTATTATATAAAGCTGTTAACTTTGTGATGTGCATAAAGCACTCTTTTAGATAATTTAGAAGATAAGTTATTACTTAAACTGTGGGGCTCATGACCCCAAAAAGATTATTTATCTCTTCTA**ATGAAAAATA**ATAGTGGTTTTGGAAGAGGTTTTATATATGGTCTCAAATTACAAAAAGGTACAAGACCTGTAATGCGGGAATCTATTCTTTTCCATGATGAAAATTTAGTAGTACTAGCAGTAATTGCTTGTTTAATTGGTGGATGTATAACTTTGTTATGATGAACTAATTACAGATCTTCTGACTTTGTAGATCATAAGTGGCTAGAGGTTGGTTGAACATTATTACCTGTATTTATTTTGGTGGGGTTAGCCCTACCATCCTTAGAATTATTATATTATATGGATAGACCTTCTTCAATAACTCCTTTTGCAACTTTGAAGGCAATAGGGCGTCAATGGTATTGATCTTATGAGATTAGGGTATCAACCACTGAGATGGTTGATAGAGTATCTTATGATTCTTATATGTTACCTGAAGGTCAAGAAAAAGAAAAAGATGATGATTTAGCGGGGTTCCGTCTGTTAGAAGTGGATAATCCTATATTTTTACCACGGGGGGAATATATTCGTTTATTAGTCACTGGTGGGGATGTAATACATAGTTTTTGTGTTCCTACTTTAGGTATAAAGGTTGATGCTGTTCCAGGCCGGTTAAACCAAACCTATTTTTTCCCACTTAGATTGGGAAGATTTTACGGGCAATGTTCTGAACTTTGTGGAGCTAACCATAGGTTTATGCCAATAAATTTAGAAATTATCCCAACCAAAGTTTTTTTGAAGTTCTTTGGGTAAAATCTTTAGGTTATTAAAGACCGTTAGCCTTCAAAGTTAGAAGAGAAAATAGCCATTTTCAAGATTTGTGTTTTCAGAATACTTAAAAAAGTTTATTTCACGTGGTTGA
