## Supplementary for "Mitochondrial Genome-Based Phylogeny of Turbellarians and Evidence for Accelerated Mitochondrial Evolution in Symbiotic Species": annotationImogine stellae MF993336.docx

GTGGCACGATGACTTTATTCAACAAATCATAAGGATATTGGTACTTTATATTTAATTTTTGGGATTTGAGCAGGATTGGTAGGAACTGCATTTAGATTTATAATCCGTTCAGAACTTGCCCAACCCGGAAGAATTTTACATGATCCTCATTTATATAAAGGTATTATAACAGCCCATGGTCTAATAATGATTTTTTTCTTTGTAATGCCAGTTCTTATAGGAGGGTTTGGTAATTGATTGATACCCTTATATTTAACTAGCCCAGATATGGCTTTCCCCCGATTAAATAAAATGAGATTTTGATTATTACCTCCTTCATTTTTTCTATTACTCGGGTCTTTTTTGATTGAAACAGGAGTTGGAACTGGATGGACAGTTTATCCTCCTTTATCAGGTAATATAGCCCAAAGTGGTCCAAGAGTTGATATGGCCATATTTTCTCTTCATCTTGCTGGTGTTAGTTCTATTTTAGGATCTATTAATTTTATTACTACTATAGTAAAAGCTAAGATACAAGTATCATGAGGACAAATACCTTTATTTTTATGAGCAGTAATGGTGACGGCCTATATGTTAGTCTTAGCATTACCTGTTTTAGCAGGAGGTTTAACGATGCTTTTAACGGATCGTAAATTTAATACTACTTTTTTTGATCCTGGGGGTGGTGGAGACCCTATTTTATTCCAACATATGTTTTGATTCTTTGGTCATCCCGAAGTGTATATTTTAATTTTACCTGGGTTTGGGATGATTTCCCAAATAGTAACTTTTTATAGAGGAAAGGATAGTGCTTTTGGTCATATGGGAATGGTTTATGCTATTCTTGGGATAGGATTTTTAGGATTTATTGTTTGAGCTCATCATATGTATACTGTTGGTCTAGATATTGATACTCGTGCATATTTTACAGGAGCAACGATGATCATTGCAGTTCCAACTGGTATTAAGATTTTTAGGTGATTAGCCACATTTTATGGTCGTCCACTAGGACAACAAGTAGATAATCTTGGACCCCTCTGGGCTACTGGGTTTATCTTCTTGTTTACTTTAGGTGGTTTAACTGGAGTTGTATTAGCCAGTGCTAGTTTAGATATCTCTCTACATGATACATATTATGTGGTTGCACACTTTCATTATGTTCTTTCTATGGGGGCAGTGTTTTCTATATTTGCAGGGTTGGTTCATTGGTGACCTTTATTCACAGGTACTGTATTAAATACTAAGATGGCTATAGCTCAATTTTGGGTATTATTTACTGGGGTTAATCTAACCTTCTTTCCACAACATTTTTTAGGTTTAAGAGGTATGCCTCGTCGTTATATGGATTATCCAGATGGATTTGCTTATTGAAATAGTGTAAGTTCCTTTGGATCTTTAATTTCTGTGGTAGGTGTTATTTTTTTTATTGTTATTGTTTGAGAAGCATTAGTTGTAGAACGATTAGTAGTATATATAGTGTCTCCTAGTTCTCAAAGAGAATGAATAAATGCAGATAATTATCCTGTACCATTTCATACAAGAGAGTCTGTAAGAAAGGTATATGTTGAAGATTATTAAAATTCTTTTTAGTTTATGATAGAATATTTGCCTTCCAAGCAAAAGGTCCTTATTATAGGAAAAGAAAGTAAGATTTAATTAATATTTATTTGAGATTTATATATTATTTTGTAAATTTTATTATTAATAAAATATATTGGGTTTATTATCTTCCTTTTTGTATCACGGTCCAATAAATTAATGAAAGGATTTTCTAATCCGAAACCCCATGATTATATTTTCTCTAGTTTTGAACATAATTGTTGAGAGGACCTCTTAATAAAATAGAATTTAAGAATAAAGATTATAGATCGGATGGGGTTATATCTGATTGAAAAGGATATTATATCTAATTATATTTAAGGGTTAATTTCTTAAATATTTTTAAAGAATTAATTATTGTTTGAAATATTGTTTAAGGTGAGAAGTTCCTTTTAATTTAACAGTGTTATAACTTTATTAAAATATTAATTATTTCTTAAAATAAAAAACTTTTCTTATATGAATTATAATGCCATAATGATGGAATTAGTATAATAAATAAGTCATATTAACTCGGTATTTTTACAACTGTTTATCAAAAACATTTCCTTTAGAATATATTTAAAGGTATTGCCTGCTCAATGTATAAATAAATAGCTGCGGTATCTTAACTGCACTAAGGTAGCATAATTAATTGTCTTTTAAATGGTGACTTGTTTGAATGGTTTTATGAGAAAAAAACTTTCTTTGACTTAAATAGAACTTATATTTTAAGGTTAAAATACCTAATTTATAAAGAAAGACGAGAAGACCCTGTTGAGTTTTACGAATATATTAAAATCGTTAAAATGGGAAATTTAATTTTTTAATAAAATTTAATAGATCCTAAATCTATAAAGATTTGGAAAAAGGATAAAATTACCACAGGGATAACAGGACTATATCTTCTGAGAGTTCTTATTGAAGAAGGTGTTTGTTACCTCGATGTTGAATCGAAATAACTTTTTAGTGCAGCAGCTAAATAAGTGGGTCTGTTCGACCTTTAAAATTTCACGTGATTTGAGTTAAGACCGGCGTGAGCCAGGTTGGTTTCTATCTTCTTATGACTGTTTTTTGTACGAAAGGAATTGAACGGTTAATTATAATTAAACTTCAGGATGGCAGATAATATGCGATAGATTTAGGTTCTATAGATGAAGAATAACCTTCTCGTGAAGACTAGTGTTGGTGTATTATATGCACGTTAAGCTTTCTACTTAAAAGAGTTCTTAGGAGCACATTAGAAAAATCTCATGTATTATATGATTTTCTTTG**ATGTTAGTAAA**ATATATATGTTTATTCTCTTTAAAATCATTAGTTTTGGGTATATTAATTTTAGTGGGTAGATTTTTAATTAGTTTATTAGTAAGAATTTATTCATTATCCTGATATTCTTTAATCTTTTTTATTGTTTATATAGGTGGTTTATTGGTTCTTTTTATATACATTTCTTCTTTAAAATTTAATCCAGTTTTTTATTTTTCACGGATAAGGGGATTTGGGAATATTTTTTTAAAGTTGAACATTCTGTTTTTTTTATTAGTAAGATTAACTCAAATCACGTGAAATTTTAAAAGATTCTCTTGGAATAATGTGGATAAAAATAAATTTAGATTTAATTTATTTAAAGAGGTAGAAGTAATATTTTTAATAAATGTTGGGATTTTATTACTTATTGTTTTATGGGTTATAACCAAGTTATCTTTTCAAAAACGAAGGGCACTCCGTCCCTTTTTTTAAAAAGGGGATTTTTATTGGGACGTTTCAATCTCGAAAATGTATTTAAGTACAAAGAAAAGTATGGATTTAGTTATACTTGTTCC**AATATTTTTTTCGCCTTAAGGATACTATTTGGATCTATATTACTATTTACTAAGAACTCATATCTATTAACACTAAGG**ATATTAGATTTAGGATCTATTAAATTAGATGTTTGTTTTTTGGTTGACTGAATTAGGCTTCTATTTTTTACTGTTTTATGTATAATAGTATCTTGTGTTTTGAAGTTCTCTTGTATTTATATGGAGAAAGACACTTTTAAAATACGATTCACTTGGATAGTTTTAAGTTTTGTATTTTCAATGTTCTGTTTGATTTTCCTTCCCCATTTTTTTTTCCTTTTGGTGGGGTGAGATGGGTTAGGAATCACTAGATTTTTATTGGTTATTTATTACTTAAGAGATTCCTCTTGGGCTGCTGGTATGAAGACTTATTTAATTAATCGAGTGGGGGATAGGTTTTTTATTATTGGCCTTGTATTGTTTTTATATAAAGGATATTGAGATATAAAAAGTTTAGGTAGAAAAAAATTATTGGCTATTATTGTAGTTTTGGGATGTTTTACTAAGAGTGCACAATTTCCATTTTCTAGATGATTACCTGCTGCGATGGCAGCACCTACTCCAGTTTCTGCACTAGTTCATTCTTCTACTTTAGTAACAGCAGGGATTTATGTAATGATACGATTCTGTAATATTTTCCCAGAGTGGTTATTTGTAATTATAGGTATTAGAGGTTTATGAACTTTATATTCGGCTAGATTAGCAGCCTGCAGTGAATTTGATGCTAAGAAGATAGTGGCGTATTCTACATTAAGTCAGTTGGGACTAATGGGAGTAGCTATATCACTAAATTTACCAATAATAGCTTTTTTTCATCTTGTAACTCATGCTATGTTTAAGGCTTTAATATTTATTTGTGTTGGCTATTTAATCAAAAATAGAGGGCATTTCCAAGACTTGCGCAGTTTAAATGGTGTTTGGGGAACGAGCCCATTATTGGGGGTAACATTAATTGTTAGAAGGTTGTCTTTAATGGGATTCCCCTTTTTAGCTGGATTCTTTTCAAAGGAGTTAATTCTTGAGAATGATATTTTGTTTATTAATAACATATTTCATAAATTGTTGTTATATTCATTACCTCTAACTTCTTATTATAGATGTCGGTTGGTATTTAAAGTATTAAATGGTAAAAATTATAAAAGAATATTTTGCGGGGAAGATAATAAAGTTTTATTATTTTCGTTATTACCTCTATATTTTGGTAGTATTATAATTGGAAGTTTATTATATCCTTTTTTTTATAGTTTAAGTTCAGTGTATCCTTGTTATTTGATGAAGTTTCTGGTAACTTTGTATATATTAATCGGGGTGATTTTAAGGTGGCGAGATATAAAGTTCAATACAAAATCTTTTATATGATTTAATAGTTCTATTTCATTTCTAGTTCCTTTTAATGGTGGTTATTGAACAAAAACTTGTAGTAATTTAGGCAGTGATTATTTCTATTTGCTAGATCAAGGAATTTTATCCAAGACTATTGGCAGAATAGATAATAATATAAATAATTCAGGGAATTGGCTAATTTCTTCTTTAGAATATTTTAGAAGTCCACAGGCAAAGTTTTATATTGGGAGTATTATAGCGTTATCAATACTTATTGGTAAAATATCTTAATAATAAAGACTTTCAAGAATTTTTTATTCGTTTTGCCTTGAAAGCTTAAAGAGGGAGTTAGTATTCCCGGAGGTCTAAAAAGTTAGTATATATATTATAATTGGTTGTCGGCCTTTAGAAGGGGATTTATTCCTCACTTTTT**GTGA**ATAAACAAGTTTTTAAATATAATAATTTTTCAAAGAGGAGTTATAGAACACCTTTTCATTTGGTAGAAGTAAGACCTTGACCTATACTTGCTTCATTAAGGGCTTTGGGAATTACCTTTGGTGGGGTATATTGGTGGCATTTTAATAATATGAACATATTTTTGCTAAGTCTATTATTTAAATTCTTAATAGCCTTTTGTTGATTTTCTGATGTTATAAAGGAAAATATGGGAGGTTTTCATAAAAGGGTAGTTATGTTTGGTTTTCGGTTTGGGATGATCCTATTTATTGTATCAGAAGTCCTTTTCTTCTTTTCTTTTTTTTGGTCATATTTTCATAACTGTTGGGGGCCTCAAGCAGAACTTGGTTTTATCTGACCTCCTTATGGATTTGATAATATAGTTATTGATCCATTTTCTATTCCTTTATTAAATACAGTGGTGTTACTTTCATCTGGGGCTAGTGTTACTTGAGCTCATCATGCCCTTGTTAATCAAGAATTTGATAATGCTACTTTAGGACTATTTGTGACTGTCTTCTTAGGTGGTTATTTTTTATTTTTACAAGGTAACGAGTATTTTTTAAGAGAATTTTCGCTTAATAGTACAATTTATGGAACAGTATTTTTTATGCTAACAGGTTTTCATGGGTTTCATGTAACCATTGGTACAATATTATTATTAGTTTGTTTCCTTCGTCATTTATCGGGACATTTTTCAACAGGGCAGCATGTAGGATTTGAAGCATCTGCTTGATATTGGCATTTTGTAGATGTCGTGTGGTTATTTTTATATTTTTTTGTATACTGATATGGTTATAATTTATAAAGCTTAGTGTGGCAGATGAATATGCGTTGGATTTAAGCTCCAAAGATGGGAGGTTTACTTCCCCTCTAAGGTATGAGACAGTGTTTAGTTGCACATAAGTTTTTGGTACTTAAAGAAAACGTAGTTTTCTCATAAATAAAACTTTAGAAATAGTTTAATTAAAATATTAGCTTTGGGAGTTAATGATCAAATTATTTATTTGTTTCTAAATTGTTTCAA**ATGTTAT**TCAGTTCAATGATATTAAGAGGAATCGTTGTTGGATTAATTGGTGCATGAATCTTTTGAACAACCCGGAAGAATTTTATAAGGTCTCGTGAAAAGTCAAGACCATTTGAATGTGGTTTTGATCCAAAGGATAAGGCTCGTATCCCCTTTTCTCTTCGGTTTTTCCTTATAGTTATATTATTTTTAATATTCGATGTGGAGCTTTCTTTACTATTACAGCTCCCCTTCCAACTAGATTTTGATAAATTTAAGGGACGTTTAGGTCTTATTGTATTTGTATGAATTTTGCTTCTTGGGACGTTAGAAGAATGACGTCGAGGAATTCTTAATTGAAAGGACTAATTAAAATAGGGGGTGTCCTCTTTAGGTATAGGAGGCTGCTAACTTTTTATAGAGTGACTAAGAATCGTTCACACCTCTAAA**ATGAA**TAATTTTAAATTGCGTCTTATACCAAAAGTTTGAAGAAGGTATTTATTTTTAGGAATATGTTTGGTTTTCGGTTTGGGAATTTCATTAGTAAGGAATAATATTTTTTTAGTCTGATTAGGGCTAGAATTAAATATGTTTGGGATAATTCCTCTTTTGAATTCTAGCCCTAATCAGACTAAAAAAATATTATTCCTTACTACAGGTGAAATAAAAGTATCCTTTTTTTATTTTTTTGTTCAAGTAATAGGTAGATTATTTTTTGCTTGAGGGAGTATATTAGGAGGATGGTTTATTATAAGAATGATAGGATTAATTATAAAGATTGGAGCAGCTCCATTTTTTTGGTGGGTTCCTCCGGTTATTACTCGATTAGATTGATTTTCTATTGGGATTATAAGAACTATACAAAAGGTACCAGGAATTTTTTTATTTCGGCTCCTATTCGATTTAAAATTAAGAATATGTTTAATTTTGGGTATTATCGGCTTTACTATTTCCGTTATAGGAATAAAATTTTCTTTTAAAAATTTAAAGCAGTTAATAGCCTGATCTTCAATAAGTAATATGAGTATATTATTTGTGTTAATTATGTTAAATAATAGGTTTGGTCTTATTTATTATCTGTTTTACAGAGTATTAGTACTATTATTCTGTTTTTTACTAAAATTGTTTTCAGAGAAAATAGTTTGTGGTTCTTTTTTAAATGGCAATTACAACTCTTATAAAATTTTAGTAGGAAGAACTTTATTAATATTTTCTGGGTTACCTCCCTTTGTTAGTTTTTTCTTAAAGATCTATTTTCTAAGGGGGTTTTATTTTTTTGATTGTGTGAAAATGTTATTAGATATAGAATTAAAAGGAACTAGAATTTCTATTTTTCATTTGGTGGGGAATGCACTAAATAGATGAAAAATAGTAATAATATTTATTATATTAATAGTATTCCAATCAGTTGGTTATGTTAAGGCTTTTATAAACTTATCTACAACAGGGTCTTCTCGTCTTTATTCAAGATCCAATGATGTTAATAAAAAGGTTAAGTTATTCTATTTATATATTTTTTTAATTTATTTATTATCAATATTCCTTGCATTTAATTAGTTTGCTTTAAAGTCAAAATGTGGAGACGTAAGTCTTGTAAACTTAAGATTGAAGTTAATCTTCTGAGGCAATTTATAATGTCTCTTGATAATTTATATTCAAAATATTAGCCTTGCATGCTAAAAAAGGGTTTAGTCCTCAAGTGATGTTATAATGTGAGGACTGCGAATGCAGGTTACTGTGATATAGTAAACTTTGGGTTTAGAAACCTCCACATTATAA**TTGGTTTATGA**TTTATTTACAAGGGCAGATTTTTGTTATTCTAAATTTTCATTTTTAAATATTATTGTTTGAGGTTTGGGATTAGTTTGGATTTTTTGGAAAATAAATCTATATTTTTTTTCTGGAAGACGTCTAAAAAGATTTTTATCATACTTGCTAACTTGGCACTATGATAAAGTTAAGACTTATTTTCTTTATAAGGTTAAGGGGTCTTATATAATAATTTCTGGAGCTTTCTTAATTATTTTAGGAAGTAATGTTTGGGGGTTGTTTCCTTATATCTTTGGGGTAACAACACAGATGGTACTAACTTTTTCAATGTCTTTAGTTATTTGACTATGTATTGTTGTATCAAGAATGGAATATTCGTTTATTGGATTCCTAACCCATCTCACCCCCCAAGGATCTCCCGGCTATTTAGCCCCTATTCTAAATTTAATTGAATTAGTTAGGAATATAATACGTCCTTTAACTTTAGCTTTACGGCTAAGAATAAAAATAACCACAGGTCATGTATTTATTAGATTGATGGGAACAAGGGGAAGCATTTGTTTATTTTCGTTCAAGTTCTTTTGATTGTTTTTTGTTATTTTAATGATGGGATATTTATTGTTTGAGGTTGGAATATGTTTTATACAAGGATTTGTTTTTAGACTGTTAAGAGTGCAATATTTAGGAGAGCACACTTAAAAGAATAGTAGTTTAATTAGAATACTGCATTGTGGTTGCAGAGGTGTATTTAAATACCTTTCTTCTTGTATAAATCTAAGTTATATATTAAATCTTTTCGCCGAGACCGTTCTGTATTAAAGGCG**GTGAAAGGAGG**GGTTTATGATTTACCCTCTCCAAAGAAAATTTCTTATTGGTGGGGTTTTGGTTCTTTACTTGGTTTATTTTTGGTTATTCAAATATTAACTGGATTATTTTTGGCTATGCATTATGTATCTGATTTAAAAGTAGCGTTTCAGTCTGTAGATTCTATTAGCCGAGAGATAGAATGAGGTTGGTTAATTCGTAGAATTCACGCTAATGGAGCATCCGCCTTTTTTCTATTTTTATATTTGCATATTGGTCGTGGGATATATTATGGTTCTTTTTTATATAGACATACTTGGAACACTGGTGTTGTTATATATATATTGGCAATGGCTACGGCTTTTTTAGGTTATGTACTACCATGAGGACAAATGTCTTATTGAGGGGCCACAGTAATTACCAAATTTTTTTCTACTGTCCCTTATATAGGGAATGATTTGGTTCAATGAATTTGAGGGGGATTTGCTGTTGGTTATCCCACATTAACCCGGTTTTTCTCTTTGCATTATTTATTGCCTTTTGTTATAGCTGCATTTGTATTAATTCATCTAATATTTTTACATGATACAGGATCTAATAATCCATTAGGGTTGAGATCTGCTGGAGATAAGGTACCTTTTCATCCTTATTTTACTATAAAGGATATATTTGGTTTTAGAATAGTTCTTTTTTTGTTCTTTTTTATTGCTGTTTATTCTCCCGATTATTTTGGAGACCCAGAAAATTTTATAGAAGCCAACCCATTAGTGACTCCTATACATATTCAACCAGAATGGTATTTTTTACCTGCTTATGCTATTTTACGGGCTATTCCTAATAAGTTAGGGGGTGTAGTAGCCTTATTGATGTCTATTTTAGTATTATTTATATTTCCTATAATTTCTTCTTTCTGTAATCGAGGATATTTCTATTCTGTTTTACATCAATCTCTTTTTTGAGTTTGAGTGGGGGATATATTAATATTATTATGAATAGGGGCTCGTCCTGTGGAAGAACCGTATGTTAGAATAGGAGCCGCTGGTACAGTGGTATATTTTTTGTTTTTTTTATTAACTCCAATTTTTAGGTGATATCATAGGGATATGTTTACAGAATATCTTAGTCTTCAAGAAGGAGAAACTGATGAAGATGATCCTGAACCACAAAGGGAATTTGGTTTGAACATGGGTAAATCATTTACCCTATTTAGAAGAGGCTAAAAACCTCCCAGATTAATTTGAAGTATTATATTTCCTTTAGCGATTTTAATATTCTTCAGGGAGATAAAATCTATTATAATTGGTTTAGTTATATTA**ATGTTATTATT**TAGCTCTTATTTAATCGGAACTCATTCCCATTTTATATATTTCCATTCTTATGGGATGTGCTTAGATAATATTTCAACTTTTTTAATAGTTTTAACATTATGAGTCACTCTGTTGATGAGATTATCCATGAAAAAAGCTAAGGAACTTCAAAAGTTGTTATTTTGTTTCATTTTATTAAATTTTATTTTAGTTTCAGCATTTGTAAAAAATAGTTTGTTAGGGTTTTATATATTTTTTGAATTGTCTTTAATTCCTACATTATTAATAATCTTAGGGTGGGGAGTGCAACCTGAGCGAATACGAGCAGGAAGTTATTTAATGATTTATACTTTAGTGGGTTCTTTACCGTTATTGGGTAGAATTTTATACATGGATTTTCATTGTGGTTCAGTTAAGCTTTTTATGCCTGAAATTTCTATATTCTTTTTCTCTAATTCTGATTATAGTTATTTTTGTCTTTTATGAATTTTAGCCTTTTTAATAAAGTTGCCAATTTATGGTGTTCATTTGTGGCTTCCTAAGGCACATGTTGAAGCTCCTGTAGCAGGTTCAATGGTTTTAGCAGGGATATTACTAAAGTTAGGGGCTTATGGTTTAATGCGTTCATTAGACTTTATTAGTTTAAATATCTGCTTTTGGGCAGATTTCTTTTTTACTTGAGGTATTTTTTCTATGTGTTTGGTAGGTTTTATGTGTTTTCGTCAATGTGATTTAAAGTCTTTAGTGGCTTATTCCTCGGTGGCTCATATGTCTTTAATCTTTGCTGCTTGTTTTTCAAGTGATGTTATAGGGATAAAAGGAATAATGGGGATGCTTATCTCACATGGATTATGCTCTTCTGGTCTTTTTTTTGGTGTACAATGTTTATATGAAAAAAGGGGATCTCGAAGATTATATTTAAATCGGGGGATATTAAGTTTGTCACCCTTATTTACCTTTTTTTGGTTTATATTGTGTGTAGGTAATGCATCAGCTCCACCAAGTTTAAATCTATTAAGAGAATTTTTTTTAATTTCTAGTATAATACTATACGGGGGTGCATTCGGTAGAATATTTTGTGGAGTATCTGTCTTCTTAGGTGGTTTATTCAGTATTTATTTATATGTATTGGTTTGTCATGGAAAGTGATCTCTCTTAAATAATTTTTGAATGCCCTTTACTTCACGCCATTATGCAGTATTGTTATTGCATGCGTTACCTTTATATAGTTTATTATTTTTAAGCAATTATATATTTTATTATATCTAAAGAATCTGTAGTTTATAAATTTAAAACCTATCGTTTACACCGATATAAAAGATTAT**GTGATCTCGG**ATTTAATTAAATTTTTTATGTTGGTGCCTTGGCTTTTTGTATTTGTAAATGTTTTACTTTCAATACCTTTTTTGACATTATTAGAACGTAAGGTTTTAAGGTATATACAATCACGAAAGGGCCCAAATAAGGTTAGTTATATGGGGCTTTTGCAACCTATTGGAGATGGGGCTAAGTTAGTATTAAAGGAATTAGGTACTCCTAATATGGCTAATATGATTTTATTCTGGGCTAGCCCAGTAATATCATTCTTTTTAATGATATTGGCTTGAGCTGTTTTCCCATCACCCTTTTTGTATTTTTCATTTAATTTAGGAGTTGTATTTTTTTTATGCATAGCCAGATTACAAGTTTATACTCTTTTGGGGTCTGGTTGGGGTTCTAAATCTAAGTATGCTTTATTAGGGTCTGTTCGTGGTGCAGCCCAAACAATATCTTATGAAGTTTCTTTAATATTTATAATTTTATTTCCATGTAGATTGGAATTTTCCTATAATTTTGCCACTTTTATTGATAAGTCATATAGGTATATATTAATTTTATCTCCTATATTTTTAATATGATTCATTTCATGTTTGGCCGAAACCAATCGTGCCCCTTTTGATTTCGCTGAAGGAGAAAGAGAGTTAGTATCTGGGTTTAATGTAGAATTCTCTGCATTTAGTTTTGCATGCTTGTTTTTATCAGAATATGGAAATATCTTATTAATGAGATTCCTCACAGCGATTTTCTTTTTCCCCTCCAGCTATTTATCTTTTATCTTATTCGGAAGATTTTTTTCCTTTTGTTTTGTGTGAGCACGAGGTTGTTTACCTCGCTTTCGGTATGATTTTCTTATGGCAATGGCTTGAAAGATTTTTTTACCCGTTTGTTTACTGGCTTTTTTTGTTATTTTAATCTAGAGAAAATTGAAATAGGATAATAGTTCCTTTTATAAAAATTATTAATTTTTATTTTCTTATT**ATGATGGGGTCTAGT**ATTTTCTATATTATATTTATATTAAAATGTTTGTTACTTTTATTTCGGTTAAAGAATATGCTTATAATATTATTTGCATTTGAATTTATGATAGTAAAAGTTTTATTTGCATTTGTATTTTTGAGAATCCCCTTTGACCCAATTACCCTATTAGTATTCCTGGCTGTAGTTGCAGGGGAAGCTAGATTGGGGCTTACATTATTAGTATCTGTACTTCGTCAAAGAGGTAAAGATAAAATTTTGAAATTATCCTTACCTTCAATTGAGGGGTTTTAAAATATTCTTTAACGTGAGAAATTCATTTGTGAAGATTGGTTTCGGCCCTTTTGTTGAGTGTTTTTCGCTCTCTCATGTATGCCACAAATGTCAAATATTCCTTTTTTGGTTGTAAGATTTGTAATATTTTCTTTTTTCGTTATTTGAATTTTAAATGTTTGATTTAATATTTATGGTGTAAGTAAATTAAAGAATGTAGACAAAAAGAAGAAGACCTTTTTCTTGTGACCTTTAAACTAGTGTTTTCTTCGATAGCTAAAATATTATAGCGTGGTCTTGAAGCGGCTAAGGTGATGTAATTTCTCGGGGAAAACGGAGTAGTCTAATTTTAGGGCATTGGGTTGTAGCCCCAAAAGTGAAGTAAATTTCCTCCGTGAAAAATTAAATAATAGGAACTGATAGTATAATAATTGCGTTGAATTGCAGATTCAAAGGTGTTTTAAAAAACTTAGTTCTTT**TATTAATTTGGTTTTGGTTAGATGAAGAATAAGCACTAAAACTATCATAAAAATATTAAAGAATCTTCTTCTTTTTC**ATTAGAGATTATTAATTTAGTGGTGGGTTTTTCTAAATAAAAATGAAAGTTTGTTTAAGTTAGTGTTGAGGAGGGGCTAACAATAGGTGCCAGCAGCTGCGGTTATACCTATTCTTATCTTCATGTTTGGGTAAAATACAAGTTGTTATTTTGTTAAAAATTAGTTTATTCCGGTTAAATGGTTTATAAACTTCTAATGTTCGAAAATCAAATAAGTAAATTGTCAAAGTTATTATAAAAACCGAGATTAGGTACCTCGTTATTAATAAATGTAAATAATAAAACCCAGAGGAGTAAGGAGCCTGAAACTCAAAAGACCTGGCGGTGGATGTTCTTCCTACCAGAGGAGGGTGCCAAGTAATAGATAATCCGCATAAAAATAAACCCTTTTAATTAAAATCAGTGTACGGCCGCTTACAGATAATTTTTATAGAAATAAGAAATTATCTTAATACTCTAATAATTAAAACTAGTTAAGACAGGTCAATGTGCTGTTGATAAAGGGGGTTAAAGGTGCGCTATTATAACATAGGATAAAAGCTTTCAAGATGTAAATTTTGAATCAATTAAGATTTGGTAGTAACGAAAAGTAATAAATATGTACGCGTGAATTAAGGTAAAGCTCTGTGTACACATCGCCCGTCGTTCTTTTTGAAAGAAAAGAGAAGTCGTAACATGGTAGCCTTAATAGAAATTGTGGCTTAAAAGGTAGTATAAATTAAAAATTATTATGTGAAGCTGTTAACTTTGTGATGTGCAAAAAGCACCCTTTTAGAAATTTATTAGAGAATAAGTTATATAAACTATAGGGCTCATGACCCTAAAAAGATTATTTTATATCTTCTCTA**ATGAAAACTGTA**GGAGGGTTTGGAAGAGGTTTTATATATGGCCTTAAATTACAAAATGGTACTAGTCCTGTTATGCGTGAAGCTATAATATTCCATGATGAAAATTTAGTAGTTTTAGTTGTTATTGCTTGTTTAATAGGAGGATGTATAATATTATTATGAGATACAGACTATAGTTCTTCAGATTTTGTTGATCATAAGTGATTAGAAGTTGGATGGACGTTACTTCCAGTTTTTATTCTTGTTGGATTAGCTTTACCATCTTTAGAACTATTATATTATATGGATAGTCCTTCATCAATTACCCCATTTGCTACATTAAAGGCAATTGGGCGGCAGTGATATTGGTCATATGAAATTAGAGTTTTAAGTCAAGAAAAAGTTGAAAAAATTTCATATGATTCTTATATGCTTCCGGAGGGGCAAGAGAAAGAAAAAGATGATGATTTAGCGGGATTTCGACTGCTTGAAGTTGATAATCCTATATTTTTACCCCGAGGGGAGTATATCCGCCTTTTAGTTACAGGAGGAGACGTAATCCATAGATTTTGTGTCCCAACCTTAGGTGTAAAGATTGATGCAGTCCCCGGTCGATTAAATCAAACTTATTTTTTCCCTTTGGTTTTAGGTAGATTTTATGGACAATGTTCCGAGCTATGTGGGGCTAATCATAGTTTTATGCCGATTAATGTAGAAGTTATTCCGGTGGATGTGTTTTTAAAGTTTTTTAAGTAAAAAGACTTTTAGGTTATATTTAGACCTTTAGCCTTCAAAGCTAAAAGAGAAAATTATTTTCAAAGTCTGTGTTTTCAGATTATTTTAAAAAGCCTTCAACAAATACTGTTAATATTAGCAGTGGTAATTGA
