## Supplementary for "Mitochondrial Genome-Based Phylogeny of Turbellarians and Evidence for Accelerated Mitochondrial Evolution in Symbiotic Species": annotationStylochoplana maculata KP965863.docx

ATAAAATGGCAATCGCGATGATTGCATTCTACTAATCATAAGGATATAGGTACTTTATACCTACTATTCGGAATATGATCAGGGCTTGTAGGTACAGCCTTTAGTTTTCTTATACGTTCAGAGTTATCCCAACCTGGCAGTATACTAAAAGATTCACAATTATATAATAGAATTATTACGGCACATGGCCTTATAATGATTTTCTTCTTTGTGATGCCTGTACTAATTGGTGGTTTTGGTAACTGGTTAATACCTTTATATTTAACTAGACCGGATATGGCCTTCCCCCGCCTTAAAAATATGAGGTTTTGGCTTTTGCCACCAGCCTTATTTTTATTACTAGGATCTTTCGTAGTTGAAGGTGGTGTGGGAACAGGTTGAACAATATATCCTCCTTTATCGTCAAATATAGCCCAAAGAGGACCAAGGGTGGATTTAGCTATATTTTCTTTACACCTCGCAGGTGTAAGGTCAATACTAGGCTCAATTAAATTCATTACTACAATAGTGAATGCCAAGATCCAAGTTTCATGAGGCCAATTACCTTTATTCCTATGGGCTGTTATGGTTACCGCCTATATGCTGGTTCTTTCTCTGCCCGTTTTGGCTGGTGGCCTAACAATGTTGTTAACAGATCGAAAATTTAATACAACATTTTTCGATCCGGGAGGTGGAGGTGATCCAATCTTGTTCCAACACATATTTTGATTCTTCGGACACCCTGAGGTATATATCTTAATCCTTCCTGGATTCGGAATGATTTCTCAAATAGTAACTTACTATAGAGGAAAGGATAGTGCCTTTGGTCATATGGGAATGGTATACGCAATTCTTGGAATAGGATTTCTAGGATTTATTGTATGAGCTCACCATATGTACACTGTAGGTCTTGACATAGATACTCGTGCTTATTTTACAGGAGCCACAATGATTATAGCGGTTCCGACTGGGATTAAGATATTTAGTTGATTAGCAACGTTTTATGGTCGTCCATTAAATCAAGAAACAGATAGAACTGGCCCCATTTGAGCTACAGGATTTATATTTTTATTTACGCTTGGAGGTTTAACAGGAGTAGTTCTGGCAAGAGCAAGCTTAGATATAAGATTGCACGATACATATTATGTGGTAGCTCATTTCCATTATGTATTGTCAATGGGAGCTGTATTTTCTATATTTGCTGGACTAGTTCATTGATGACCCTTATTCACAGGAACTGCTCTAAACTTTAAGATGGCCATGGGCCAGTTCTGAATAATGTTCACAGGAGTAAACTTAACCTTTTTCCCCCAACATTTCTTAGGGTTGGCAGGTATGCCCCGTCGTTATTCGGATTACCCAGATGGGTTCGCTTACTGAAATAGGGTTTCTTCATTAGGGTCACTAATTTCTGTAATAGGAGTAGTGTTCTTTGTGGTAATTATATGAGAAGCTATAATAAGCGGCCGTAAGGTTCTATACGTAATTTATCCTCCTTCACAAGTTGAATGGATTAGTGGGGAATGTTATCCCTCAAGCTTTCACACAAACGAATGTGTTAGAAAGGTTTTTAATTAAAGCTCGAAAGCTATTACTTTGAGCGTCAGATTCTTACTCTGAAGAAGAACATCTAGAGTTCTCGAGCTAAGACTTTATTTGGTATGGCAGATAAATGCGCTGGATTTAAGCTCCAGAGATGAGATATTTTTACTCTTCCTATAAGAAAGTGTAAACATTGAGCACATAAGCTTTTGAAGCTTAAGGTAAACGTAGTTTTCTTATAATTTTGGACTTCCAAGAACAATGAGTTCGTTTTGTTTTGAACACAAAAAGAGGGGACTGAATATACCCGGGGGTCTTTGCTAGATAAC**ATTAAAAA**ATATAAATTCAGAAGCGAAAACAAGGCTGGTCCTTTTCATCTTGTTGAAGTTAGACCTTGACCAATTTGCGCCTCTCTTAGTGCGCTGGGAATAACCTTCGGTAGAATATATTGGTGGCACTTCAAAAAACCCAATATAGTTATATTGGGTTTTGCACTCAAAATATTAGTCGCTTTTTGTTGATTCTGGGATGTTATAAAGGAAAACATAACAGGGTACCACAATAGGGTAGTAGCTTGGGGTTTCCGTTTTGGAATGATATTATTCATTCTATCAGAAGTAATGTTCTTCTTCTCATTTTTTTGATCCTATTTTCATAAATGCTGAGGACCCCAATTTGAGTTGGGCTACAATTGGCCTCCTTATGGGTTTAACACAGTAGTAATAGATCCTTTTTCCATACCGTTGTTAAAAACGGTTATACTTTTATCTTCCGGGGCGAGAGTAACATGGGCACATCATTCGTTAGTAAGACAAGATTATGATAAAGGATTAATAGGGTTATTTTTAACTGCTGGGTTGGGCGCCTACTTTTTATTTTTGCAAGGTAATGAATACTTTCTTAGCCCTTTCTCAATAAAAAGAACCGTTTACGGAACTGTATTCTTTATGTTAACAGGATTTCATGGATTTCATGTAACTATAGGAACAATACTGTTAATAGTGTGCTTTTTGCGCCATTTTTATGTTCATTTCTCTAGAGATCAACATGTAGGCTTTGAAGCATCAGCATGATACTGACATTTTGTAGATGTAGTTTGATTATTTCTATATTTTTTTATATATTGATATGGTTTCAATATGTAAAATGCTTTAAAGTCAAATAGCGACGTAAGTTTTGTAATCTTAAGATTGAATTTAAATTCTGAAGCAAACACTTATACGTGAGGAATTCAATAAATGAAGGTTGGTTTCGGCCCATCACGAGAGTATACTCTTACTCCCTCATGTTTGCCTCAAATGTCAAAATTACCTTTCGTTTTTATTAGGTTTGTTTTTTTTGTCTTTTTGTGTTTATGATTATTAAAAGGGTTATTTAATAAATCAAGTGTTTATTCGGAAACACAAAAAGACAAAAAAAACAAACCTAATAAGATATGACCTTTTTAAGTTAGAAGGTAGTATAAAAATTATTATATAAAGCTGTTAACTTTATGATGTGATCTAAAACCACCCTACTAGATTATTATATTTAAATTTGTAGTTTAAATAAAATATTCCGTTTACACCGGCAAGATAGATAGATTGTCTCAAATTTAATTGTTAGATTTTTCTAGA**ATAATACCTTGA**CTTTTGGTTTTTGTTAATGTATTGATAGCTATTCCCTTTCTAACTCTTTTAGAGCGAAAGGTGTTAGGGTATATACAATCTCGTAAGGGGCCAAACAAGGTTAGATATTTAGGACTTCTACAACCAGTTTCCGATGGGGTAAAGTTAATTTTAAAGGAAATTGGAACGCCCAATATGGCCAAAATGCTATTATTCTGGTTAAGACCATTACTATCTTTCATATTGATGATCTTAGCATGATGTGTGTTCCCCTCACCGACCTCATATTATTCTTTCGGCACAGGAGTAGTGTTCTTTCTATGCATCGCTAGATTGCAAGTTTATACCTTGCTGGGTTCGGGGTGAGGATCGAACTCTAAGTACGCTTTATTAGGATCCGTTCGAGGGGCCGCCCAAACAATATCATATGAAGTGTCACTTATATTTATACTATTTTTCCCTTGTAGTATGGAAATAAGATATAATTTATTTTATTTCGTTGATAAGACATATTGGTATGTTTTGTTTATGTTTCCATTATTCCTAATGTGAGGAATTTCTTGTTTGGCTGAAACAAATCGAGCACCTTTTGACTTCGCAGAAGGGGAAAGTGAGTTAGTATCAGGTTTTAATATAGAATTCGCAGCATTGGGTTTCACGTTCTTATTTCTATCCGAATATGGTAATATATTACTTATGAGATATTTAACGGCACTTCTATTCTTTCCTTGTGAGTACTTATCTTATGTTTTATTAGGTAGAGTTTTCGCCTTCGCATTTATTTGGGTTCGGGGAACCCTACCACGATTTCGATATGACTTCTTAATGAATGTGGCTTGAAAGTATTTTCTACCTATCTCTTTATTTTGTCTTTTCATCTTAATAAAATAAATAAGAAGAGGTGTTTATTAAAATAAAAAAAGCTGCTAACTTTTTTTAGAGTGGGTAAGATCATTCACATCTCTTAG**ATGAGGCCTC**GCTTTAATAGTAAATTAACATTGTCACTTAATCAATGATCAAGGTCAATAGGATTAAGAATTATATTATTAAGTGGGATATTTATTTCCCTTATTAGTAAAAATATGTTCATATGTTGGATTGGTCTAGAAATTAAAATGTTTGGTGTAATTCCTTTTTTAACGTCAAAAAAAAGAGAAAATAGAACCTTCTTTGATAGAAATAAAGAGATAAATGTTAGATTTTTCTACTTCTTCGTGCAAGTTATAGGAAGTTTATTTTTTGGGTGAGGGGCAATACTAGGTGGTTGATATGCTATAAGAATTATAGGATTAATGATAAAGATGGGTATAGCTCCCTTTTTTTGATGAGTTCCATCAGTGATACCCCGCCTAAAATGATTATCGATAGGTTTATTGAGAACACTCCAAAAGGCCCCCGGTCTATTTTTATTTCGTTTAATGTTTGATATAAGGTTAGATATTTGCATCTTATTTAGCTTGGTGGGATTTTTTATAGCAAGGATTGGTATAAAATTTTCATATAAAAACATTAAGAAATTGGTTTCATGATCATCCATTAGAAATATGAGTATTCTTTTTCTATTAATAGTTCTGAAAGGTAAGTTAGGATCAATCTACTACCTATTTTATAGTATTCTAGTCCTTTCTTTATGTTTTTCCCTCCAAAGGAGGGAAGTAAACCTAATATCAAGTAGATTTGTTGGAGGGAAAAACATAAAGAAAATAGTAAGGCTAAATAGGTTATTACTAGTTTTCACAGGATTACCCCCATTTATAAGTTTCCTTTTGAAGGTGTACTTCCTAAGGGGATACTTTTTAAATGATAGGTGTCAAATGCTGATAGAGATTGATTTCAAAGGTAATGAAGTGGGGTTCTTTTATTTGCTAGGTAGATATCTAAAAGGTTGAAATATAGTGTTAGTTTATATAATACTAATAGTGATACAATCCATAGGTTATGTTAAGGCTTTCATAAATATTAATACAAGGAGAAGTTCTTCTCCCTTGAAATCAATACGCTTAAAAAATAAGGATAAGCTAATCTTCACTATTTTTCTTATATTATATTTTAGGAGAATAATGGTTATATGACTATAATATAAGTTAGTCAAAGAATGGCATTGGGTTGTAGCCCCAAAGATGATAATAAAACTTCACTTATAAATAATATACGGCCAAATAATAGAGTATTTAGCCTAAAGAAGTTTATATAAGGGTTAAAAACCCTTAAGATTTCAAAAGTCGCTTCTTATTAGAAAAAGACAGTTGTAAACTGCAGTATATGTACAAGATGGTTTAGGGGTTTTCCGCTATCCATATATATATACGGCCAAATAATAGAGTATTTAGCCTAAAGAAGTTTATATAAGGGTTAAAAACCCTTAAGATTTCAAAAGTCGCTTCTTATTAGAAAAAGACAGTTGTAAACTGCAGTATATGTACAAGATGGTTTAGGGGTTTTCCGCTATCCATATATATATACGGCCAAATAATAGAGTATTTAGCCTAAAGAAGTTTATATAAGGGTTAAAAACCCTTAAGATTTCAAAAGTCGCTTCTTATTAGT**TTGAAAAAAACAGATTT**GTTCCTTTTTGTCTTTTTCCTAAAAATATTGTTATTACTATTTCGCTTAAAGAGAGTTTTAATAATATTGTTTGCCTTCGAGTTTTTGATTGTGAAATTACTATTTTCTTTTGTCTACTTAGGCACACCATTCGATCCAATTTCAATATTGGTATTTTTGGCCGTAGCGGCGGGGGAGGCAAGTATAGGGCTAAGACTCCTTATATCTTTACTTCGCCAAAAGGGTAAAGATAAATTAAAAAGACTACAATTCACCAAAGAATATGAAGGATTCTAAATTATTAAATTTAGATAAAGAGTTTATAGCATATATAGTGTGTTGAATTGCAGATTCAAAGGTGTATTAATTTACTAAACTCTAATTTATTTGGTCTTGGTTAAATAAAGAGTTATTACTAAAACTACCATAAAAATATTATATCTTATAAAATTTATTAAATTATTTTAAAGGTTAGTGGTGAATTTTTATAAATAAGGGCGAAAACCCGTTTAAGTTAGTAAACCAGAGGAGTAAATTTAGGTGCCAGCATCTGCGGTTATACCTATCCTCCATCTTTATTTTTGGGTTAAATGCAATGTAGATTGTTCAACAAAAGCGACTTTGTAAGGTTAAATGATTAAAAGTTACTCAATCTTAATTTGTAACATAATTTCATTGCAAAAATTTTTATAAAAACTAAGATTAGGTACCTTATTATTATAAAAAGTAAATCCTTTGAACCCAGAGGAGTAATAAAACGAAACTTAAAAGACCTGGCGGTGGAAATCTTCCTATCAGAGGGGTGTGCCAATTAATAGATAATCCGCTAACTAGCTTACTCTAAGTTTTTAGTCAGTGTACGGCCGCTTTCAGGAAATTTTGTTAGAATTTAAAAATTTCCTATACCTAAATCTTAAATATGTAAGTATTTAATAAAGCTTAGTCAAGACAGGTCAATGTGCTGCTGATTTTAGAGAAAATGGTGCGCCATCATAAATGTGAAAAGGGATTCTAAAAAGAAATATTAGGATGAATTAAGATTTGATAGTAATGATATTAAAGTATGGTATCATGAAGTAGGTAAAGATCTGCGTACACATCGCCCGTCGCTCTTCTTAAAAAAGAAGAGAAGTCGTAACATGGTAGCTCTAATAGAAATTGGGGCTAAAGATTAACAAATATAATTAAAGTTTAACTTTAATGTGAACACAATTTCATAATCTTCCTAATGTTAGTGTAAATGGTAACGTTAAGCTTTCTACTTAAAAGAGTTTTTTGAACACATTAGAACTTTGTCA**ATGAAATTTTGGTTATT**ATTACTTATAGCAAAATATTTATGCTTGTTCTCTTTAAAATCAATACTTTTAGGCGTGTTCTTATTTTTATGTAGATTTTCAATAAGAATATTAATAAGGATATATTGTTTATCATGATATTCTTTAATATTTTTCCTTGTTTATATAGGGGGTCTACTGGTGTTATTCATATATATATCTTCTTTAAATTATAAACCAGCGTTTTACTCTTTAGAAGTAAGTCCGATAAGCAAAATGTTATTGAAGGTGAAAGGGATACTAATACTAACACTCAGCATCCTTCAGATAAGGTGGAAATTTAAAAGAGGGTTCTTTAAAAACCAAGATACTAACAATTTTAGTCTAAACCTTTTCAATGAAACGGAATTAATATTCTTAGTCAATGTAGGCCTATTATTATTGCTAGTACTTTGGGTTATAACAAAGCTATCTTTTTTAAGGCGGGGTGCTTTACGCCCAGTGTATTAATTAAAGGAAGAAGAGGTTAGTATATTTATTATTATTGGTTGTCGGCTAATAGAAGGAAGGTTAACCCTCCCACCTTTTGTTCTGTCATGAAATTTGAAATTTAAGGAAATAATTTCTATAGGTAAAAAGTATGGATTCCCTTATATATGTTCAAATCTTTTCCTT**ATTCTTAGA**ATTCTCCTCGCTTCAATTTTTAGTATTACTAAAAAAAGATTCATGATTGTTTTTAAAGTACTCCAATTAAAAAGTCTAGAACTAGATATATGTTTAATGGTGGATAAAATATCCTTATTATTTTCTTGTGTACTTTGTATAATAGTATCATGTGTACTAAAGTTTTCTTGTGTATATATGGAAAATGATTCTAAAAATATTCGCTTTACATGATTGGTACTAAGGTTTGTCTTATCAATGTTATGTTTGATATTTATACCTCATTTCTTCTTTCTCTTGATAGGTTGGGATGGTTTAGGGATTACGAGTTTTATTTTAGTTATTTACTATCTAAGAGATTCTTCGTGGGCGGCTGGAATGAAGACCTATCTAATCAATCGAATAGGTGATAGTTTCTTTATAATAGGATTAATTCTCTTTATATCAAAAGGATGCTGAGAAATAAAAAGAATAATAAATAAAGATACATTAGTTTTAATGGTTATCATAGGATGCTTTACTAAGAGTGCCCAATTCCCGTTCTCTAGGTGATTACCAGCAGCGATGGCTGCTCCAACACCTGTATCATCTTTGGTGCACTCATCAACTCTGGTAACAGCTGGTATTTATTTATTGATACGGTTTAGTCATATGTTTCCAACCTGAGGATATACAATCATTGGTATATGTGGAATGTGAACGCTATACTCTGCAAGACTAGCAGCTTGCTGCGAATATGATGGAAAGAAGGTTGTGGCTTACTCCACTTTAAGACAATTAGGGTTGATGGCCGTAGCCATTTCATTAAGTTTACCAATGATTGCATTTTACCATTTAATAACTCATGCAATGTTTAAGGCTCTAATCTTTATTTGTGTGGGATATCTTATTAAAAAAAGTGGACATTTCCAAGATTTACGAAGATTAAAAGGATTATGAATAAAAAACCCTGTTCTTGCAACAACATTATTAGTTAGGAAAGCTTCTTTAGCAGGATTTCCATTCTTGGCGGGGTACTTTTCTAAGGAATTAATAATCGAAAATAATATACTTATAATAAATTGAGTATTTCATTCTTTACTATTATTGTCACTCCCCCTAACTTCTTATTATAGTTGTCGACTTTGTTTTAATATTTTAAACGGGGCGAAATATAAATCAATAACAACGCCTAATAATGAAAGTAATGTATTGTTCTTTTCTATTTTACCATTGTATTTAGGATCTATTTTCGTAGGAAGCCTACTATACCCTAAATTTTACAGTGTAAGATCAATATATCCTTGTCACCTTATGAAGGTGTGTGTATTTATATTTATAGCTCTCGGGGCCTTACTGAGTTGATACGATGTTAAGATTAACAAAAAATCATTGATAGTTTTCTTTAGAAGAATATGTTATCTTACACCTTTCAAAGGATCTTTCTGAAAAAATAATTTCTCTTTAAAGGGTAAAGAATTATTTTGTATATTAGATCAAGGTATATTAAGTTCTACTATCAGAGTTCTAGATGGTTGTGTTAACAATGCTGGAGAAAAATTACATCAAAGGGTAAGATATTTCTTATCTCCACGAAGAGGCTACTATATTTTTAGTACAGCTGGTTTAGCTTTACTAGTAGGAACAACGTCTTAAAGGGGCTTTTAGTTAAAGATAATTTAAATCAAAATACCGGCTTTGGGAGTGGGAAATCAGAGTATTCTGTCTTTAAATA**ATGGGAAATATAATT**TTTAGATCCATACTAATTATTTCCGGACTTAGGGCCTTAGTAATATTATGGACCTTTTGGACATCCCGAAAGAAATTTTTAAAATCTCGAGAAAAGTCTAGACCTTTCGAGTGTGGATTTGATCCAAAGGATAAGGCCCGAGTTCCCTTTTCTCTTCGGTTTTTCTTAATTATTATAATATTTCTTATTTTTGATGTAGAAATTTCTGTCTTGTTGGGACTTCCTTACCAACTAGACTATGACAAATTTAAGGGACGTTTAGGGGTCCTTCTATTTCTATGGGTTCTGCTCCTAGGGACACTAGAGGAATGACGGCGGGGAATTTTAGATTGAAAGAAGTAGTTAAATCTAGTGTGAGGACTGCGAATGCAGGTTACTGTGATATAGTAAACTTTAAGTATGGATACTTCCACACTATGGTATATGATTTATTTTCTAGTGCAGATTTCTATTATTCAAAATTTTCTCTATTAAATAGAGCTGTGTGAATATTAGGGTTGGTTTGGTTATTTTGAAAA**ATAAACTTATTTTTTTCTACTCCTAATCGTATGAAAAAAATATTATCACTTCTTTTAAAATGGCATTATGAAAAA**ATAAAGATGGAAATGATAGATAAGTGTAAGAGATCATACTTAGTTAATTCTGGTCTCTTCTTTTTAGTACTAGGGATGAACTTATGAGGTTTATTTCCATATGTGTTTGGTATAACAACACAAATAGTATTAACTTTTAGAATATCCATAATAATGTGATTAAGGGTTGTATCATCATCAATAGAGTTCTCATTAAGAGGATTCCTATCTCACATGACTCCAGCGGGTAGACCCGGATATTTAGCTCCCATACTAAATCTTATAGAATTGGTCAGAAATCTTATTCGTCCTCTTACTTTAGCCTTACGTCTTAGAATAAATATAACTACAGGACACGTATTAATAAGTCTTATTAGTAGTAGAGCCGCTATAGTCTTCTTTAATTCCTTTCCAATGTGAACCCTATTAAGATTTTTAATAGGAGGTTATTTACTCTTCGAAGTAGGAATATGTTTTATACAAGGGTTTGTATTTAGATTACTAAAAGCACAATACTTAAGAGAACATACACATTAATACAAAAAAGAATTAGTAGTTTAAAAAGAACGTTGTGTTGTGGCCGCAAAAGTATACATTTATGTATCATTCTTCATTGAAAAGAAAAAGTGGTTCTTAT**TTAAGATCTTTTCGTCGAGATCATTCAGCATTAAAGATTGTTAAAGGAGGT**GTATATGATTTACCCTCTCCTAAGAATATTTCTTATTGATGAGGTTTCGGTTCATTATTAGGCTTATTTTTAGTAATACAGATTTTAACCGGACTATTTTTGGCTATGCATTATGTCTCAGATCTATATGTGTCTTTTAGTTCAGTAGATTCAATAGGCCGAGAGGTGGAATATGGATGATTAATTCGAAGAATCCATGCAAAAGGCGCTTCAGCATTCTTTTTATTTTTATATCTTCATATAGGCCGAGGGATTTATTATGGTTCATATATATTTATCCATACATGAAAAACTGGAGTAGTAATATATGTTCTATCAATGGCTACTGCATTTTTTGGATACGTACTACCTTGAGGGCAAATGTCTTATTGAGGGGCAACGGTAATAACTAAATTTTTCTCAACAGTGCCTTATGTAGGGGGAGATTTAGTGCAGTGAATATGAGGAGGGTTTGCTGTTGGATACCCAACACTAACCCGATTCTTTTCCCTGCATTTTTTATTACCATTTGTTATAGCAGCTTTTGTTGGAATTCATTTGATATTTTTACATGAAACTGGATCTAAAAATCCATTGGGGCTAAAATCCGAAGGAGACAAGCTGCCTTTTCATCCTTATTTTACAGTCAAGGACATTTTAGGTTTCAGAATTGTACTATTCTTTTTCTTCATAGTTGCAATTCTGCTTCCCGATTATTTTGGAGACCCTGAAAATTATATAGAAGCAAAAGCACTTGTAACCCCCATTCATATCCAACCGGAATGATACTTTTTACCTGCATATGCTATACTTCGTGCTATCCCTAAAAAGTTAGGAGGAGTAATAGCCCTCTTAGGTTCTATATTGGCTTTATTCTTATTTCCTTTTATAGCTTCCAGTTGTAATCGAGGATATTTTTATTCTCTAACACATCAAGCAGCCTACTGAGGATGAGTAGCTACAATATTAGTACTGTTATGAGTTGGGGCACGACCCGTAGAAGATCCATATATTTTCTTAGGTCAAGTTGCAACTATAAGATATTTCATATTCTTCGTAATAACATGAGGGATAGGAAAGGGCAGAAAGCCTTCTGTAAGTTAAACAATTTATAGCATACTTTTTTTAAGAAGAGGGCTAACCTTATTTCGTAAGGTCAACAGGATCATAACTTCCTTAGTTATTATT**ATGTTGATAACTAT**ACCTTACCTTAAAAGATTCCATTCGCACAAAATTTTTTATCATTCCTTTTGAATAAGGTTAGATAAATTATCTAGATTCTTAATAATATTAACTGTGTGAATTAGCCTATTAATGTGTTTAGCTATGAAAAGGGCCAAAAAAATAGGAAAGTTGTTACTTAGCTTTACTATTTTAAAATTAATCTTAATTCTGAGATTTATGAGTAAAAGTTTATTAGGGTTCTATATATTCTTCGAATTGTCATTAATTCCAACCTTAATAATTATACTAGGGTGGGGAGTCCAACCTGAACGAATCCGGGCAGGAAGCTACCTAATGATATATACTTTAGTTGGGTCTTTACCATTACTGGCAGGAATATTATACATGGATTTTCATTGTGGAACTATAAAGACATTTATTCCTATAGGAGAAATATTCTTCTTTAAAAAAAGTGACTACAGAATATTTTGCATGTTCTGGGTATTAGCCTTTTTAATAAAGTTGCCAATATATGGTGTACATTTATGGTTGCCAAAGGCCCATGTAGAAGCTCCTGTGGCTGGATCAATGGTATTGGCAGGAATACTGTTAAAGTTAGGGGCCTATGGTTTATTACGCACACTACACTTCCTGGATATGTCCCTATGTATATGAGGAGACTCTCTATTTATATGAAGAGGAGTTACCATGTGTATAGTAGGAATAATATGTTTCCGTCAATGTGATTTAAAGTCTTTAGTGGCCTATTCTTCTGTTGCTCATATGTCCTTAGTCCTGGCCGCTTGTTTCTCAAGGGACATTATAGGTATTAAAGGGGCTGTAGGGATGCTAGTATCCCACGGTCTATGTTCTTCTGGCCTATTTTTTGGGGTGCAGTGTCTATATGAAAATAGTGGAAGACGTAACATATTACTCAATCGCGGGTTGATTTGTTTATCACCTACGTTTACATTATTTTGATTCCTGCTATGTGTAGGTAAAGCTTCGGCCCCACCAAGCTTAAACCTACTAAGAGAGTTTATGTTAATGTCAAGAATGGTATTTTATGGAGGACCTTTTGCCGGAATTATGTGTGGGATAACTATATTCTTAGGAGGTCTGTTCAGAATCTATTTATATGTATTAATATGTCACGGTAAGTGATCTCCAAGCAATAAATATTGAAATCCTCCTACCTTTCGGAGACTATTGGTGCTCTTTTTACACATTATACCTTTATATGGTCTTATATTTATGAGAAAAAGAATTTTCTACAGCATATAAAAGGCCTTTAGGTTATTTAGACCTTGAGCCTTCAAAGCTCAGAGAGAGATTAGATATTTCATGGTCTGTAATTAGGCTCAAAAAAAGCTATAAGGGTACCCCCTTATAGGAAAAAAAACATTCAGAGGTTGATAGAAAAAAACTTCTATTTTTTTTCTATCAACCTCTGAATGTTTTTTTGATAGCTAAAATTTTATAGCGTGGTCTTGAAGCGGCCAAGGTGAAAGGAATTTCTCAGAAAATAAATTTATATTAAATTGTTAGAGGGTAAGTTATGTAAACTATAGGGCTCATGACCCTAAAAAGACATTTGTCTCCTCTAAAAAATCTTTTTTAGTTTATAAGAATATTTGCCTTCCAAGCAAAAGGTCCTCCATAAGAGGAAAAAGAAAAATATATATAGTTAATTGTTGTAAATACTATAAATTTCTGTAAAGAAAAACAAAACGTTGTATATTAGGATAAAACTCTCCCTTTTGAATCATGGTTCAATAAAGTCGTTTTTAAATAAATAAA**CCCGAATTCTTG**GGGTCATTTTTTATCCAATTAAGAATTATATATGTAAAAAAGCATTTTTATATAGAGATAAAATAAGAATAGATAATTTAGCCGACCAAGACAATATCTGGTTACAATCTATAATTTCATAGCATAATTAAAAAGGTTTTAACTTTTTTAATAAATAAAAGATTCAATATTTAAAATAATAAATGTTTAGGATGAAAAGTGTCCAAAACTAGAAAGTTTTATAACAGTAATTAAATTCTATGTTGTCTTTTAAAAGAGGTTGTTAAAAACTTTAACTAATTCTGATGAAATTAGTAAATAAAATAGATATAAGGAACTCAAAATATTTACACCTGTTTATCAAAAACATCTCCTTAAGTATATATTTAAGGTATTGCCTGCCCAATGTGAAAATTAATGGCTGCGGTATATTAACTGCACAAAGGTAGCATAATTAATTGTCTTTTAATTGGTGACTTGTTTGAATGGTTTGATGAGAAATATTTTTTCTTTTATCTAATTATAACTTATATTTTAAGGTACGAATACCTAAATAATAAAGAAAGACGAGAAGACCCTGTTGAGTTTTACAAAATATATTTGTTTAAATGGGAAATTTAGCTTTTTCTTAAAGTTCTTTTTTGATCCTGGTTTTCTTAAACTGGAAAAAGGATAAAATTACCACAGGGATAACAGGACAATAATTGTTGAGAGTTCTAATCGAAGCAATTGGTTGTTACCTCGATGTTGAATCGAAATTTCTCCATGATTGCAGAAGTCATAAGAGTGGGTCTGTTCGACCTTTGAAATTTCACGTGATTTGAGTTAAGACCGGCGTGAGCCAGGTTGGTTTCTATCTTCTATTGATCAATTTTTGTACGAAAGGAATGGATTGAAAAGTTAATCTTACTTTTATGAGGTGGCAGATTTAGATGTGATAAGCTTAGGACTTATTTAGGTGCTCATACCACCCTCATAACCCCTATTAGTTTAAACAAAACACTAGATTTGCAATTTAGAGATGGGAGATCCCCTTAGGAGATTGGGTAGAGGTAAA**ATTTA**TGGTTTAAATTTACAAAATGCAATAAGTCCAGTAAGACGAGAAACTGTATTTTTTCATGATGAAAAAATGTTTGTTATTATCATGATTGCAGCTCTAGTAGGAGGTAGTATATTAGGACTATGATTAAAATTTTATAGTTCCTATGACTTTGTTGATCACAAGTGATTAGAAGTTGGTTGAACAATGTTTCCAGTGTTCATACTCTTAGGGTTAGCCCTTCCTTCTTTAGAACTACTATATTATATGGATAGTCATTCTTCAGTACCCCCATTTGCAACCTTAAAGGTAATTGGGCGGCAATGGTATTGATCATATGAAATTAGTAGAAAGAATAGAGAATTTAGGTTCGATTCTTATATGCTACCATCCAAAGAACTAAATGGGGGCTTTCGACTTTTAGAAGTTGATAACCCTATTTTTGTTCCGCGGGGGGAGTATATCCGTTTACTTGTAACAGGAGGAGATGTTATACATAGATTCTGTATTCCGAGCCTAGGCATTAAGGTAGATGCAGTCCCGGGGCGATTAAATCAAACATATTTCTTCCCTTTAAACTTAGGTATATTCTATGGACAATGTTCAGAAATCTGTGGGGCAAATCACAGCTTCATGCCTATAAACATGGAGGTAATATCTAAGAAGGTATTTGACAAGTTTTAAATAAGGTTTTTAAGCGGAAAGCTAAAATTTTTAAAAAAATTTTATGATTGATTTCAGAACCTATGAGTAAGTTT
