## Supplementary figures and images for "Mitochondrial Genome-Based Phylogeny of Turbellarians and Evidence for Accelerated Mitochondrial Evolution in Symbiotic Species"

### circos.png

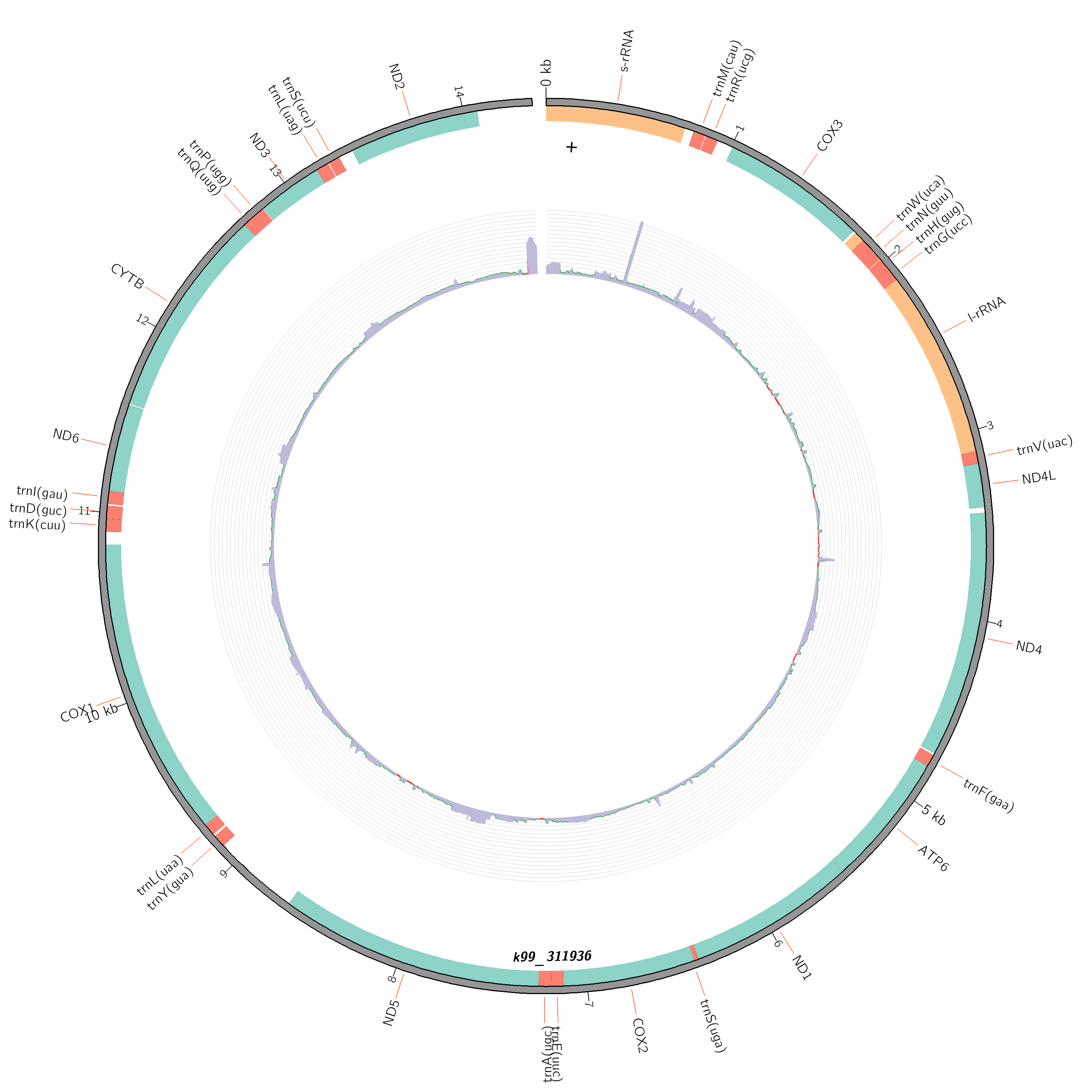

### circos.png

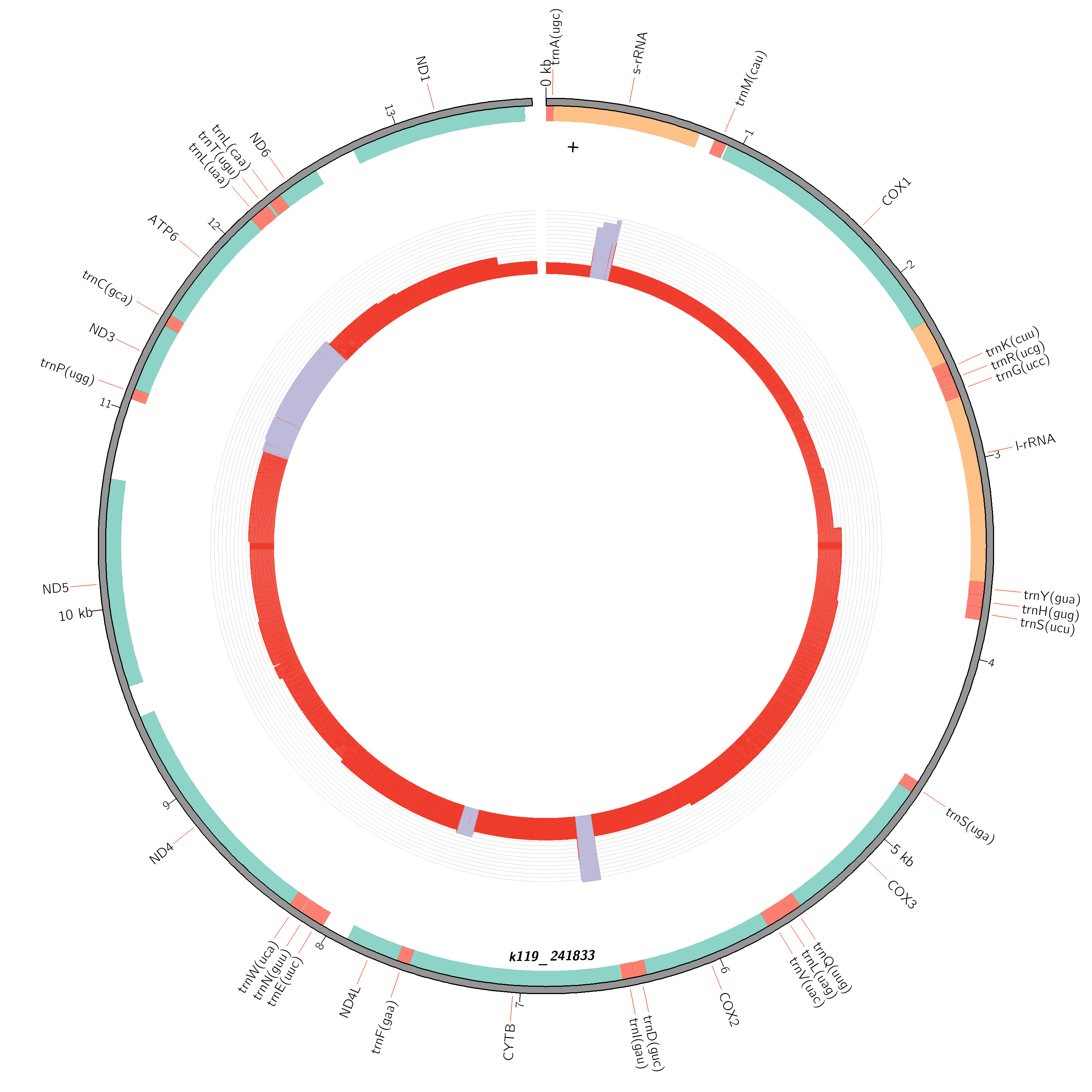

### Fig. S1 saturation-AA.pdf

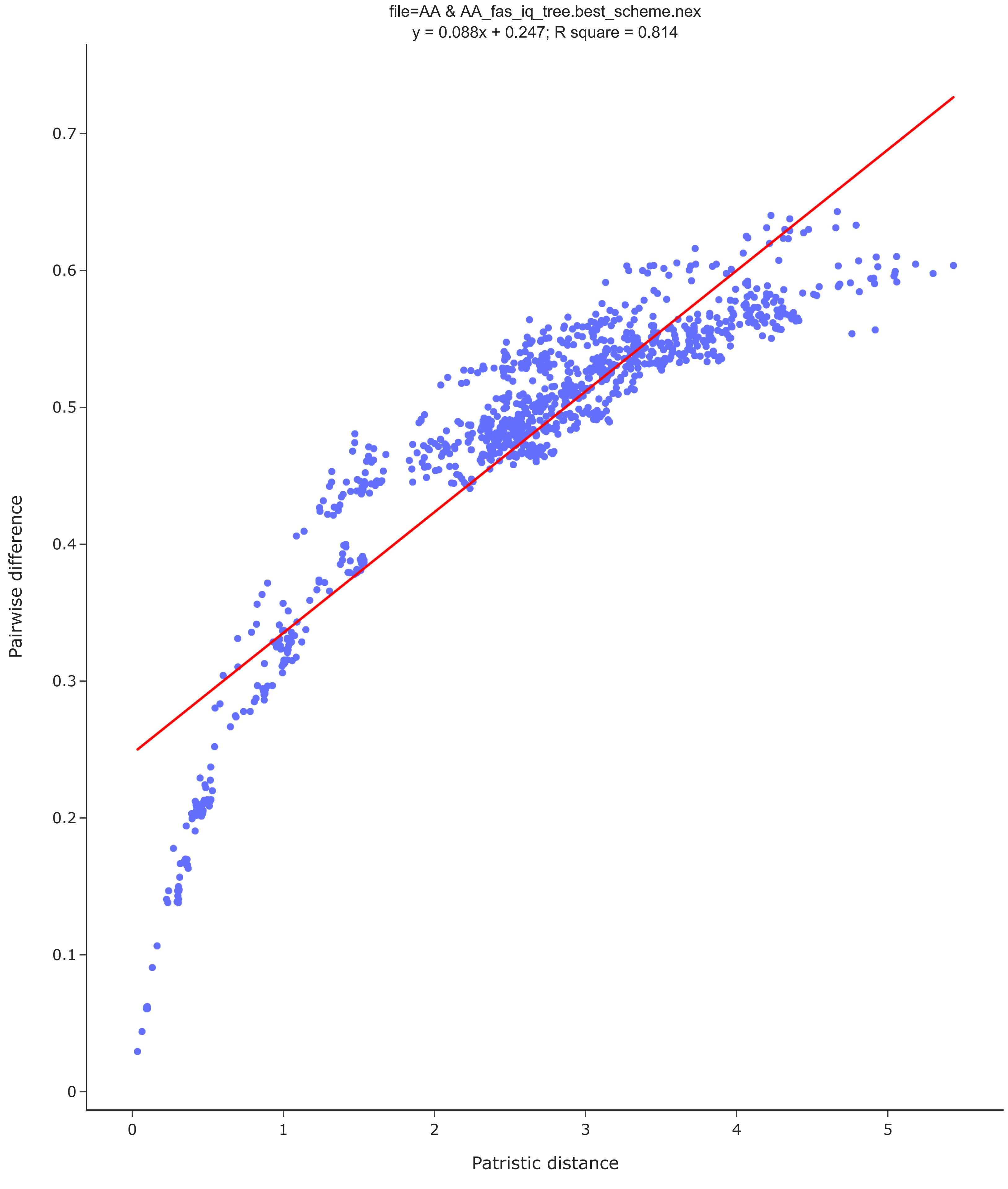

### Fig. S2 saturation-PCGsRNA.pdf

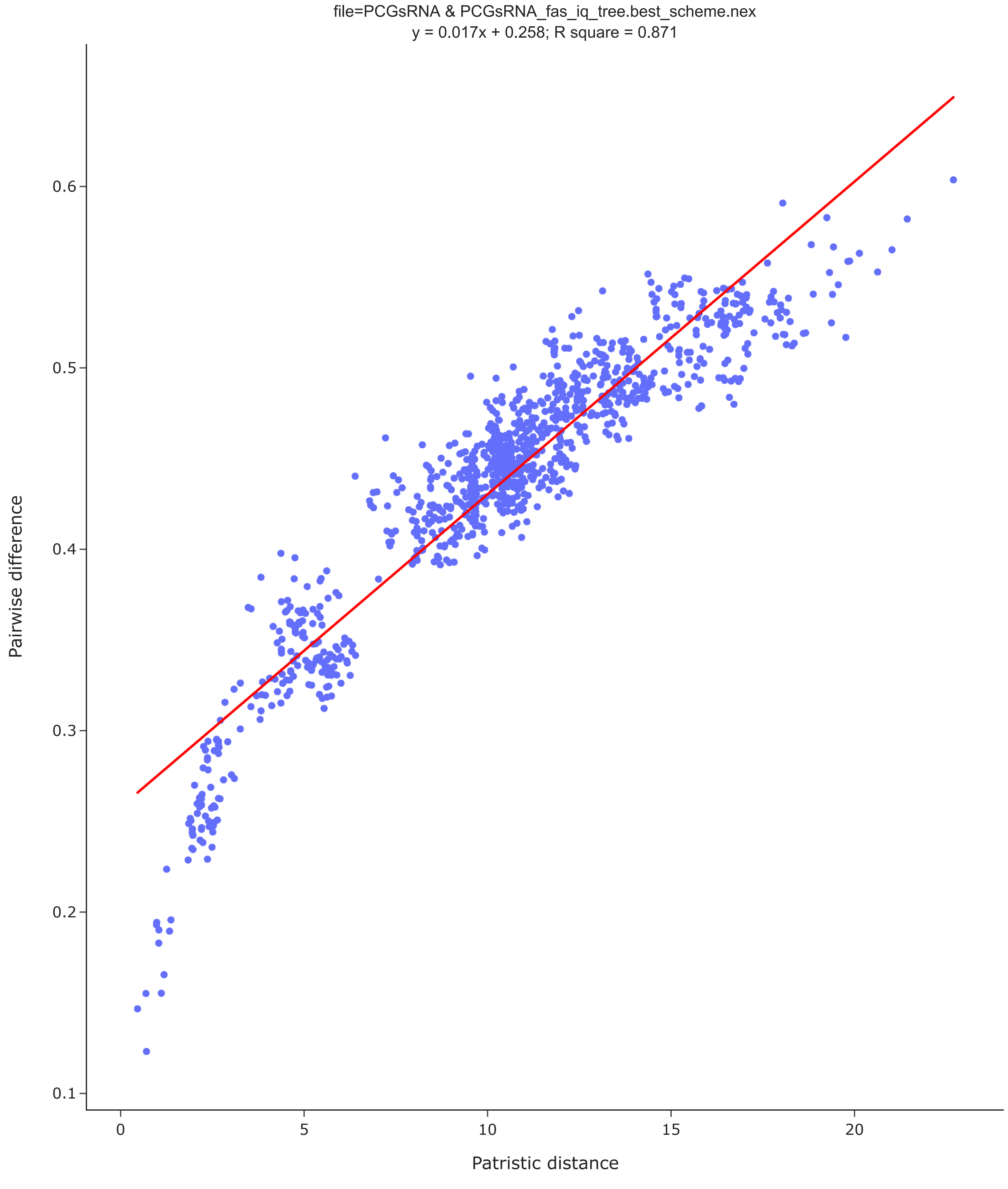

### Fig. S3 tRNA secondry structure.pdf

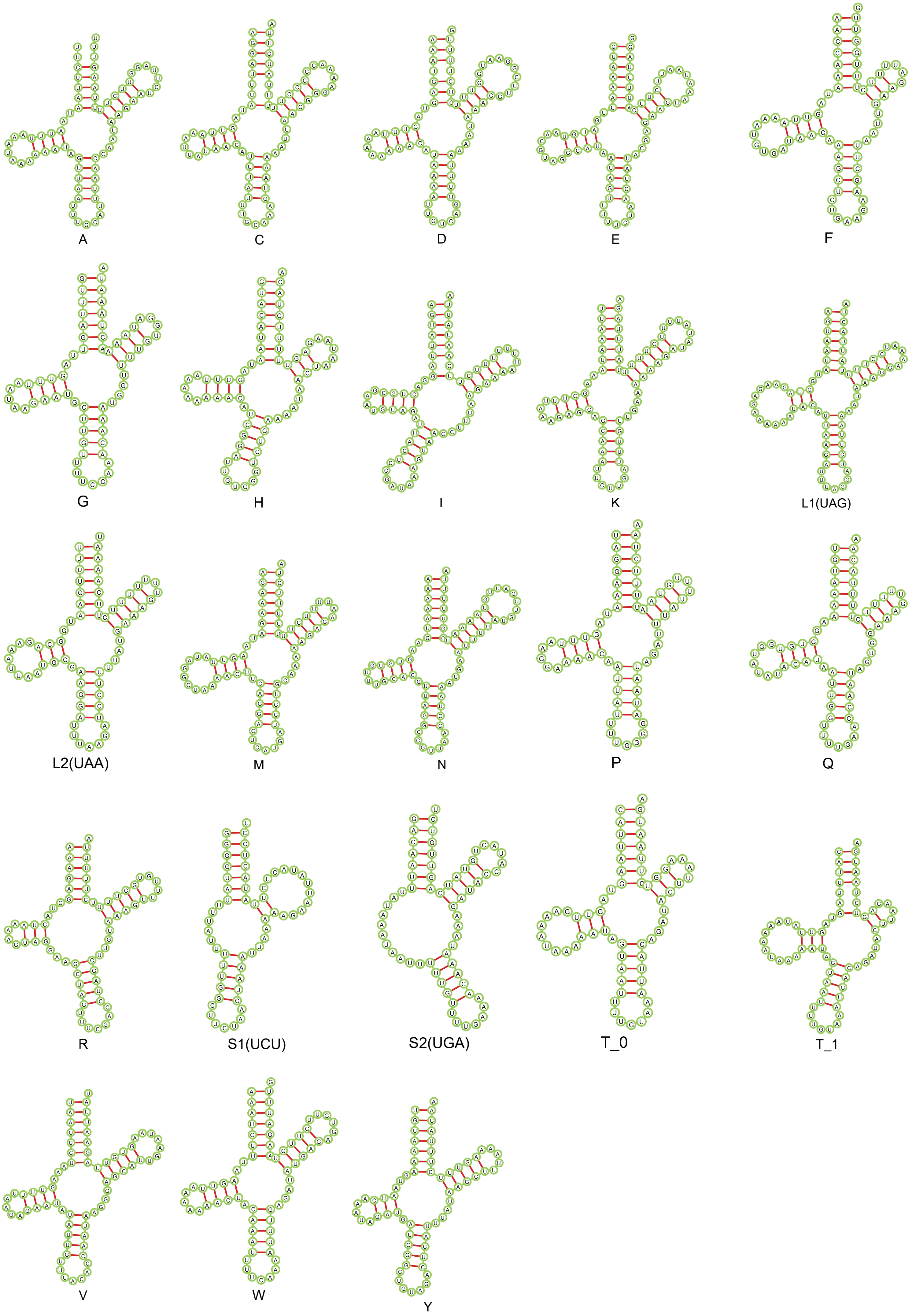

### Fig. S5 all_NCR_mafft_detail.pdf

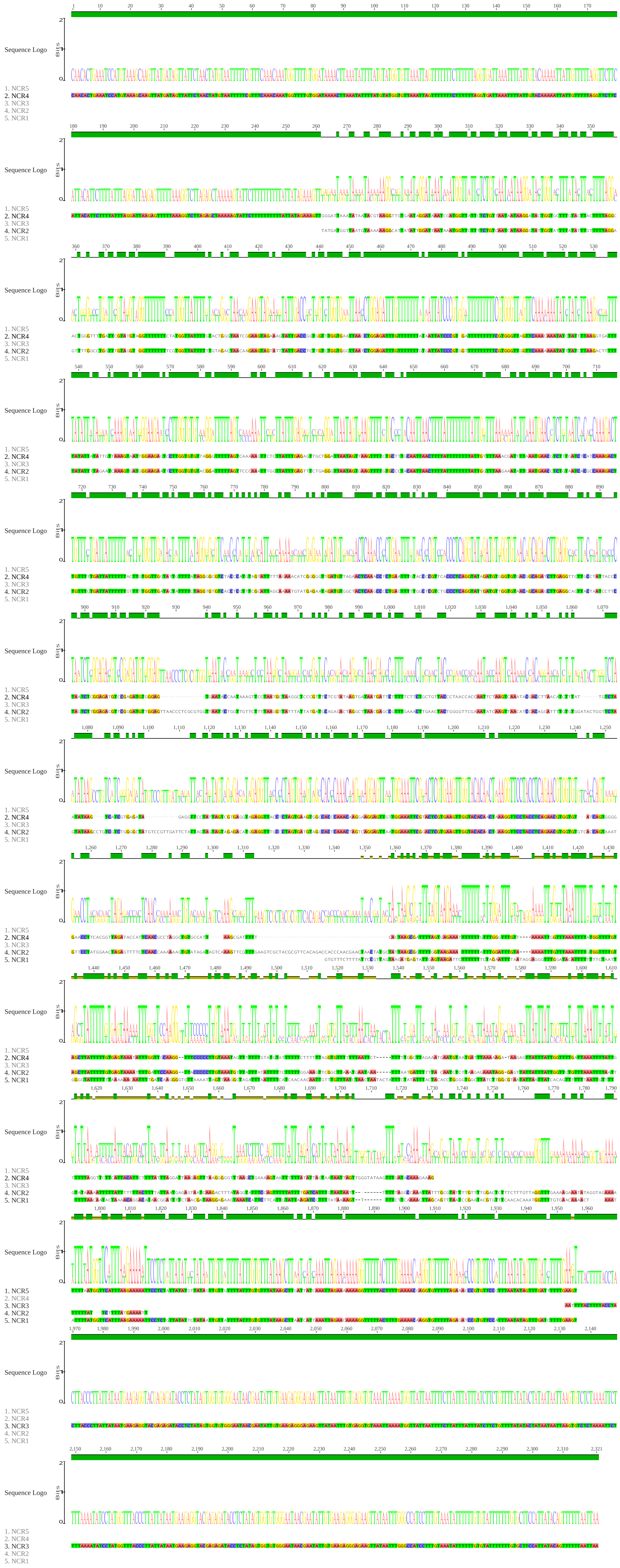

### Fig. S6 all_NCR_overview.pdf

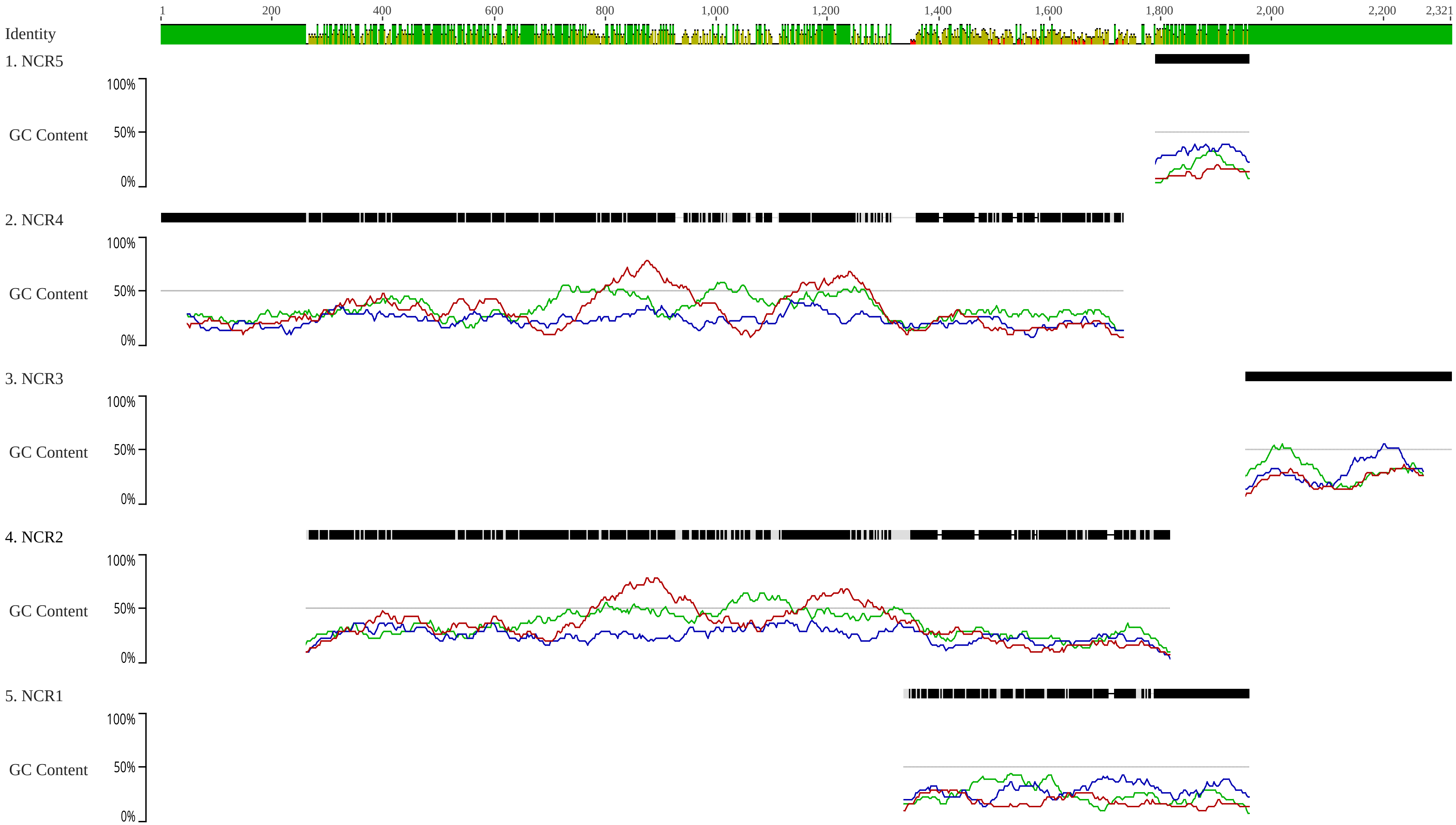

### Fig. S8 hydrophobicity patterns of the putative atp8.pdf

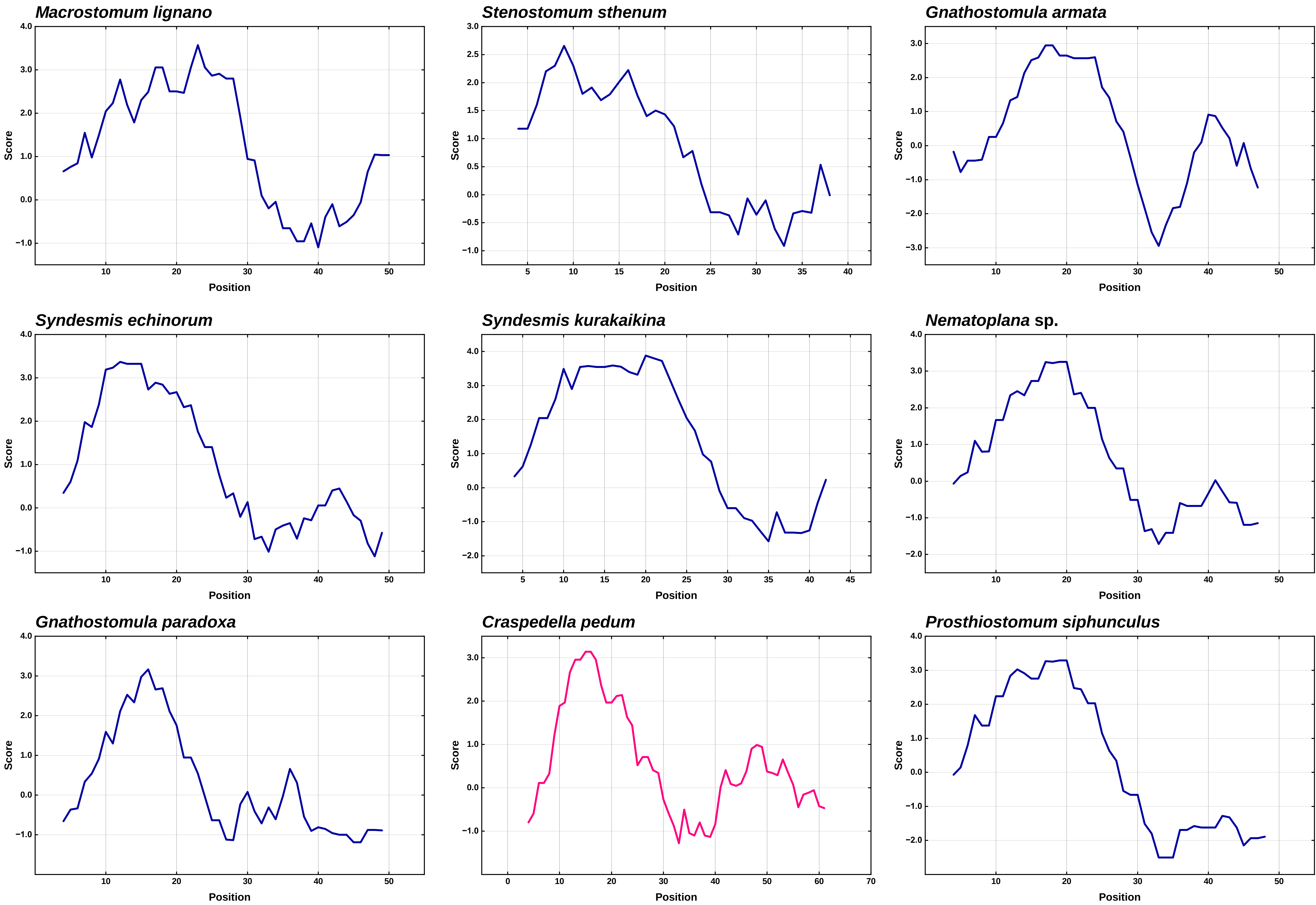

### Fig. S9 possibility of a signal peptide of cox1_frag.pdf

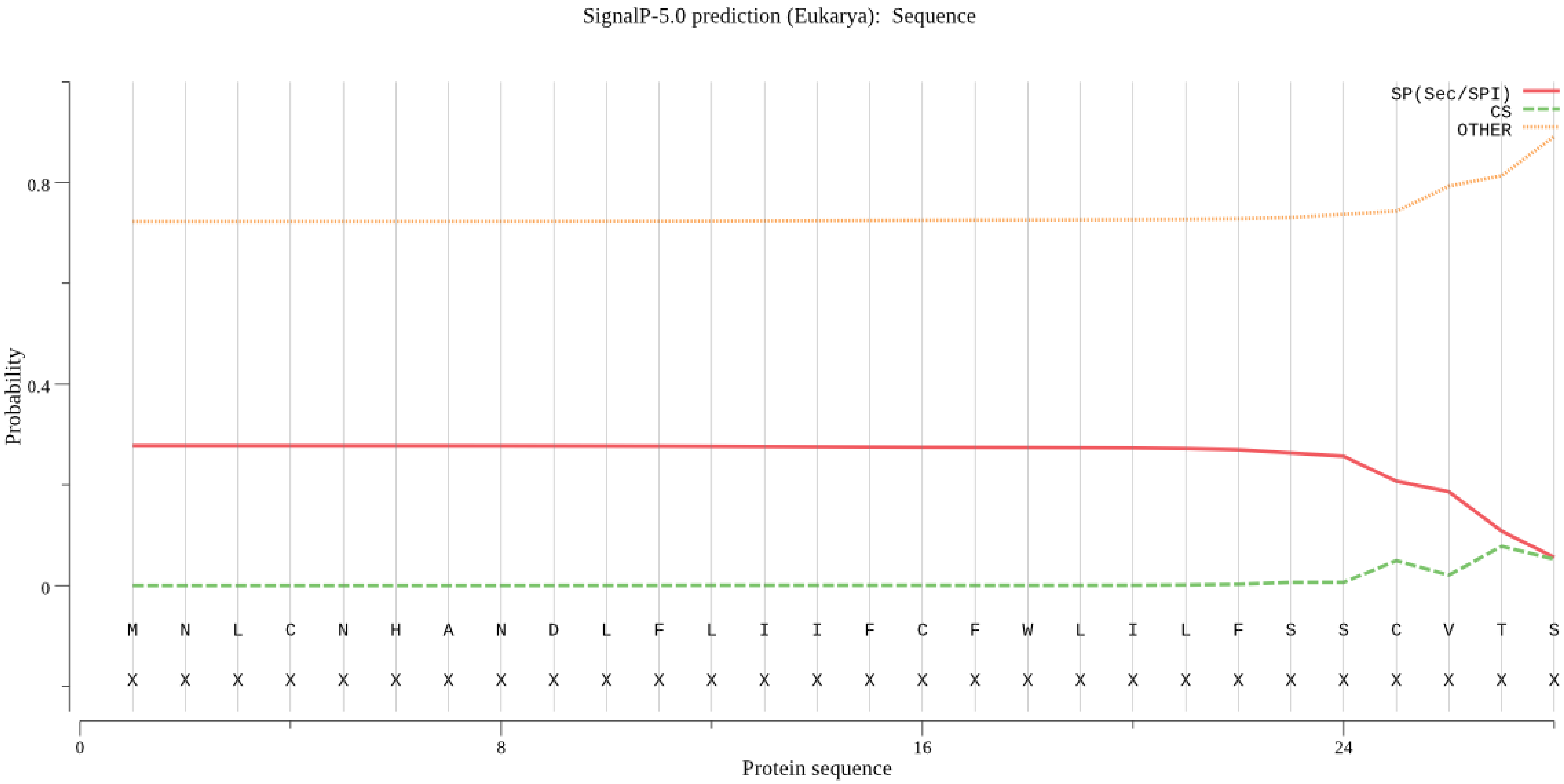

### Fig. S10 all_overlap_mafft.pdf

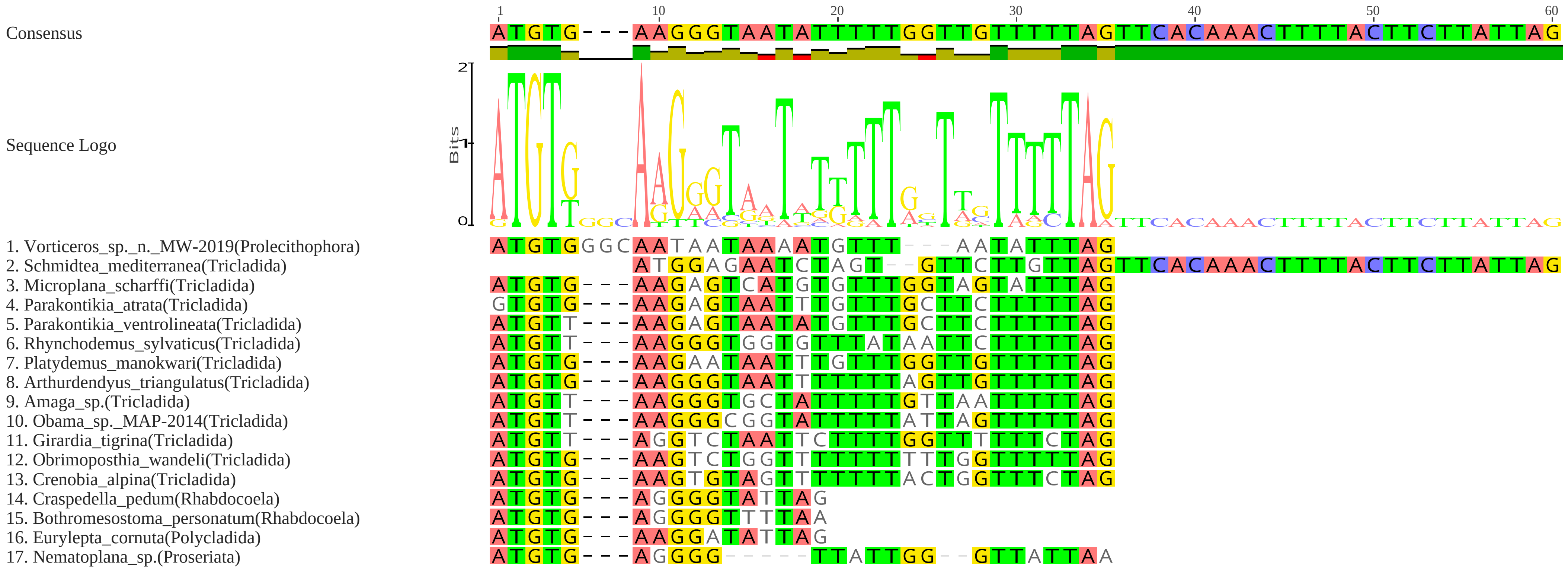

### Fig. S11 AA-BI.pdf

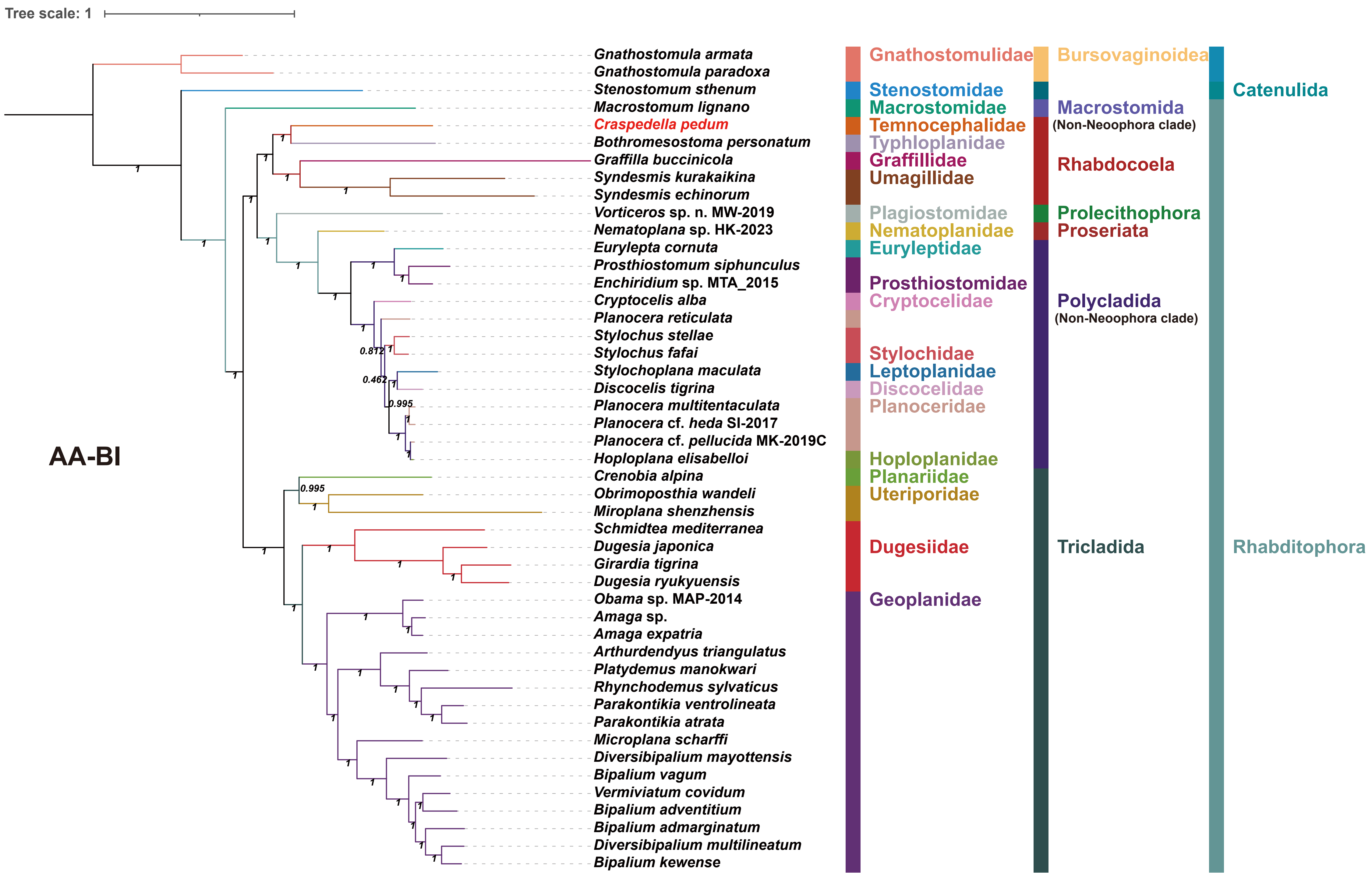

### Fig. S12 AA-IQ.pdf

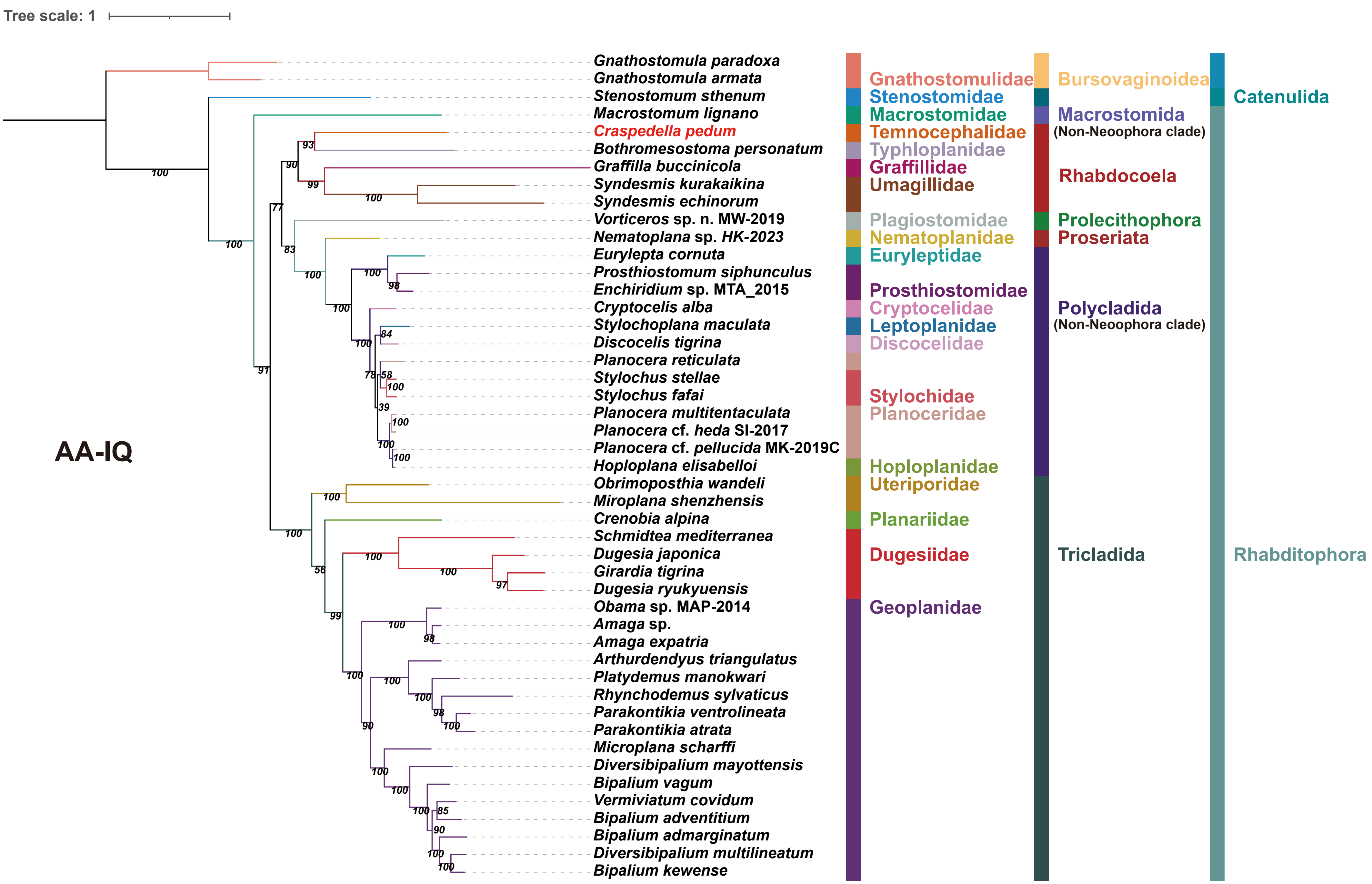

### Fig. S13 CAT-GTR.pdf

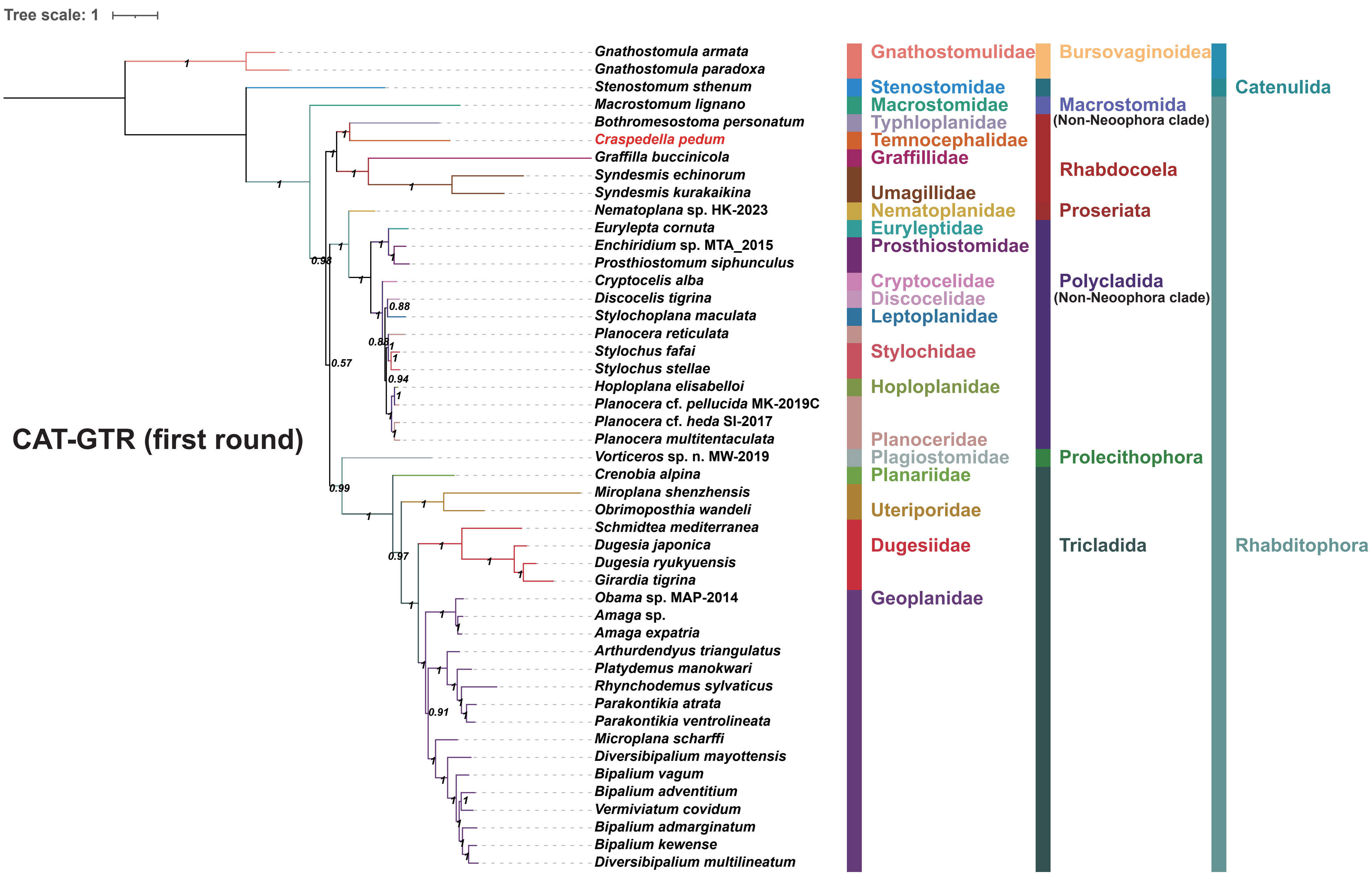

### Fig. S14 PCGsRNA-BI.pdf

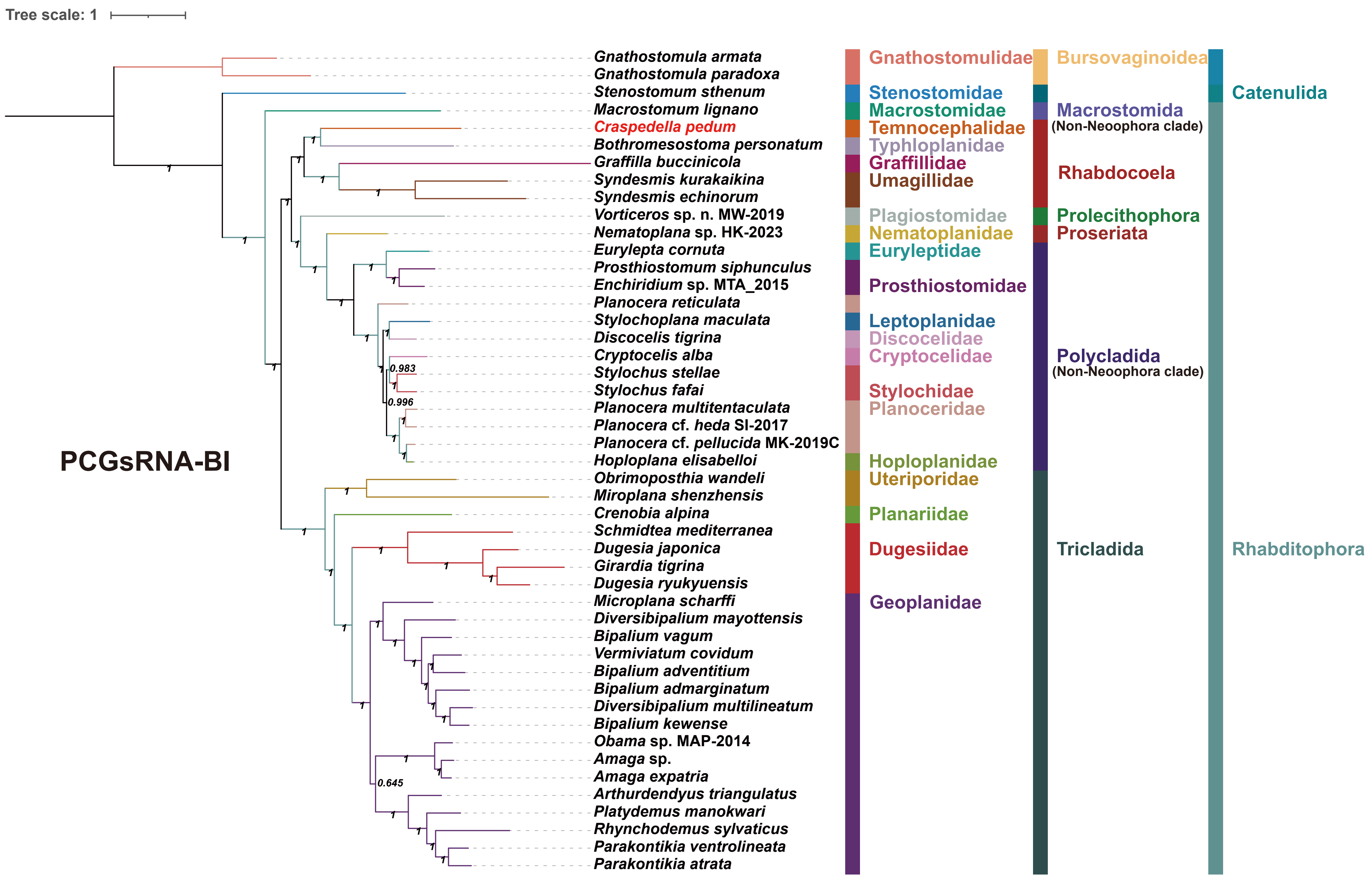
